# A three-node Turing gene circuit forms periodic spatial patterns in bacteria

**DOI:** 10.1101/2023.10.19.563112

**Authors:** J Tica, M Oliver Huidobro, T Zhu, GKA Wachter, RH Pazuki, E Tonello, H Siebert, MPH Stumpf, RG Endres, M Isalan

## Abstract

Turing patterns^1^ are well-known self-organising systems that can form spots, stripes, or labyrinths. They represent a major theory of patterning in tissue organisation, due to their remarkable similarity to some natural patterns, such as skin pigmentation in zebrafish^2^, digit spacing^3,4^, and many others. The involvement of Turing patterns in biology has been debated because of their stringent fine-tuning requirements, where patterns only occur within a small subset of parameters^5,6^. This has complicated the engineering of a synthetic gene circuit for Turing patterns from first principles, even though natural genetic Turing networks have been successfully identified^4,7^. Here, we engineered a synthetic genetic reaction-diffusion system where three nodes interact according to a non-classical Turing network with improved parametric robustness^6^. The system was optimised in *E. coli* and reproducibly generated stationary, periodic, concentric stripe patterns in growing colonies. The patterns were successfully reproduced with a partial differential equation model, in a parameter regime obtained by fitting to experimental data. Our synthetic Turing system can contribute to novel nanotechnologies, such as patterned biomaterial deposition^8,9^, and provide insights into developmental patterning programs^10^.

## Main

The engineering of spatial reaction-diffusion (RD) systems has been a major focus of synthetic biology^11–20^. This has improved our understanding of synthetic systems, helped us explain developmental patterning phenomena, and will potentially contribute to tissue engineering and other novel technologies^8,9^.

In the 70 years since the mathematician Alan Turing first proposed his famous reaction-diffusion patterns^1^, biologists and theorists alike have tried to understand how these emergent, self-organising patterns might form from deceptively simple interaction networks. The minimal reaction topology for Turing patterns involves only two diffusers: a slow-diffusing activator that activates its own production, and a fast-diffusing inhibitor that feeds back to inhibit the production of the activator^1,21^. Despite this simplicity, this model can produce rich spatiotemporal behaviour from travelling waves, to oscillations and stable regular patterns^1,22^.

From an engineering standpoint, the main problem has been that the classical 2-component Turing circuits are not very robust: even small changes in the reaction parameters destroy the patterning^5,6,23^. Turing patterns were nonetheless observed in chemical reactions, such as the CIMA and TuIS systems^24,25^. Biological systems, for example those involved in the embryonic development of bone^4^, tooth^26^ and hair formation patterns^7^, are thought to stabilise the patterning with larger, more complex dynamical networks^4^, chemical gradients of positional information^3^, and growth^27^. Noise has also been shown to relax the tuning requirements^28^, enabling engineering of stochastic Turing patterns in *E. coli*^15^, that appear as noisy patches of gene expression on a lawn of cells. However, how to accomplish regular, well-ordered patterns in an artificial system remained unknown.

To begin to address this, we previously identified circuits with improved parametric robustness to Turing patterns, with a high-throughput *in silico* study that assessed over 7,000 two- and three-node circuits with distinct topologies, resulting in 10^11^ distinct parameter and model combinations^6^. While pattern formation was commonly observed in more than 60% of the networks, very few parameter combinations (<0.1%) were found to support patterning, even in the most robust circuits^6^. Here, we sought to engineer the resulting best-performing variant in *E. coli*, with fully synthetic components, based on networks #3954 and #1754^6^.

The circuit is built with six genes, and activation and inhibition functions are assigned according to the circuit topology. With the help of a mechanistic, six-equation partial differential equation model, and close integration between model and experiments, we identified circuit tuning conditions that are particularly conducive for patterning. We then found an experimental condition where stationary stripes form sequentially on the growing edge of a colony. We used the model to fit the parameters to experimental circuit dose-response data, which yielded realistic spatial simulations in growing colonies. We thus found a local Turing condition that reproduces the experimental result well, in terms of outer ring addition and an anti-phase pattern. Overall, this study engineers a self-organising reaction-diffusion system in *E. coli* and puts forward model-based evidence in support of a Turing mechanism.

### Implementation of the circuit in *E. coli*

The patterning circuit was designed according to the three-node topology that forms Turing patterns with the largest number of parameter combinations in a computational search (Fig. 1a, inset)^6^. The topology consists of a classical self-activation and lateral inhibition motif between nodes A and B, where the positive feedback on node A is instead implemented with repressible constitutive activity^6,29^. An additional non-diffusible node C is used to increase the number of parameters that produce Turing patterns^6^, and to relax the differential diffusion requirement^30,31^.

**Figure 1:**
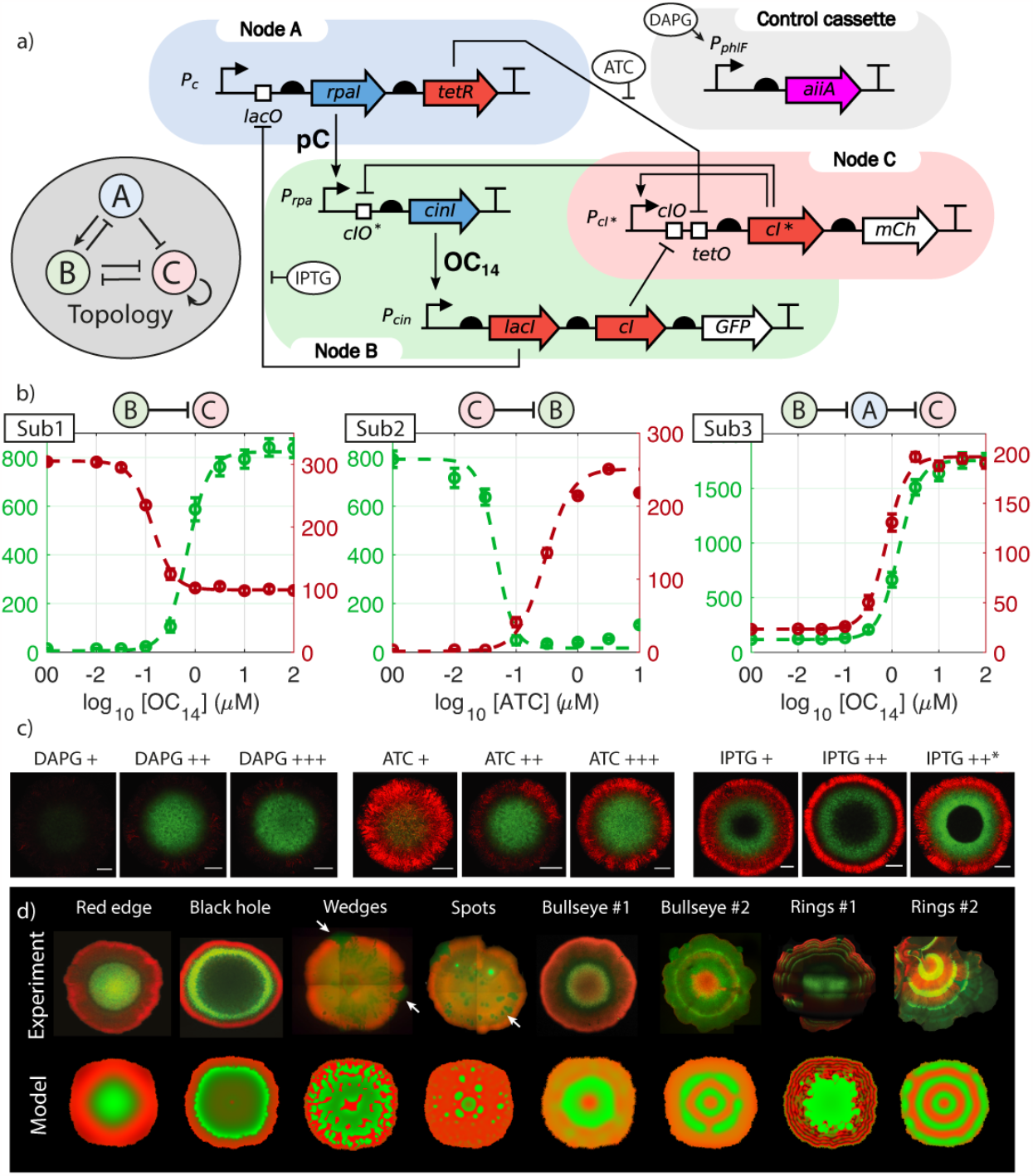
one tunable genetic circuit produces many different spatial patterns. (a) Gene circuit architecture in standard notation: diffuser synthesis enzymes are in blue, non-diffusible transcription factors in red, fluorescent proteins in white. The circuit produces diffusers OC14 and pC, and can be regulated with small molecules ATC, IPTG and DAPG as shown. The circuit topology of network #1754 is shown in the grey inset. (b) Circuit characterisation: liquid culture fluorescence 18 hours after induction of three subcircuits: subcircuit #1 (Sub1), #2 (Sub2) and #3 (Sub3). GFP on left and mCherry on right axis (unit AU/A_600_, mean ± SEM, n = 3). The tested interactions are shown above the respective plots, further information in Suppl. Info. 1. (c) Full circuit tested in growing colonies on agar, 24 hours after induction with DAPG (+++ 3.2 μM; ++ 1 μM; + 0.32 μM), ATC (+++ 0.1 μM; ++ 0.032 μM; + 0.01 μM) and IPTG (++ 10 μM; + 1 μM; ++* 10 μM IPTG & 0.01 μM ATC). Image acquisition settings are constant for each inducer, for ease of comparison. Scale bars: 200 μm. (d) Various spatial patterns are observed when the full circuit is tested in growing colonies in different experimental conditions (upper row). The white arrows show two wedges and a region of spot formation. Colony size and tuning conditions are listed in Table S1. Patterns are reproduced with the circuit model (bottom row); for parameters see Suppl. Info. 4

Nodes A and B were implemented with orthogonal quorum sensing molecules pC-HSL (pC) and 3OHC_14_HSL (OC_14_), respectively (Fig. 1a)^32,33^. Node C was implemented with an orthogonal transcription factor cIλ variant, cIλ-5C6GP, with dual activation-repression dynamics^34^. Transcription factors LacI, TetR and cI were used to implement the repressive interactions. A DAPG-inducible^32^ AiiA lactonase was used for tuneable diffuser degradation. The circuit could also be tuned with ATC and IPTG to adjust the repression levels by TetR and LacI, respectively. The pattern reporters GFP and mCherry measured the activity of nodes B and C, respectively.

The experimental system was first tested in liquid culture in a series of control experiments, to validate its function and optimise the genetic components where needed (Fig. 1b, Suppl. Info. 1). Subcircuit #1 included the Pcin cassette of node B, a constitutive TetR, together with node C. In the presence of high ATC to fully disinhibit TetR, the gradual induction of Pcin with exogenous OC14 led to an induction of node B GFP, and to a repression of node C mCherry, as expected. Subcircuit #2 tested node C, in combination with node B with a cI deletion, and a constitutive TetR. Disinhibiting TetR with ATC gradually induces node C and mCherry, and represses node B and GFP, as expected. Subcircuit #3 included the Pcin cassette of node B with a cI deletion, together with node A and node C. Activating Pcin with exogenous OC14 gradually induces node C in a double-inhibitory interaction through node A, as expected. The diffusible interactions (Fig. S7), crosstalk between the components (Fig. S8) and AiiA-dependent degradation (Fig. S9) were also tested. Taken together these results show that the circuit components function according to our design.

The system was then tested in growing colonies in a variety of tuning conditions. The colonies generally showed a green centre and a red edge (Fig. 1c). The fluorescence intensity of the green centre could be tuned with DAPG and ATC inducers, where both DAPG and ATC visibly increased green fluorescence intensity. The spatial profile of fluorescence was visibly altered by IPTG, which led to a sharpening of the red outer ring at high concentrations and caused the appearance of a dark non-fluorescent region in the colony centre at the imaged timepoint. Overall, the circuit responded to the tuning molecules when grown as colonies.

We investigated conditions under which bacterial colonies would form more complex patterns. We identified a particularly interesting regime with 10 μM ATC, where smaller colonies showed a repeating red-green-red pattern, the beginning of a periodic pattern (Fig. 1d; Bullseye #1). This is consistent with the model, which predicts that a high ATC concentration is beneficial for patterning (Fig. S18). We sought to grow larger colonies to determine whether more repeats would form. This was achieved by carefully diluting the sample and plating a single colony without any neighbours. We indeed observed periodic green and red stripes forming on the growing edge of the colony (Fig. 1d; Rings #1, #2).

In summary, the circuit was successfully engineered and produced complex patterns in colonies of growing *E. coli*. We next ventured to capture this behaviour with a computational model.

### Model fitting to experimental data

A six-equation partial differential equation model was used to simulate the concentrations of the Turing circuit species in time and space (Suppl. Info. 3, 4, 5). A nondimensionalised version of the model was studied, which expresses species’ concentrations on a relative fold-change scale, and where all parameters are expressed in a form that is convenient for model analysis and fitting.

We explored the circuit’s ability to form patterns within a biologically relevant parameter space at steady state using linear stability analysis and Latin hypercube sampling. A linear stability analysis of the circuit model in non-growing domains^35^ confirmed that stationary Turing patterns occur for 0.022% of the tested parameter combinations (Fig. 2a). Other interesting types of RD patterning were found, including Hopf in 4% of cases, which produce travelling waves, and Turing-Hopf in 0.004% of cases, which also produce regular stationary patterns (Fig. 2a)^36–38^. This linear stability analysis does not consider that growing domains and noise could further increase the Turing patterning percentage (Fig. S26)^27,28^. Modelling in discrete systems has also revealed that pattern robustness might be greater than previously thought^29,39^. Moreover, the local parameter space around Turing conditions is more highly enriched with patterns. For example, adding a relative uncertainty of 1% around a Turing I solution produces 33% Turing I solutions, whereas an uncertainty of 5% produces 5% Turing I solutions (Fig. S24). The average relative uncertainty in the *V*_*m*_ and *K*_*m*_ parameters between biological repeats in liquid culture data of Fig. 1b was 4.8%. This indicates that if a patterning region is found, then Turing patterns are sufficiently common to be reproduced.

**Figure 2:**
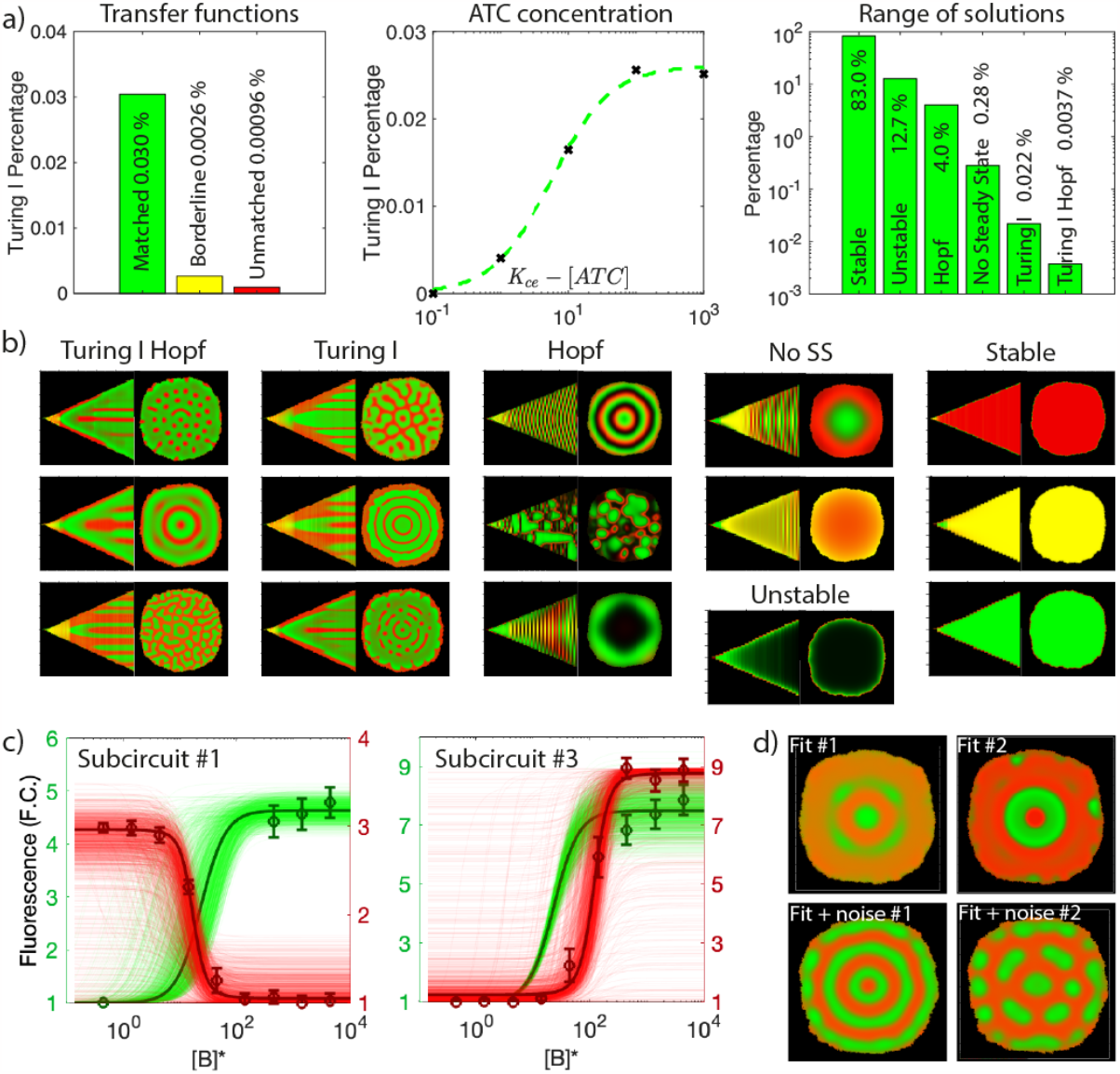
modelling genetic circuit with growth. (a) Turing pattern robustness optimisation and analytical solutions of the gene circuit model. Robustness can be increased by matching the transfer functions of the circuit components (green) compared to unmatched (red) (Left) or by increasing ATC concentration (Centre). The optimised parameters produce a wide variety of analytical solutions, expressed as percentages (Right). (b) Simulations of the hybrid PDE-bacterial colony solver for all the types of analytical solutions of network #1754. Kymograph (left) showing the timeseries of the cross-section and the final snapshot of the simulation (right). Green and red channels are superposed. (c) Fitted OC_14_ dose-response curves produced using multivariate analysis optimisation, for subcircuit #1 (left) and subcircuit #3 (right). Dots show experimental data, the thick line is generated from best fit parameters and the thin lines are derived from multivariate Gaussian distributions cantered around the best fit (Suppl. Info. 4). This distribution has a probability density function p(x; k, 10· Ck)where k is the best fit parameter vector and *C*_*k*_ is the covariance matrix (d) Turing solutions in simulated bacterial colonies using the parameters obtained after dose-response curve fitting (top) and when the fitted parameters were added a small amount of noise (bottom). Refer to Suppl. Info. 3 and 4 for details.

Exploration of the model’s parameter space guided our tuning of the circuit, to increase the probability to achieve a patterning regime. Turing pattern probability is dramatically increased (30-fold) in sub-sections of the parameter space where the transfer functions of components are ‘matched’ (Fig. 2a, Suppl. Info. 3.8). A transfer function is ‘matched’ when the gene expression levels are compatible with the sensitivity of the regulatory components they act on (Fig. S17). For example, a transfer function is not ‘matched’ when a very sensitive operator site is completely repressed even with background levels of transcription factor e.g., if these are sufficiently high because of a leaky promoter or strong RBS. In this case, an induction of the transcription factor above background would not result in further repression, leading to a loss of dynamic range. The transfer functions of the circuit components were matched by tuning plasmid copy number, the strength of ribosome binding sites (RBSs), start codons and degradation tags^40–42^. Transfer function matching yielded a well-functioning circuit (Fig. 1b) with a better capacity to produce spatial patterns.

Next, we used the model to explore the effects of the tuning molecules ATC, IPTG and DAPG. High ATC concentrations dramatically improved Turing patterning probability (Fig. 2a, centre), while DAPG and IPTG did not. These observations are consistent with the experiments, where spatial patterns were detected with high ATC but not with DAPG or IPTG (Fig. 1d, S16).

The system was then simulated numerically in growing colonies. Colony growth was implemented with a stochastic cellular automata algorithm^43,44^, where the spatial points containing cells were modelled to undergo both reaction and diffusion, whereas the points not containing cells were only modelled with diffusion, omitting the reaction component. A variety of patterns were discovered, including stationary rings, travelling waves, stationary and non-stationary spots, labyrinths and bistability wedges (Fig. 1d, 2b). This global model analysis, with biologically relevant parameters, showed that our circuit can produce a broad range of spatial patterns.

The model’s parameters were fitted to liquid culture data (Fig. 1b) with the aim of reproducing the patterns observed in the experiments. The nondimensionalised model allowed for a simple and intuitive parametrisation (Suppl. Info. 3). A multivariate analysis approach^45^ was employed to fit to subcircuit #1 and #3 data (Fig. 2c, 1b, Suppl. Info. 4). Turing patterns were found in the multivariate Gaussian distributions obtained by fitting parameters (Fig. 2d). We identified a local solution whose appearance was consistent with the experiments, where stationary, concentric rings formed with growth (Fig. 2d, Fit + Noise #1). Taken together, these findings suggest that the stationary patterns are not inconsistent with a Turing mechanism.

### Stripe formation in growing colonies

Stripe formation was captured with timelapse confocal microscopy, where three stripes formed with a constant wavelength, consistent with a regular Turing pattern, in parts of the colony with the most prominent outgrowth; two stripes formed in parts where growth was slower, and only a single stripe formed where growth was slowest (Fig. 3a-d, S14; Video S1). The number of stripes was determined by the field size, thus maintaining a constant pattern wavelength, a characteristic of Turing systems^4,46^. The green and red fluorescence are out-of-phase, as predicted by our circuit model (Fig. 3d). The stripes are added sequentially at the growing edge of the colony and are unlikely to be a result of temporal circuit oscillations.

**Figure 3:**
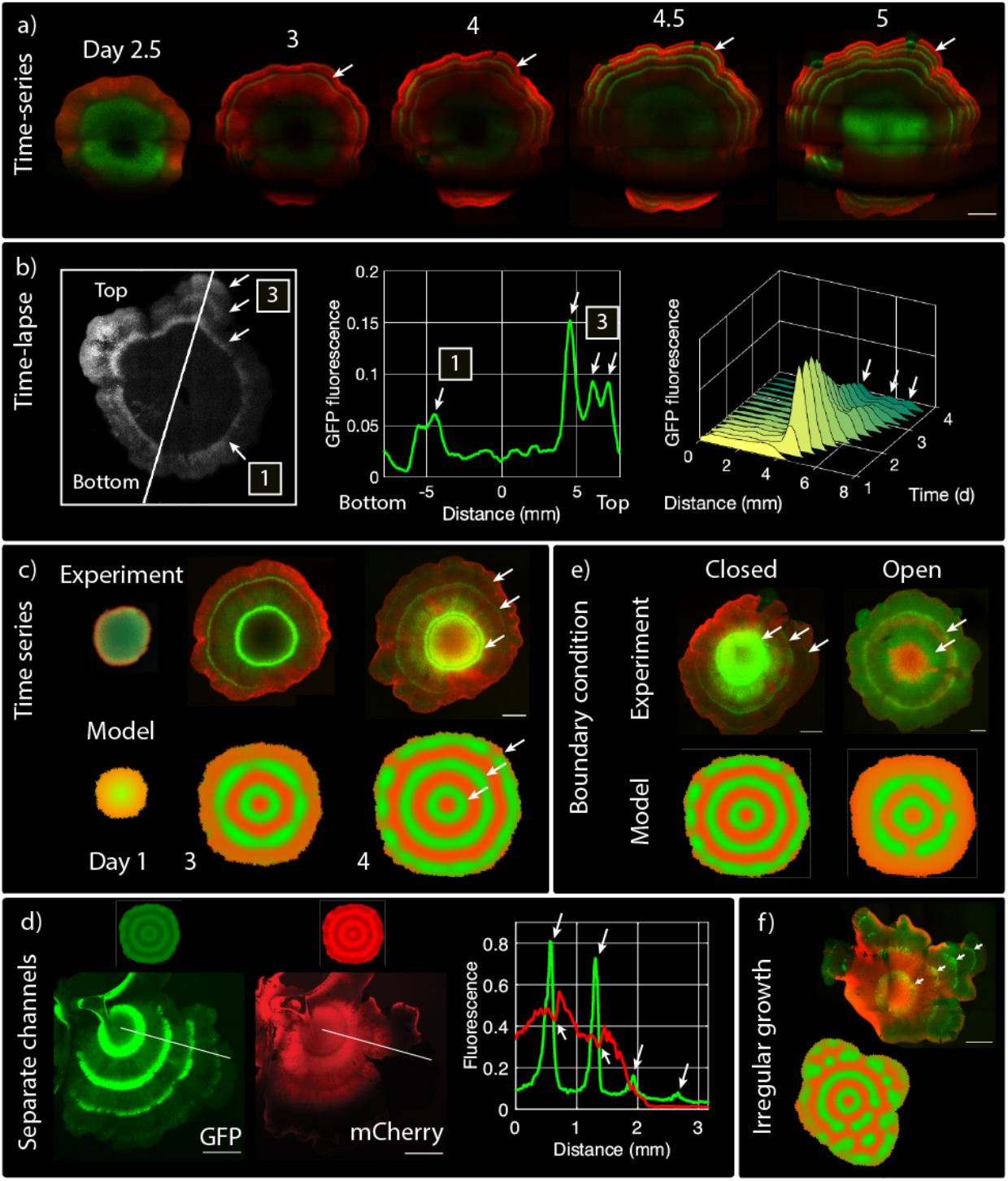
pattern formation in growing colonies. (a) Time-series of stripe formation over 5 days at 37 °C; GFP and mCherry channels are merged. Stripes form at the edge of the colony. (b) GFP fluorescence was imaged over 60 hours at constant temperature 37 °C (Video S1). Imaging began after the colony had grown for 24 hours. (Left) The image of the final timepoint shows three stripes in the top right corner of the colony where growth is most prominent (3), only a single stripe forms at the bottom where there is less growth (1). (Centre) Plot of the fluorescence along the white slanted line in the micrograph with moving average smoothing. (Right) The spatiotemporal profile of GFP evolution shows that the stripes form at the edge of the colony. After an initial burst of fluorescence, the signal decreases over time. (c) The fluorescence of a growing colony is imaged over 4 days. The model reproduces the concentric rings of the experiment. (d) A colony grown on a thinner layer of agar shows pronounced patterns. The green and red channels are shown separately. Anti-phase concentric rings are visible. The model simulations are shown above and reproduce the anti-phase rings. Fluorescence with moving average smoothing is plotted along the white axes in the micrographs, peaks (GFP) and troughs (mCherry) are shown with white arrows. (e) The boundary condition affects the patterns. Multiple rings form when cells are grown in smaller wells (closed boundary), whereas fewer rings form when grown in larger dishes (open boundary). (f) Irregular colony growth generates discontinuous concentric rings, consistent with a Turing model as indicated by white arrows. Scale bars: 1 mm.

Other types of structures were also observed, such as colony “wedges” that form on the growing edge of the colony, with visibly different fluorescence expression levels (Fig. 1d; Wedges). These are likely caused by the inheritance, expansion and competition of subpopulations occupying different circuit states^35–37^. Mutations of the circuit were not recorded in the complete sequencing of its plasmids, in the different regions of the colony, a week after the start of the experiment (Suppl. Info. 6).

The pattern generator circuit produced different outputs under different experimental conditions. For example, the pattern was eliminated by adding DAPG to increase diffuser degradation (Fig. S16). Similarly, the pattern did not occur at 30°C (Fig. S16), likely due to the dependence of kinetic rate parameters on temperature^47^. Taken together, these results suggest that the patterns are unlikely to be formed by a mechanism independent of the circuit.

Additionally, reducing the thickness of the agar gave much brighter patterns (Fig. 3d, S15), consistent with Turing models where diffuser dilution in the underlying agar negatively impacts pattern formation^48^. The model predicted a disruption of stripe formation when using a boundary condition permeable to the diffusers at the edges of the spatial domain; this was achieved experimentally by growing the colonies in larger dishes (Fig. 3e). Lastly, the model also reproduced stripe formation with irregular colony growth (Fig. 3f). Collectively, these results show that the pattern dynamics are consistent with an RD model of a Turing pattern.

## Discussion

Our synthetic gene circuit, implemented according to a more robust Turing circuit topology, led to self-organising, stationary, periodic stripes in growing colonies. The model reproduced the out-of-phase stationary stripe pattern with a Turing solution, obtained by fitting the model parameters to the experimental data. Taken together, our results show that the best condition for stationary concentric rings are when colonies are grown from single cells on a thin layer of 2xYT agar, with a thickness of around 1 mm, supplemented with a high ATC concentration of 10 μM, plated in circular wells with a diameter of 32 mm, incubated at 37 °C. Engineering such patterns can contribute to novel nanotechnologies for example by linking patterns to biomaterial deposition^8,9^, and give insights into developmental biology by showing that Turing patterns can be produced by genetic systems^10^.

Consistent with the mathematical model, our circuit exhibits a variety of spatial patterns. The circuit exhibited varied dynamics in growing colonies, where different behaviours were often observed within the same tuning condition, and even within the same colony. For example, the colony in Fig. 3a exhibited stationary stripes on the edge together with oscillations restricted to the green centre, which fluoresces at the 2.5 day and 5 day timepoints but not in between. Another colony exhibited stationary stripes on the edge together with a propagating wave of mCherry fluorescence in the north-western section of the colony (Fig. 3b, S14, Video S1). The circuit model can generate all these dynamical behaviours in the immediate vicinity of the best-fit Turing condition (Fig. S24). The stationary stripes are consistent with a Turing solution, whereas the propagating waves with a Hopf solution.

Other models that can generate stationary ring patterns include the clock and wavefront model^49^, where a growing domain interacts with a cellular oscillator to deposit a repeating pattern. In synthetic biology, this behaviour was observed in growing colonies bearing an oscillating circuit^50,51^. However, in our study there is no evidence of oscillations at the growing edge of the colony, so this explanation is unlikely. Alternatively, certain irregular patterns can be explained by solitary localised structures, which was previously proposed for an engineered mammalian patterning system^19^. Both theories would struggle to describe the observed behaviours, including regular stationary stripes, which are captured well by our model.

While the stripe patterns were readily reproducible in multiple independent experiments, performed on different days, the patterns and colony dynamics in each experiment were unique. This is likely because of small differences in colony growth, epigenetics, concentrations of the inducers, thickness of the agar, and all the other experimental variables. Natural systems can also be variable. For example, the circuits for zebrafish skin produce a slightly different pattern each time^2^, whereas the digit formation networks of up to 7 nodes are pattern invariant^4^. Future work can explore pattern variability in more depth, to see if larger topologies and positional information can explain these differences^4,5^. The self-organising nature of Turing systems has long fascinated scientists and, for the first time, these complex systems are ripe to be explored with the tools of synthetic biology.

## METHODS

### Cells and media

*E. coli* 5α (NEB) were used for cloning, whereas MK01 cells^52^ were used for all the experiments. 2xYT media was the primary outgrowth medium for fluorescence assays and microscopy (Sigma-Aldrich Y1003), SOC was used for cell recovery after transformation.

### Electroporation of *E. coli*

Cells were grown in LB media, washed with MilliQ water, resuspended in 10% glycerol to make them competent. Cells were electroporated with a pulse of 1.8 kV, 200 Ω, 50 μF, recovered in 500 μL SOC broth shaking for 1 hour and 37°C, plated on LB agar with antibiotics and incubated overnight at 37°C. Cells were first transformed with the pCC1 plasmids, made electrocompetent using the protocol above, and then transformed with the other circuit plasmids.

### Molecular cloning

Gene-synthesised DNA fragments were amplified with Q5 Hot-Start Polymerase (NEB), DpnI digested at 37°C for 1 hour, extracted with the Monarch Gel Extraction Kit after running in 1% agarose at 90 V for 1 hour, assembled with the HiFi Assembly Kit for two or more fragments, or with the KLD kit for simple mutagenesis or deletions. Assembled DNA was transformed to NEB 5α chemically competent cells, colonies were screened with colony PCR after overnight incubation, correctly assembled clones were grown up in 8 mL 2xYT, purified with the Qiagen QIAspin MiniPrep kit and sequenced with the Eurofins SupremeRun Sanger sequencing service. Plasmid sequences are listed in Suppl. Info. 7 and plasmids have been deposited on Addgene (TBC: Ref. numbers).

### Whole circuit sequencing

At the end of microscopy experiments, different parts of the colonies were picked and grown in 2xYT medium overnight at 37°C. Cells were resuspended in 20% (v/v) glycerol and frozen to -80°C. The glycerol stocks were grown up in 8 mL 2xYT medium, the plasmids were purified with the QIAprep MiniPrep kit, and sent for NanoPore sequencing.

### Liquid culture fluorescence assays

Electroporated MK01 cells were grown to mid-exponential phase (OD_600_ 0.4 – 0.6) shaking at 37°C, transferred to a 96-well plate and mixed with the relevant inducer(s) dissolved in 2xYT medium to a final volume of 150 μL. The plates were transferred to the Tecan SPARK plate reader, readings of absorbance at 600 nm and fluorescence were collected over 18 hours. Settings for GFP: ex. 495/25 nm, em. 525/25 nm, gain 25. Settings for mCherry: ex. 625/25 nm, em. 625/25 nm, gain 50, Z-position 17806 μm. Shaking at 150 rpm, 2 mm.

Data analysis was performed in MATLAB R2022a. The final fluorescence values were background-subtracted with 2xYT-only readings. Data were normalised to absorbance where mentioned. Dose-response data was fitted with Hill functions (Eq. 1). The background expression (α) and maximal regulation (*V*_*m*_) parameters were calculated directly from the data by taking the smallest and largest values of fluorescence, respectively. The *Km* was fitted using the in-built least-squares fitting routine *lsqcurvefit*. The best fitting cooperativity parameter *n* was selected after manually testing various integer values.

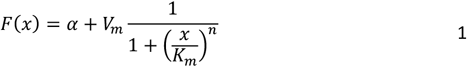

Percentage variability in the fitted parameters was calculated by fitting Hill functions with the routine above to each biological repeat individually. The relative deviation from the average was then calculated for *V*_*m*_ and *K*_*m*_ parameters for each of the three subcircuits (Fig. 1b). The reported value is an average across the *V*_*m*_ and *K*_*m*_ parameters for all three subcircuits.

### Confocal microscopy of pattern formation

Cells were grown to mid-exponential phase (OD_600_ 0.4 – 0.6) shaking at 37°C, dissolved in 2xYT by a factor of 1:10.000 and plated in serial 1:5 dilutions to 6-well plates, with 1 mL 2xYT agar containing the relevant antibiotics and inducers on the bottom of each well. The plates were incubated at 37°C, sealed with parafilm and wrapped in aluminium foil. Microscopy was performed with the Leica Stellaris 5 confocal microscope.

Image acquisition settings for the Stellaris 5: objective HC PL APO CS2 10x/0.40; camera Leica DFC9000 GTC; excitation with white light laser (WLL) at 489 nm for GFP and 587 nm for mCherry, laser power 85%, relative laser intensity between 5% and 20%; emission with HyD detectors at 494-499 nm for GFP and 592-660 nm for mCherry, detector gain between 5% and 50%; fluorifier disc NF 488/561; pinhole 2 AU (106 μm). Relative laser intensity and detector gain were adjusted to suit the individual images, except where it is explicitly mentioned that they were kept equal across a series of images.

Time-lapse experiments were performed in 35 mm μ-dishes (ibidi 81156) with the Leica Stellaris 5 at 37°C, using the adaptive focus features to identify the focal plane at each timepoint, images were acquired every hour. Temperature conditions were maintained with the Okolab H101-BASIC-BL with an Okolab stage top chamber. Images were prepared and analysed with Fiji and MATLAB R2022a.

### Widefield microscopy of pattern formation

Cells were grown on 1.4% LB-Agar at 37°C overnight. The colonies were then spotted in the centre of a standard 12 well tissue culture plate using a sterile toothpick with the relevant antibiotics and inducers. Different agar thicknesses and media types were tested. The plates were incubated at 37°C, sealed with parafilm and wrapped in aluminium foil. Microscopy was performed with a Carl Zeiss Cell Observer microscope. 5x objective: EC PlnN Ph1 DICO, NA 0.16. 10x objective: EC PlnN Ph1 DICI, NA 0.3. GFP fluorescence: ex. 470/40 nm, em. 525/50 nm, filter set Zeiss 38. mCherry fluorescence: ex. 545/25 nm, em. 605/70 nm, filter set Zeiss 43. Fluorescence light source: HXP 120V, intensity 40%. Image acquisition: 4x4 binning, automatic exposure time. Images were prepared and analysed with Fiji.

### Mathematical model

To reproduce the results from the reaction-diffusion system in bacterial colonies, a partial differential equation (PDE) solver was developed which includes colony growth, shape, and boundary conditions.

The genetic circuit is described with a PDE system that models the concentrations of the two diffusible species pC-HSL and 3OHC_14_-HSL and intracellular transcription factors TetR, LacI, cI and cI5G6GP in time and space. The circuit receptors are modelled with constant concentration, consistent with their constitutive expression. Quasi-steady state is assumed for the mRNA production step as well as for diffuser production. Each equation includes a basal production term due to promoter leakiness, a regulated production Hill term, a linear degradation term and a diffusion term. The model was studied in a non-dimensional form, where most of the variables and parameters are expressed as ratios and have dimensionless units. The non-dimensional model is easier to parameterise with data from experiments by comparing the model’s output to fluorescence data, as both concentration and fluorescence have dimensionless units, and are expressed as a fold change from the basal steady state for each species. For details see Suppl. Info. 3.

The dynamics of the colony are modelled using a cellular automaton^43^ which is a discrete model of computation that can accurately describe how the bacterial colony evolves over time. The cellular automaton consists of a 2-dimensional grid of cells which can be in an *on* or *off* state. If an *on* cell has any *off* neighbours, it will divide into the neighbouring *off* cell with a probability p_d_. Over time, a stochastic circular *on* domain will be generated, which resembles a bacterial colony. In this case, *on* states correspond to *E. coli*, while *off* states correspond to agar. No cell death (*on* to *off* transition) is permitted. This cellular automaton is integrated with the PDE system by computing reaction and diffusion terms in the *on* cells and only diffusion terms in the *off* cells. Newborn cells inherit the full concentration of their mother cells. Dilution is neglected in this algorithm, but accounted for in the degradation terms of the PDE. This hybrid algorithm captures the genetic circuit and colony growth. For details see Suppl. Info. 5.

The PDE is then solved using the alternating-direction implicit method (ADI). The outer square boundaries can have reflecting boundary conditions (Neumann boundary condition where derivative at the boundary is zero) or absorbing boundary conditions (Dirichlet boundary condition where solution at the edge is zero) depending on the experimental system. For a small agar plate, we define reflecting boundaries while for a larger plate we assume absorbing boundary conditions.

## ACKNOWLEDGEMENTS

We thank Natalie Scholes, Alexandre Djehizian, Haobin Chen, Lucy Joy Cornell and Nina Bahun for contributing to the foundations of this work. This research was supported by the Volkswagen Foundation (LIFE: 93 065).

## AUTHOR CONTRIBUTIONS

M.I. and J.T. conceived the research and designed the circuits. J.T. and T.Z. performed most of the experiments. G.K.A.W. performed widefield microscopy experiments. M.O.H. performed the simulations, linear stability analysis, model fitting, and some experiments. R.E. supervised the modelling work. J.T. and M.O.H. analysed data. R.H.P. and J.T. conceived the model non-dimensionalisation. E.T., H.S. and M.P.H.S contributed to circuit design by modelling and discussions. J.T., M.O.H., M.I. and R.E. wrote the manuscript. All authors reviewed and edited the manuscript.

