## Supplementary Materials for "A three-node Turing gene circuit forms periodic spatial patterns in bacteria"

### Supplementary 1: Circuit Characterisation

We sought to verify and characterise the behaviour of all the genetic components of our circuits and their interactions. We wanted to be sure that everything functions according to our design and is consistent with the model, before testing the circuit for patterning.

Closed feedback loops often lead to non-monotonic dose-response behaviours, which are difficult to interpret, especially when studying a new system. For example, a band-pass dose-response function can be produced by a band-pass circuit, or simply be a result of an induction at low inducer concentrations, combined with metabolic burden-dependent crashing at high inducer concentrations[1]. Hence, it was useful to develop subcircuits with no closed feedback loops to simplify the interpretation of the data, to identify bugs and fix them, to move towards the desired circuit behaviour.

Five control experiments were sufficient to test all the circuit's genetic components and interactions. All the experiments were performed in liquid culture and were also repeated in growing colonies on agar. All the small molecule receptors are expressed from constitutive promoters on a low copy plasmid pCC1 (Fig. S**1**).

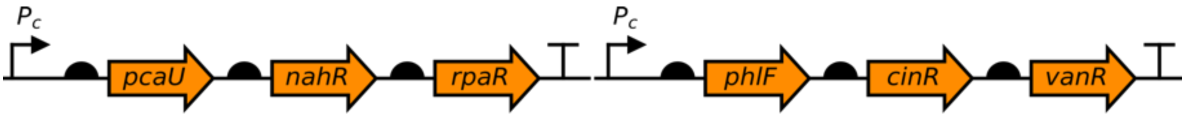

**Supplementary Figure 1: control cassette with constitutively expressed regulators.** The regulators that are used in the system are: RpaR is the pC receptor (Node A diffuser), CinR is the OC14 receptor (Node B diffuser), and PhlF is the DAPG receptor (used to tune inducible diffuser degradation). The other three receptors PcaU, NahR and VanR are for the protocatechuate, salicylate and vanillin inducers, which are not used in the present circuit, but were tested in other versions of the system. These three systems can be used to introduce additional regulatory components to the system.

The five control experiments presented here consist of: (1) three subcircuits of the full system, containing no closed feedback loops; (2) a system to test the inter-cellular, diffusive interactions between circuit elements, achieved by OC14 and pC; (3) crosstalk between the three inducible elements of the circuit, OC14-Pcin, pC-Prpa and DAPG-PphlF; (4) DAPG-inducible degradation of OC14 and pC by AiiA lactonase.

All experiments described in this section are performed in 2xYT liquid culture, shaking continuously at 37 °C, measurements are collected 18 hours after induction. Results are plotted on a logarithmic x-axis, where the no inducer controls are added manually under the '00' datapoint.

More information about the system plasmids and the sequences are in Supplementary 7:

##### 1.1. Subcircuit #1: repression of Node C

The first subcircuit tests the inhibitory Node C inputs from TetR (from node A) and cI (from node B). The system consists of node C, the Pcin cassette of node B, and a variant of node A with a constitutive promoter lacking lacO (Lac operator). The latter is used to avoid Node B-dependent repression of Node A, which would complicate the interpretation of the results.

This subcircuit was optimised to show good expression in the absence of cI and TetR repression. The induction of Pcin in Node B with OC14, or the removal of ATC, gradually repressed Node C, as expected. The dose-response relationship to OC14 shows a negative correlation between GFP and mCherry, as expected; both responses occur with a good dynamic range; no burden[1,2] is observed at either end of the OC14 and ATC concentration ranges (Fig. S**2**).

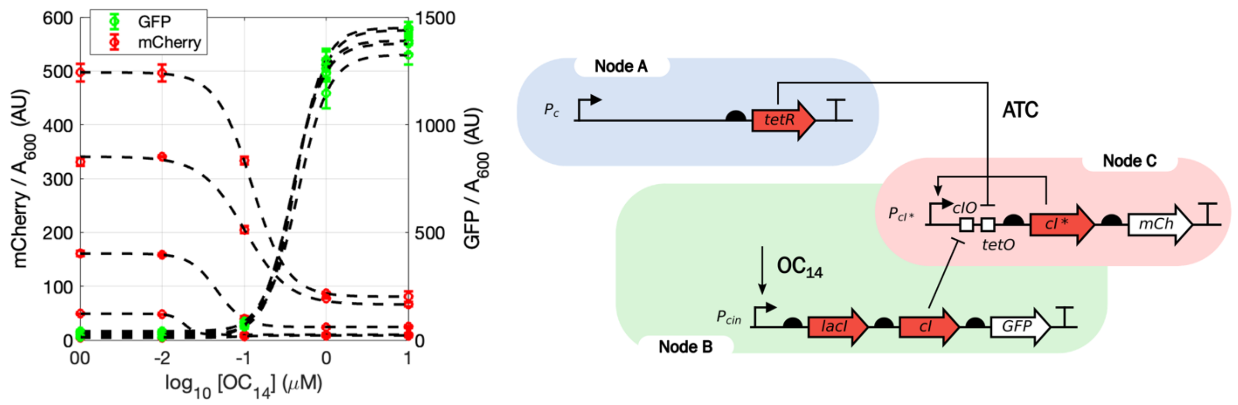

**Supplementary Figure 2: design and induction of subcircuit #1.** The subcircuit was designed to test the behaviour of the node C positive feedback and its repressive TetR and cI inputs. An increase in $OC_{14}$, and a reduction in ATC induce mCherry through cI and tetR, respectively. A negative relationship between mCherry and GFP is observed, as expected. The ATC concentrations are $10^{0.5}$, $10^{0}$, $10^{-0.5}$, $10^{-1}$ and 0 μM, where the higher mCherry expression levels correspond to the lower ATC concentrations. Arbitrary units of fluorescence, mean ± SEM, n = 1 (2 technical replicates).

##### 1.2. Subcircuit #2: Node C repression of Node B

The second subcircuit tests the inhibitory Node C – Node B interaction. The expression assays were performed in the presence of a constant concentration pC for Prpa induction. Node C was then gradually derepressed with ATC, which was expected to inhibit node B activity and GFP expression.

Node B is active and expresses GFP in the absence of ATC, when Node C is repressed by TetR. As TetR is disinhibited with increasing concentrations of ATC, Node C and mCherry expression gradually turn on and inhibit Node B and GFP. Consistent with expectations, the dose-response relationship of ATC shows a negative correlation between GFP and mCherry; no burden[1,2] is observed at either end of the ATC concentration range (Fig. S**3**).

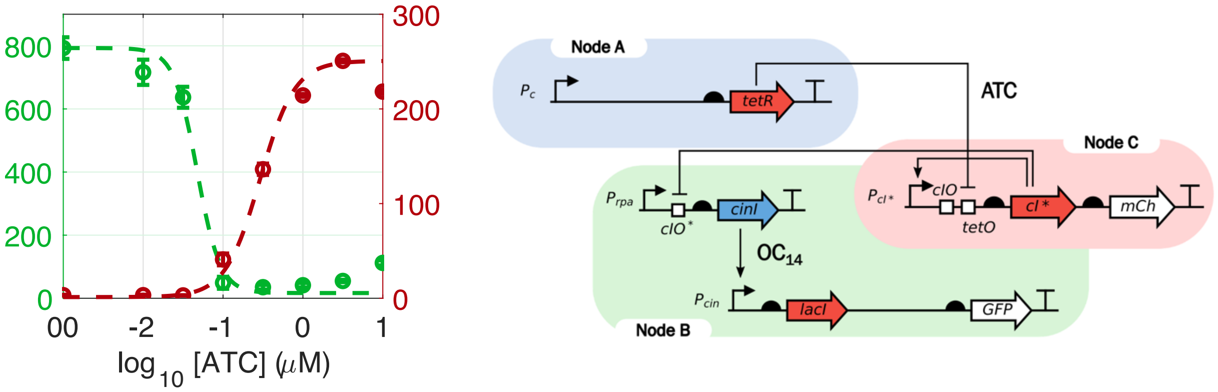

**Supplementary Figure 3: design and induction of subcircuit #2.** The subcircuit tests the behaviour of the Node C – node B repressive interaction. In the presence of background pC for Prpa activation, an increase in ATC induces mCherry through tetR, and represses GFP through cI*. A negative relationship between mCherry and GFP is observed according to expectations. Arbitrary units of fluorescence, mean ± SEM, n = 3.

##### 1.3. Subcircuit #2b: Node B activation

In this variant of the subcircuit #2, Node B is tested and optimised in isolation. Promoter Pcin is induced by endogenous OC14. The feedforward Prpa – Pcin motif was initially showing high levels of GFP even in the absence of any Prpa induction (data not shown). This was due to the amplification produced by CinI, where each unit of enzyme produces many $OC_{14}$ molecules at a high efficiency. Even the background levels of Prpa activity were sufficient to strongly induce the downstream Pcin.

To overcome this problem and achieve a good dynamic range of GFP expression, where there is low expression in the absence of pC-dependent induction of Prpa, the expression of CinI was weakened by using a weak RBS (Biobrick BBa_B0033) and weakened start codons GTG and TTG[3,4] (Fig. S**4**). The weak RBS variant with the ATG start codon was used in the final circuit.

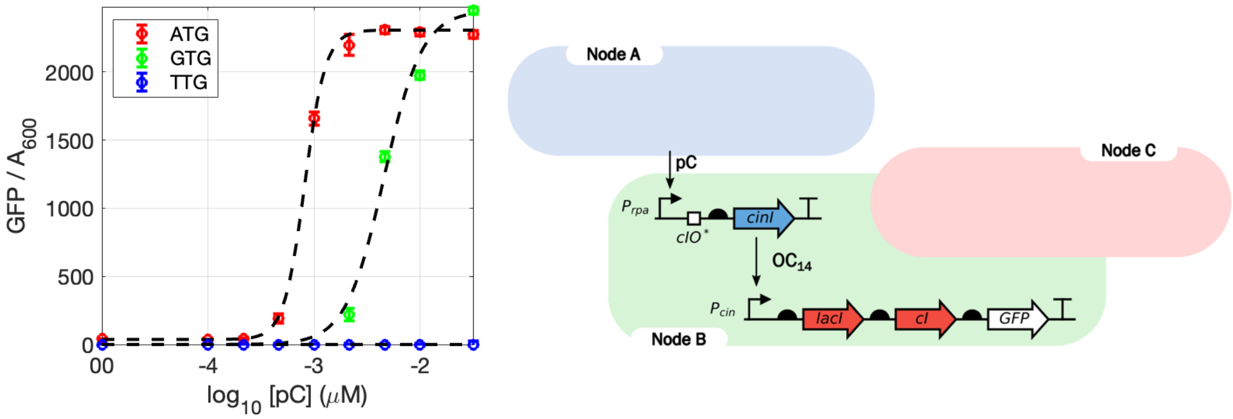

**Supplementary Figure 4: testing the node B feedforward motif.** This OC14 synthesis gene CinI is an amplifier that easily overloads the downstream Pcin promoter. CinI was therefore expressed with a weakened RBS to achieve low background expression levels and an inactive Pcin promoter in the absence of pC. The amplification in the cascade is demonstrated by the very steep dose-response curve, where even a small amount of induction highly induces the downstream Pcin system. Arbitrary units of fluorescence, mean ± SEM, n = 1 (3 technical replicates).

##### 1.4. Subcircuit #3: Node B -> Node A -> Node C axis

The third subcircuit tests the Node B -> Node A -> Node C inhibitory axis. The cI gene in Node B is removed to avoid the double repression of Node C, which would complicate the interpretation of the results; this interaction was verified as part of subcircuit #1. In subcircuit #3, exogenous OC14 should activate LacI from Pcin, LacI then represses TetR expression from Node A, leading to an induction of mCherry in Node C.

The dose-response relationship shows a positive correlation between GFP and mCherry, as expected; both responses occur with a good dynamic range (Fig. S5). No burden[1,2] is visible at high levels of OC14; therefore, the cells can sustain high GFP and high mCherry production simultaneously.

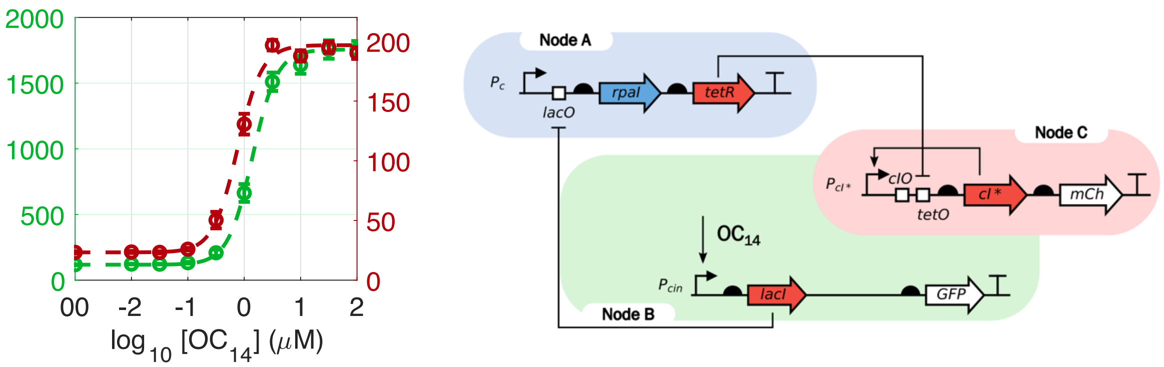

**Supplementary Figure 5: design and induction of subcircuit #3.** The subcircuit was designed to test the behaviour of Node A, and the Node B – Node A – Node C inhibitory axis. Induction with exogenous OC14 leads to a repression of Node A, and to an activation of Node C. A positive relationship between GFP and mCherry expression levels is expected and observed. Arbitrary units of fluorescence, mean ± SEM, n = 3.

##### 1.5. Subcircuit #3: tuning with ATC and IPTG

Subcircuit #3 was then tuned with the small molecules IPTG and ATC, which bind and derepress LacI and TetR, respectively. This allowed us to identify the working ranges for these tuning molecules and assess their effects on the circuit dynamics.

The effects of ATC are first seen at the high end of the OC14 concentration series where node A repression is high and TetR expression levels are low (Fig. S**6**). With maximal node A repression, ATC removes background TetR activity and increases mCherry expression levels beyond the maximum that occurs in the absence of ATC. With higher ATC concentrations this derepression also happens at low OC14 concentrations, leading to a reduction in the dynamic range of the Node A-dependent mCherry repression. Note that in the full circuit mCherry is also repressed independently by cI of Node B.

The effects of IPTG are focused on the high end of the OC14 concentration series where LacI expression is most active (Fig. S**6**). IPTG derepresses LacI, leads to greater Node A activity and increased TetR-dependent Node C repression; hence, a reduction in peak mCherry expression is observed in the presence of IPTG. There is no change at the low end of the OC14 concentration series, where LacI expression is not induced.

Together, these data confirm that the circuit can be tuned with ATC and IPTG, and that all the circuit components function according to our design.

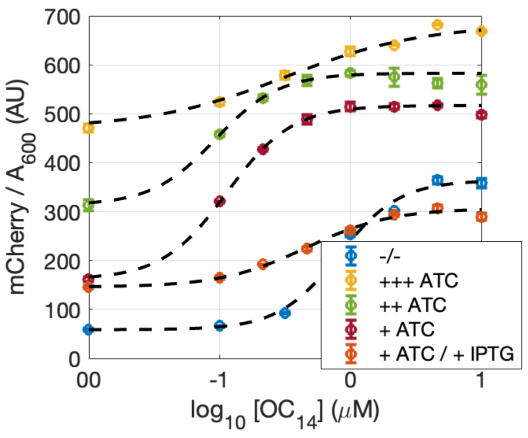

**Supplementary Figure 6: ATC and IPTG tuning curves for subcircuit #3.** The mCherry (right) expression levels are plotted against OC14 concentration in the following conditions: $10^{-0.5}$ µM ATC (+++); $10^{-1}$ µM ATC (++); $10^{-1.5}$ µM; ATC (+); $10^{-1.5}$ µM ATC with $10^{2}$ µM IPTG; in the absence of ATC and IPTG (-/-).

##### 1.6. Intercellular communication via pC and OC14

Next, the diffusive interactions in the circuit were tested, including pC signalling of Node A and OC14 signalling of Node B.

The first experiment (Fig. S7, red bars) involved co-culturing cells containing Node A with cells containing Node B. Results show a clear induction of GFP fluorescence in the co-culture compared to the Node B-only controls. This confirms the pC produced in the Node A sender cells crosses the membrane, diffuses to receiver cells and induces Node B.

The second experiment (Fig. S7, green bars) involved co-culturing cells containing the Prpa cassette of Node B with cells containing the Pcin cassette of Node B. Results show a clear induction of GFP fluorescence in the co-culture in the presence of pC. This confirms that Node B can produce the OC14 diffuser in sender cells, which diffuses to receiver cells and induces the Pcin.

The third experiment (Fig. S7, blue bars) involved co-culture three types of cells, the first containing Node A, the second containing the Prpa cassette, and the third containing the Pcin cassette. There is significant induction of GFP in the absence of any exogenous stimulation, compared to the control where Prpa cells are excluded. This shows that the circuit's endogenous activity can strongly activate the production of both diffusers and trigger inter-cellular signalling.

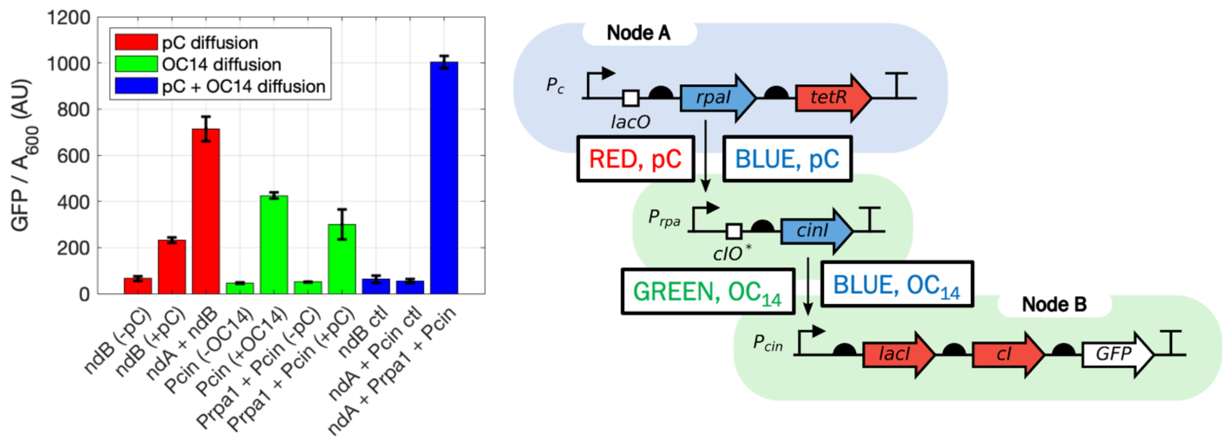

**Supplementary Figure 7: testing intercellular diffusive interactions.** Three different co-culture experiments were performed. **(Red)** Node A cells induce Node B cells. In a cell culture of only Node B cells there is a relatively small induction with exogenous pC at 1 µM (ndB -pC vs. ndB +pC). **(Green)** Pcin Node B cells can induce Prpa Node B cells in the presence of exogenous pC (1 µM) but not in its absence. A control where Pcin Node B cells are stimulated with exogenous OC14 (10 µM) also shows induction. **(Blue)** Node A cells induce Prpa Node B cells, which in turn induce Pcin Node B cells without any exogenous stimulation. Node B cells-only control (ndB ctl) and Node A cells with Pcin Node B cells control (ndA + Pcin ctl) show no induction, as expected.

##### 1.7. Crosstalk between small molecule signals

Crosstalk between the three small-molecule inducible systems, OC14-Pcin, pC-Prpa and DAPG-PphlF, was tested in liquid culture (Fig. S**8**) and in colonies on agar (data not shown). Preliminary experiments were done with two other inducible systems[5], Sal-Psal and PCA-P3B5B, but were excluded from further study due to unfavourable results in the subcircuit experiments (data not shown).

In liquid culture, each of the inducible systems expressing mCherry were activated with each of the small molecule inducers (activation crosstalk). The ability of a non-cognate inducer to repress the cognate inducer was also tested by exposing the reporters activated by their cognate inducers to each of the three non-cognate inducers (inhibition crosstalk). No crosstalk was observed between the three systems, either by activation or repression (Fig. S**8**).

To maintain the context of the full circuit the experiments were performed also in the presence of a DAPG-inducible lactonase AiiA. The inhibition of the Prpa and Pcin systems by DAPG occurred because of AiiA induction (Fig. S**8**, right).

On agar, the inducers were plated as small droplets and three colonies carrying mCherry under the control of the three inducible systems, were placed to immediately surround them. The inducers activated only their cognate receptors; no unwanted crosstalk was observed between any of the species (data not shown).

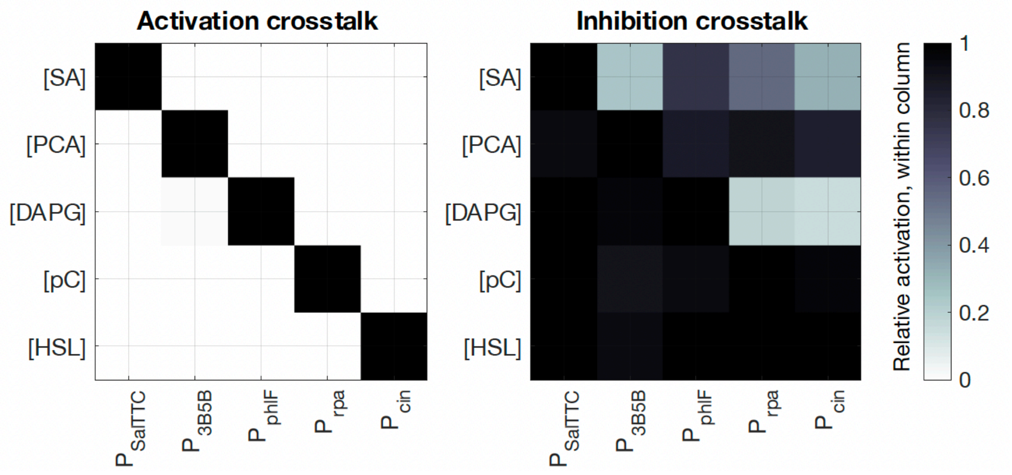

**Supplementary Figure 8: crosstalk by activation and inhibition of non-cognate systems.** Activation crosstalk is measured by inducing five promoters (columns) with five small-molecule inducers (rows). Each inducer activates only their respective promoter. Concentrations of inducers: [SA] = $10^{3.5}$, [PCA] = $10^{3}$, [DAPG] = $10^{1}$, [pC] = $10^{3}$, [OC14] = $10^{2}$ (μM). Inhibition crosstalk shows that the small-molecule inducers (excluding SA) cannot repress the response of a promoter to its cognate inducer. Concentrations of inducers to stimulate their respective promoters: [SA] = $10^{2.5}$, [PCA] = $10^{2}$, [DAPG] = $10^{0.5}$, [pC] = $10^{0}$, [3OHC14HSL] = $10^{0}$ (μM). Concentrations of inducers to stimulate non-cognate promoters: [SA] = $10^{3.5}$, [PCA] = $10^{3}$, [DAPG] = $10^{1}$, [pC] = $10^{2}$, [OC14] = $10^{1}$ (µM). The concentrations were chosen so that the non-cognate inducer concentrations are greater than the cognate inducer concentrations, to make sure that the non-cognate inducers can outcompete the cognate inducers in the event of crosstalk.

##### 1.8. Inducible degradation of pC and $OC_{14}$ by AiiA lactonase

The enzymatic degradation of the small molecule diffusers pC and OC14 was tested by expressing mCherry from their inducible promoters Prpa and Pcin, and by gradually inducing AiiA lactonase expression with increasing concentrations of DAPG. The expression of mCherry and therefore the induction of Prpa and Pcin decreases with increasing AiiA induction (Fig. S**9**). This confirms that AiiA degrades both pC and OC14.

The efficiency of AiiA degradation of pC and OC14 is similar, as indicated by their similar Ki parameters (Fig. S**9**; OC14 Ki = 1 µM, pC Ki = 0.4 µM), representing the level of AiiA induction needed for half-maximal degradation.

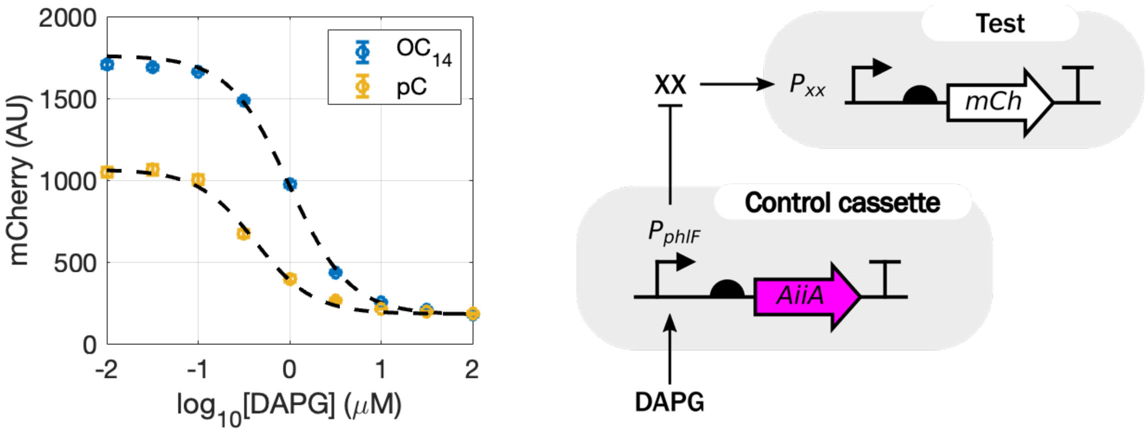

**Supplementary Figure 9: AiiA-dependent degradation of pC and OC14.** The first experiment tested degradation of pC, where XX is pC and Pxx is Prpa, corresponding to the yellow curve in the plot on the left. The second experiment tested degradation of OC14, where XX is OC14 and Pxx is Pcin, corresponding to the blue curve on the left. Both diffusers degrade with similar kinetics. The differing maximal induction levels at [DAPG] = $10^{-2}$ µM are due to the different strengths of Pcin and Prpa promoters at the given inducer concentrations; [OC14] = [pC] = 1 µM are used throughout.

##### 1.9. Growth curves of the tested subcircuits

The growth curves are shown across the induction conditions for the subcircuits #1 – #3 (Fig. S**10**). The induction of subcircuits #1 and #2 with OC14 leads to mild reduction of stationary phase absorbance. The endogenous production of OC14 in response to node B stimulation by exogenous pC in subcircuit #3 does not cause growth disturbances for any of the tested induction concentrations. The tuning of subcircuits #2 and #4 with ATC does not show dose-dependent disturbances in growth.

Overall, the growth of the circuits is relatively unaffected for the various tuning conditions. Even though there is a small reduction in absorbance for subcircuits #1 and #2 with OC14, this disappears when OC14 is produced endogenously in subcircuit #3 as will happen in the final circuit.

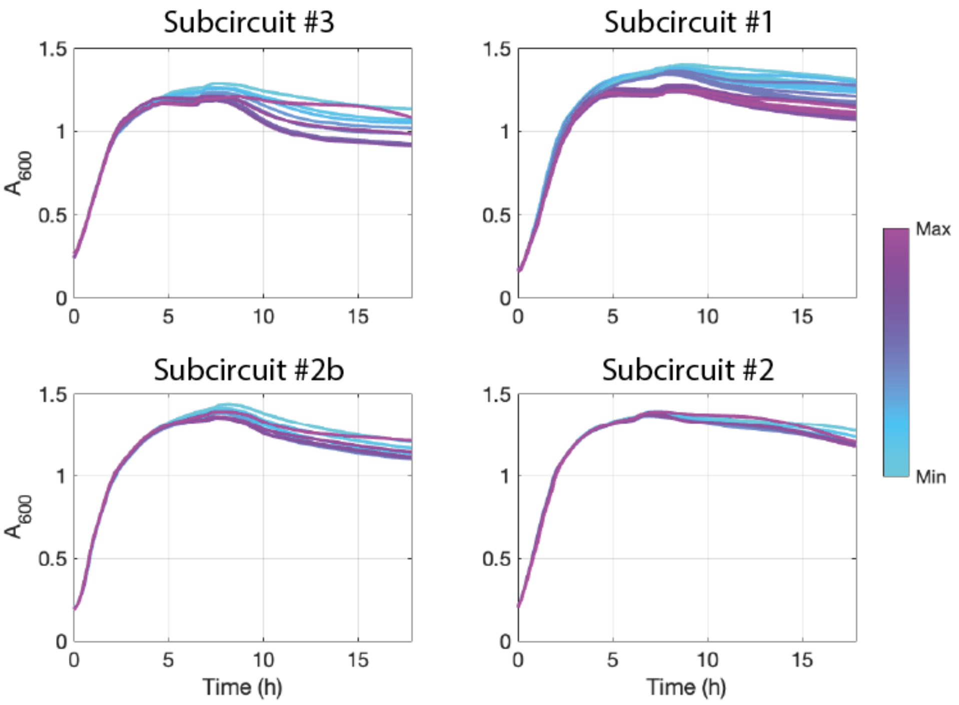

**Supplementary Figure 10: growth curves across induction conditions.** Subcircuit #3 on the top left, induction of Pcin with $OC_{14}$ series. Subcircuit #1 on the top right, induction of Pcin with $OC_{14}$ series and disinhibition of constitutive TetR with ATC series; colours change across the $OC_{14}$ series. Subcircuit #2b, only node B on the bottom left, induction of Prpa with pC series. Subcircuit #2 on the bottom right, disinhibition of constitutive TetR with ATC series in the presence of background induction of Prpa with pC. Darker shades of green represent increasing inducer concentrations.

### Supplementary 2: Spatial Patterning in Growing Colonies

After having optimised the circuit components, bacteria were grown as colonies on semi-solid agar for spatial patterning. Interesting periodic patterns were identified particularly with high [aTc] concentration (~ 10 µM). Colonies were imaged with confocal fluorescence microscopy periodically. Some of the observed stripe patterns are shown in the main paper and below. The experimental conditions for all the images are presented in Table S**1**.

**Supplementary Table 1: concentrations of inducers and colony sizes for the patterning conditions shown in the main text Figures.** Spots and wedges are shown with an asterisk (*) because they were obtained with widefield microscopy rather than confocal microscopy, in 4xYT media rather than 2xYT. Agar is 1.4 % (w/V) in all conditions. The Fig. 3a colony was first grown for 24 hours and then imaged for 60 hours; the Figure shows the colony at the final timepoint (**).

| Figure | Inducers | Timepoint (hours) | Colony Diameter (mm) |
|---|---|---|---|
| **Fig. 1d, Red Edge** | 10 µM ATC | 38 | 3.5 |
| **Fig. 1d, Black hole** | 10 µM ATC | 26 | 3.3 |
| **Fig. 1d, Wedges*** | 15 µM ATC | 100 | 10.6 |
| **Fig. 1d, Spots*** | 30 µM ATC | 100 | 6.4 |
| **Fig. 1d, Bullseye #1** | 10 µM ATC | 50 | 4.6 |
| **Fig. 1d, Bullseye #2** | 10 µM ATC | 120 | 8.6 |
| **Fig. 1d, Rings #1** | 20 µM ATC | 124 | 6.5 |
| **Fig. 1d, Rings #2**<br>**Fig. 3e** | 10 µM ATC | 120 | 4.0 |
| **Fig. 3a** | 10 µM ATC | 65, 77, 102, 110, 124 | 4.8, 5.3, 5.6, 6.4, 6.7 |
| **Fig. 3b** | 10 µM ATC | 89 | 10.5 – 13.5 |
| **Fig. 3c** | 10 µM ATC | 24, 72, 96 | 2.3, 5.7, 6.3 |
| **Fig. 3d, Closed** | 10 µM ATC | 96 | 5.4 – 6.3 |
| **Fig. 3d, Open** | 15 µM ATC | 120 | 8.8 |
| **Fig. 3f** | 10 µM ATC | 120 | 3.7 – 5.9 |

#### 2.1. Fluorescence timeseries of stripe pattern formation

This is an example where stationary, concentric stripes are formed on the edge of a growing colony (Fig. S**11**). A single snapshot of this colony is shown in Fig. 1d, Rings #1. Here the two fluorescence channels are shown separately, and show anti-phase patterns in GFP and mCherry, which is consistent with the negative regulation between nodes B and C. Tuning condition [aTc] = 10 μM, temperature of 37 °C, 2xYT agar (1.4% w/V).

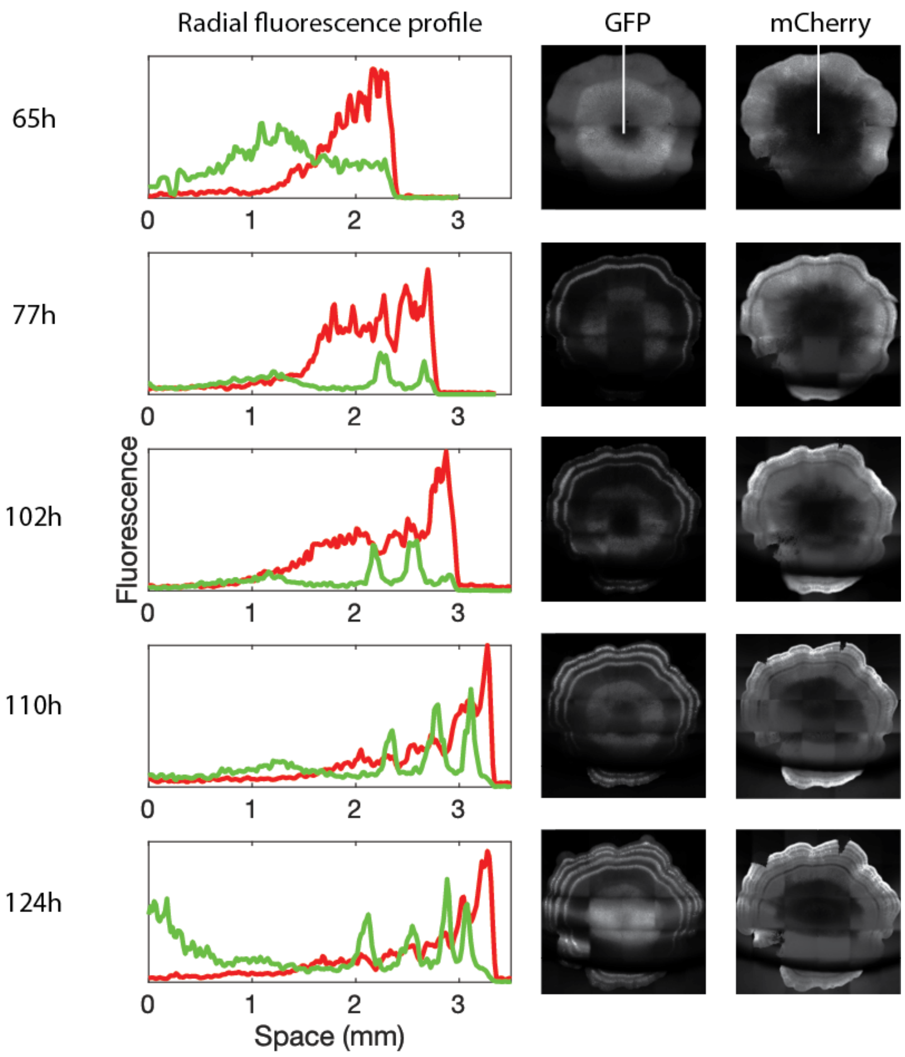

**Supplementary Figure 11: stripe formation at the growing edge of a colony.** The radial fluorescence profile is plotted along the white axis shown in the images of the 65h timepoint. Colony images not to scale between different timepoints; for scale refer to x-axis of the plots on the left.

The timeseries of the two colonies growing into each other shows that the concentric ring patterns form independently in the two colonies (Fig. S**12**).

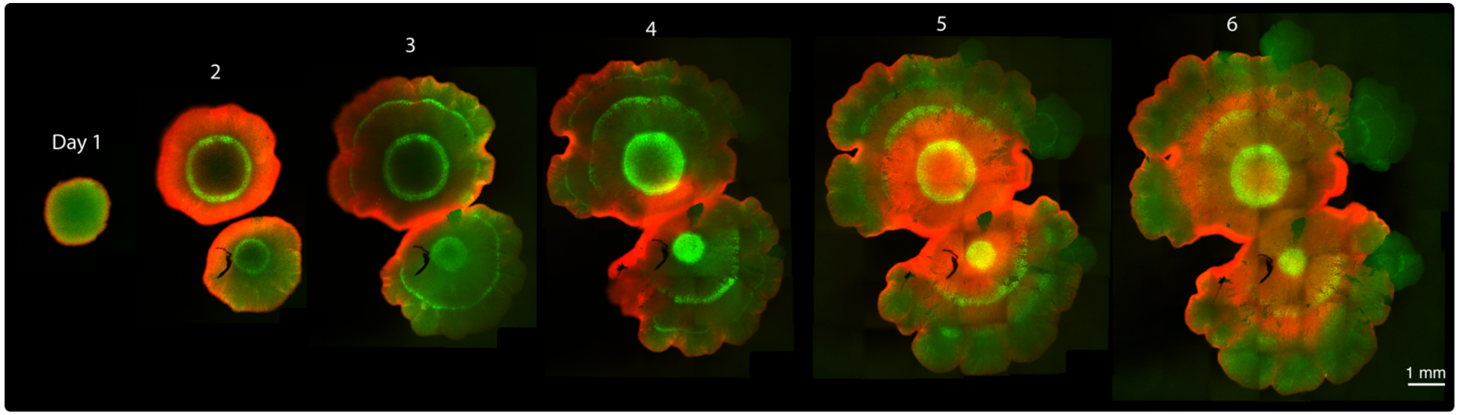

**Supplementary Figure 12: timeseries of two colonies growing into each other.** The rate of stripe formation is approximately one per day. The intensity of the stripes is more pronounced at the earlier timepoints, the pattern starts fading on and after day 4.

The timeseries of the colony showing irregular growth of Fig. 3e (Fig. S**13**).

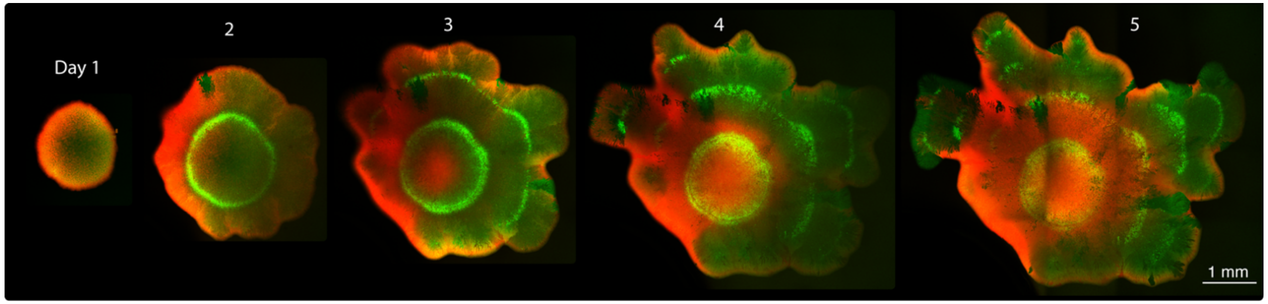

**Supplementary Figure 13: timeseries of the colony with irregular growth.** The rate of stripe formation is approximately one per day. The intensity of the stripes starts fading on and after day 4.

#### 2.2. Timelapse imaging of stripe formation and travelling waves

The colony shown in Fig. 3b is analysed though another cross-sectional axis (Fig. S**14**). On the top side of the colony (+ve) there are two stripes, highlighted with the two arrows in all the plots. On the bottom side of the colony (–ve) there is only one faint stripe. The kymographs on the bottom show the stripe formation process in time. From the kymographs notice that the GFP and mCherry stripes are clearly out-of-phase (white arrows).

The timelapse data reveals a highly dynamic spatial behaviour, coexisting with the stationary patterns. A wave of mCherry fluorescence that propagates towards the edge of the colony (Fig. S**14d**). The black dotted box and dots show the approximate location and progression of the wave. This behaviour is reminiscent of Hopf solutions of the model.

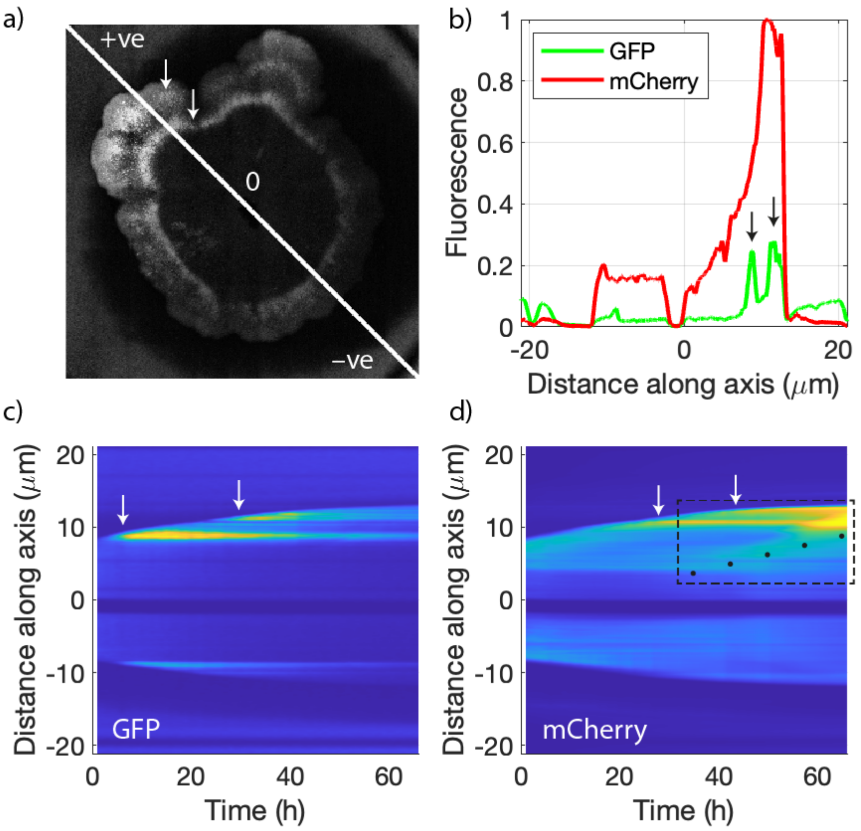

**Supplementary Figure 14: stationary stripes and propagating waves in a growing colony.** (a) Image of the colony at the final timepoint with the diagonal axis along which fluorescence is plotted in the other graphs. (b) Plot of the GFP and mCherry fluorescence along the diagonal colony axis. The two GFP stripes are highlighted with the black arrows. The peak of the mCherry signal is out-of-phase with the GFP signal. On the negative side of the colony (bottom) a single GFP stripe forms due to less growth. (c, d) Kymographs showing the spatiotemporal dynamics of GFP and mCherry expression. The points of formation of the white stripes are labelled with the white arrows. The mCherry propagating wave is shown with a dashed box and black dots.

##### 2.3. Thickness of agar layer

The thickness of underlying layer of agar on which colonies are cultured affects the intensity of the pattern (Fig. S**15**). The pattern is more intense with thinner agar, probably because the dilution of the diffusers into the underlying layer of agar negatively impacts the activity of the circuit and pattern formation. This was independently reported in computational simulations of Turing pattern where they show how pattern robustness decreases with thick agar layers[6]. This is also additional evidence that the pattern we are observing is a reaction-diffusion pattern dependent on diffusive signalling.

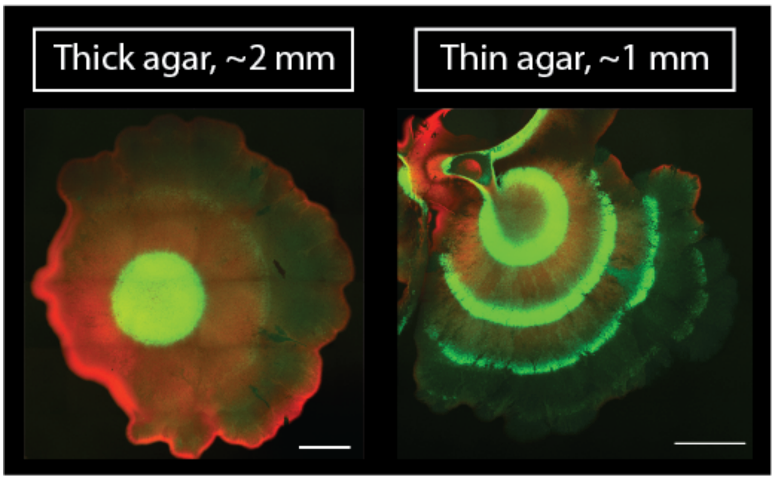

**Supplementary Figure 15: agar layer thickness affects pattern formation.** Colonies grown in the presence of 10 μM ATC. Images taken at day 5 after plating. Scale bars: 1 mm.

##### 2.4. Conditions where no green stripe patterns are produced

No patterns were observed in a wide range of conditions, including when colonies were grown (1) at 30 °C; (2) with 10 μM DAPG (Fig. S**16**). The central GFP-rich region is a common feature observed in most of the imaged conditions. This central GFP ring/disc forms at day 1 at 37 °C and is generally maintained throughout the experiment. On the other hand, the concentric stripes are only observed with 10 μM ATC (Fig. 1d, 3).

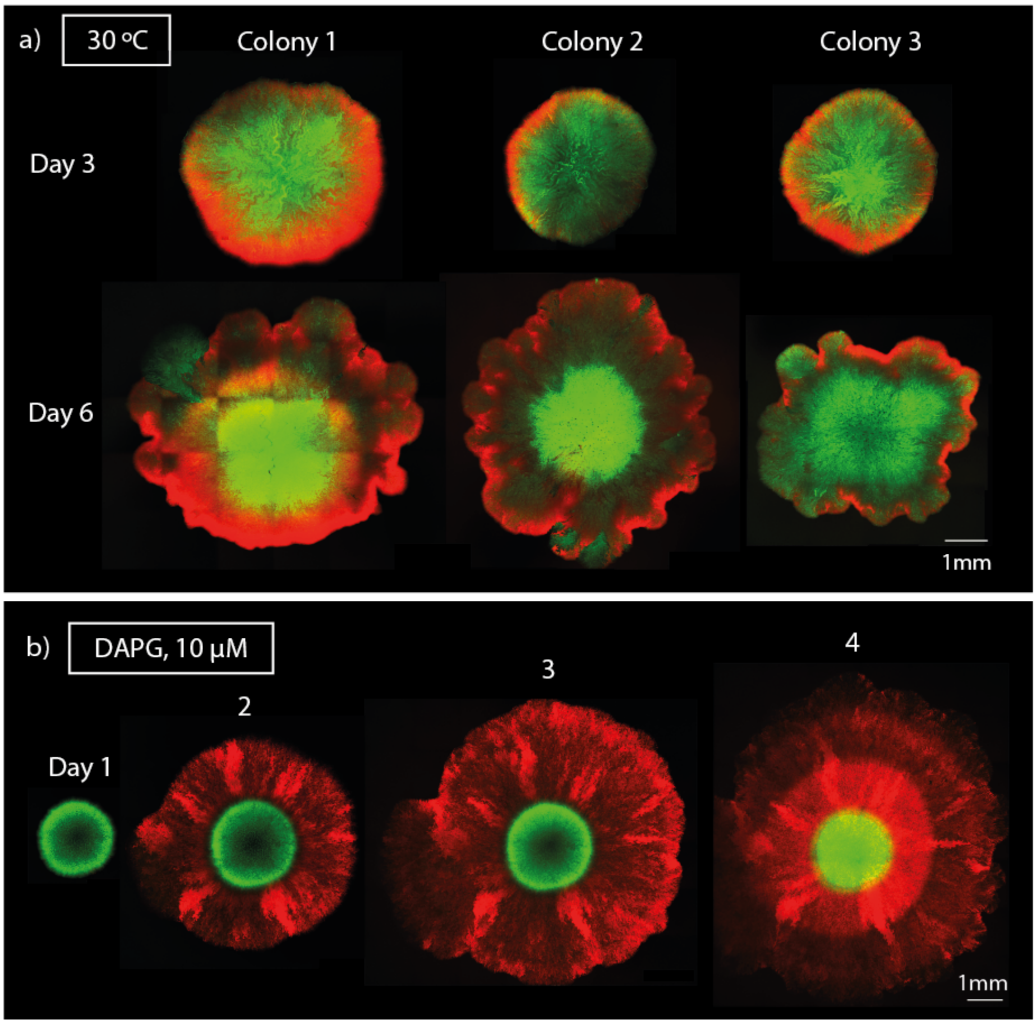

**Supplementary Figure 16: no patterns observed at 30 °C or with 10 μM DAPG.** a) No patterns are consistently observed when colonies are grown at 30 °C either in the GFP or mCherry channel, 3 days, or 6 days after incubation. The GFP-rich centre is a feature that is also observed at 37 °C (Fig. 3); however, no concentric stripes are formed at 37 °C. b) Similarly to all other patterning conditions, the central GFP ring forms at day 1 and is maintained throughout the 4 days; however, no stripes are observed in the presence of 10 μM DAPG.

### Supplementary 3: Model Derivation

In this section, a partial differential equation model of the synthetic gene circuit is presented. This model includes constant background production, activator/repressor-regulated production using Hill terms, and the effect of tuning molecules aTc and IPTG on the circuit and diffusion. The dynamics of the protein (X) and diffuser (U) species of the gene circuit are modelled as

$$\frac{\partial[X]}{\partial t} = b_X + V_X \cdot \frac{1}{1+\left(\frac{KA}{[A]}\right)^{n_A}} \cdot \frac{1}{1+\left(\frac{[I]}{K_I}\right)^{n_I}} - \mu_X \cdot [X] \qquad \text{1a}$$

$$\frac{\partial[U]}{\partial t} = k_1 \cdot [A] - \mu_U \cdot [U] + D_U(\partial_{xx} + \partial_{yy})[U] \qquad \text{1b}$$

This results a PDE model with 8 equations that corresponds to the 2 diffusers (pC and $OC_{14}$) and 6 proteins species of the circuit: RpaI, CinI, TetR, LacI, cI* and cI. This PDE model is reduced to a 6-equation system by assuming quasi-steady state for the diffusers, with production and degradation kinetics much faster than those of proteins. Finally, the model is non-dimensionalised to enable its parametrisation using liquid culture dose response data.

In the subsections below, the different terms of Eq. 1a,b are discussed. Furthermore, the process of model reduction, non-dimensionalisation and fitting is also described.

##### 3.1. Protein equations and gene regulation

The rate of protein production can be defined as

$$V = V_{max} \cdot \Theta \qquad \text{2}$$

where Θ represents the fractional activation of the system. Full activation of protein production is denoted as $\Theta = 1$ while full inhibition is represented by $\Theta = 0$. $V_{max}$ is the maximal rate of expression. $\Theta$ can be represented by a Hill function to describe cooperative binding, derived using the law of mass action with all-or-none binding to multiple binding sites[7]. We further apply the quasi-steady state assumptions for activator and inhibitor binding to the promoter, as well as for the mRNA dynamics, as these timescales are much faster than protein production[8,9]. This leads to the following expression of $\Theta$ for non-competitive activation (A) and inhibition (I)

$$\Theta = \frac{1}{1+\left(\frac{KA}{[A]}\right)^{n_A}} \cdot \frac{1}{1+\left(\frac{[I]}{K_I}\right)^{n_I}} \quad 3$$

This Hill function is used in the model equations (Eq. 3) and describes promoter activity as a function of the two inputs [A] and [I], where $K_A$ and $K_I$ are half-activation/inhibition concentrations, $n_A$ and $n_I$ are the Hill coefficients. Additionally, most promoters are leaky, which we account for by introducing a small rate of background production $b_X$. This $b_X$ corresponds to the first term in Eq. 1a.

##### 3.2. Tuning molecules: aTc regulation of TetR and IPTG regulation of LacI

The circuit was designed so it can be tuned in a variety of ways. Experimentally, this tuning was used to achieve parameter combinations that are more favourable for patterning. The tuning can be performed with: aTc, IPTG and DAPG. The following section introduces aTc and IPTG tuning into the model.

aTc binds to TetR and inactivates it. Only free, unbound TetR can bind TetO and inactivate the expression of cI*. The binding of aTc to TetR is modelled by a reversible equilibrium. The affinity of binding is given by the $k_{on}$ and $k_{off}$ rate constants.

$$naTc + TetR_{free} \underset{kon}{\overset{koff}{\rightleftharpoons}} TetR\text{-}naTc \quad 4$$

This equilibrium happens at much faster rates than the protein production and degradation reactions of the model. For this reason, quasi-steady state is assumed

$$\frac{\partial TetR\text{-}naTc}{\partial t} = k_{on}[TetR_{free}][aTc]^n - k_{off}[TetR\text{-}naTc] \approx 0 \quad 5$$

By rearranging terms and using $K_D = k_{off}/k_{on}$, a simplified expression for $TetR_{free}$ is obtained, which is then used in the model

$$[TetR_{free}] = K_D \cdot \frac{[TetR\text{-}naTc]}{[aTc]^n} \quad 6$$

However, because [TetR-naTc] is unknown, the [$TetR_{total}$] and [$TetR_{free}$] are used instead

$$[TetR\text{-}naTc] = [TetR_{total}] - [TetR_{free}] \quad 7$$

to obtain the following expression

$$[TetR_{free}] = K_D \cdot \frac{[TetR_{total}] - [TetR_{free}]}{[aTc]^n} \Longrightarrow \quad 8a$$

$$[TetR_{free}](1 + \frac{K_D}{[aTc]^n}) = \frac{K_D \cdot [TetR_{total}]}{[aTc]^n} \Longrightarrow \quad 9b$$

$$[TetR_{free}] = \frac{K_D \cdot [TetR_{total}]}{[aTc]^n + K_D} = \frac{[TetR_{total}]}{1 + \frac{[aTc]^n}{K_D}} \quad 10c$$

which says that the concentration of unbound TetR is given by [aTc], [$TetR_{total}$] and the equilibrium constant $K_D$.

For non-cooperative systems where $n = 1$, $K_D = [Lhalf]$. However, for cooperative systems where $n > 1$, $[Lhalf] = \sqrt[n]{K_D}$. We define a new variable $K_A = \sqrt[n]{K_D} \Leftrightarrow K_D = K_A^n$ that we use to replace $K_D$ in the following expression, where ${K_A}^n = {K_{TetR-aTc}}^{n_{aTc}}$. This was also done for the other $K_D$ parameters of the model.

$$[TetR_{free}] = \frac{[TetR_{total}]}{1 + (\frac{[aTc]}{K_{TetR\text{-}aTc}})^{n_{aTc}}} \quad 11$$

Parameter $K_{TetR\text{-}aTc}$ is the binding affinity of aTc to TetR, whereas $n_{aTc}$ is the cooperativity of TetR and aTc binding.

The same logic can be applied for the IPTG regulating LacI inhibition, leading to the following expression for unbound LacI [$LacI_{free}$].

$$[LacIfree] = \frac{[LacI_{total}]}{1 + (\frac{[IPTG]}{K_{LacI\text{-}IPTG}})^{n_{IPTG}}} \quad 12$$

##### 3.3. Degradation

All the species of the circuit were modelled to undergo linear (first order) degradation

$$-\mu_{P_1}[P_1] \quad 13$$

as seen in Eq. 1a and 1b. The degradation rate parameters for the protein species are readily available in the literature[8,10] (Table S**2**). They are scale-free parameters that only depend on time, which makes them more translatable between different experimental contexts and units of measurement.

**Supplementary Table 2:** degradation coefficients of protein species with degradation tags LVA, AAV and ASV[8]. The proteins of the circuit were added short peptide sequences at their 3' ends, known as degradation tags, that promote their breakdown by the cell's proteolytic enzymes.

| Degradation tag | Species | Rate [$h^{-1}$] |
|---|---|---|
| LVA | CinI, LacI, cI, cI*, TetR | 1.14 |
| AAV | / | 0.63 |
| ASV | RpaI, GFP, mCherry | 0.30 |

##### 3.4. Diffuser receptors

The circuit receptors are expressed constitutively from a low-copy pCC1 plasmid, and are therefore modelled with a constant concentration.

Quasi-steady state was assumed for the very fast equilibrium that forms between the receptor and the diffusers. The receptor-inducer and receptor-promoter binding equilibria were modelled with mass action kinetics.

##### 3.5. Diffuser equations

The enzymatic production of the diffusers was modelled with a simple linear production term dependent on synthesis enzyme concentration ([A], [B]) and a rate constant ($k_1$, $k_2$). It was assumed that the precursor substrate concentration is in excess and does not influence the reaction rate.

The diffusers were also modelled to undergo linear degradation, parameterised by $\mu_U$ and $\mu_V$. This is a combination of a spontaneous hydrolysis in water, enzymatic AiiA-dependent degradation, and other cellular metabolic processes[11–13].

The diffusers' movement through space was modelled with simple diffusion terms and parameterised with diffusion coefficients $D_U$ and $D_V$

$$\frac{\partial[U]}{\partial t} = k_1 \cdot [A] - \mu_U \cdot [U] + D_U(\partial_{xx} + \partial_{yy})[U] \qquad 14a$$

$$\frac{\partial[V]}{\partial t} = k_2 \cdot [B] - \mu_V \cdot [V] + D_V(\partial_{xx} + \partial_{yy})[V] \qquad 14b$$

The time dynamics of diffuser synthesis is much faster than that of protein production. Therefore, the rate of change of diffuser is expected to be much higher than that of proteins. These rapid fluctuations of diffuser can be approximated to equilibrium by using the quasi steady state approximation, meaning the rate is set to zero. This leads to an expression of diffuser concentration that is linearly correlated with their synthesis enzyme and rate constant, and inversely correlated with their degradation rate

$$\frac{\partial[U]}{\partial t} = k_1 \cdot [A] - \mu_U \cdot [U] = 0 \longrightarrow U = \frac{k_1}{\mu_U}[A] \qquad 15a$$

$$\frac{\partial[V]}{\partial t} = k_2 \cdot [B] - \mu_V \cdot [V] + D_V \cdot \Delta[V] \longrightarrow V = \frac{k_2}{\mu_V}[B] \qquad 15b$$

##### 3.6. Full 6-equation model

All the terms described above (basal production, regulated production, linear degradation, and diffusion) are combined into an 8-equation PDE system that describes protein concentrations and diffusers. Then, the model is reduced into a 6-equation system where the quasi-steady state diffuser expressions (Eq. 15a and 13b) are substituted into the protein equations both in the diffusion terms and in the Hill terms. This was done with the aim to obtain the simplest possible model, describing the slowest processes of the system[8–10]. The model dependent variables that describe protein concentrations are listed in Table S3.

**Supplementary Table 3:** list of dependent variables, the concentrations the circuit molecules.

| Variable | Molecule |
|---|---|
| [A] | RpaI |
| [B] | CinI |

| | |
|---|---|
| [C] | TetR |
| [D] | LacI |
| [E] | cI* |
| [F] | cI |

In this process, the diffusion of U and V is artificially assigned to A and B, since

$$U = \frac{k_1}{\mu_U}[A] \longrightarrow D_U \nabla^2[U] = D_U \frac{k_1}{\mu_U} \nabla^2[A] \qquad \text{16a}$$

$$V = \frac{k_2}{\mu_V}[B] \longrightarrow D_V \nabla^2[V] = D_V \frac{k_2}{\mu_V} \nabla^2[B] \qquad \text{16b}$$

This results in the following system of six equations:

$$\frac{\partial[A]}{\partial t} = b_A + V_A \cdot \frac{1}{\left(1 + \left(\frac{[D]}{K_{da}}\right)^{n_{da}}\right)} - \mu_A \cdot [A] + \frac{k_1 D_U}{\mu_U} \cdot \nabla^2[A] \qquad \text{17a}$$

$$\frac{\partial[B]}{\partial t} = b_B + V_B \cdot \frac{1}{1 + \left(\frac{\mu_u Kab}{k_1[A]}\right)^{n_{ab}}} \cdot \frac{1}{1 + \left(\frac{[E]}{K_{eb}}\right)^{n_{eb}}} - \mu_B \cdot [B] + \frac{k_2 D_V}{\mu_V} \cdot \nabla^2[B] \qquad \text{17b}$$

$$\frac{\partial[C]}{\partial t} = b_C + V_C \cdot \frac{1}{\left(1 + \left(\frac{[D]}{K_{da}}\right)^{n_{da}}\right)} - \mu_C \cdot [C] \qquad \text{17c}$$

$$\frac{\partial[D]}{\partial t} = b_D + V_D \cdot \frac{1}{1 + \left(\frac{\mu_V Kbd}{k_2[B]}\right)^{n_{bd}}} - \mu_D \cdot [D] \qquad \text{17d}$$

$$\frac{\partial[E]}{\partial t} = b_E + V_E \cdot \frac{1}{\left(1 + \left(\frac{[C]}{K_{ce}}\right)^{n_{ce}}\right)} \cdot \frac{1}{1 + \left(\frac{[F]}{K_{fe}}\right)^{n_{fe}}} \cdot \frac{1}{1 + \left(\frac{Kee}{[E]}\right)^{n_{ee}}} - \mu_E \cdot [E] \qquad \text{17e}$$

$$\frac{\partial[F]}{\partial t} = b_F + V_F \cdot \frac{1}{1+\left(\frac{\mu_V K_{bd}}{k_2[B]}\right)^{n_{bd}}} - \mu_F \cdot [F] \qquad \text{17f}$$

Additionally, $K_{da}$ and $K_{ce}$ are expressed in terms of their respective tuneable molecules as the following

$$K_{da} = K_{lacI-lacO}\left(1+\frac{[IPTG]}{K_{lacI-IPTG}}\right)^{n_{IPTG}} \qquad \text{18a}$$

$$K_{ce} = K_{tetR-tetO}\left(1+\frac{[aTc]}{K_{tetR-aTc}}\right)^{n_{aTc}} \qquad \text{18b}$$

The parameter $K_{TetR\text{-}TetO}$ is the binding affinity of TetR to tetO, $K_{tetR\text{-}aTc}$ is the affinity of aTc to TetR, $n_{aTc}$ is the Hill coefficient of aTc and TetR, whereas $n_{da}$ is the Hill coefficient of TetR to tetO. Similarly, $K_{lacI\text{-}lacO}$ is the affinity of LacI to lacO, $K_{lacI\text{-}IPTG}$ is the affinity of IPTG to LacI, $n_{IPTG}$ is the Hill coefficient of IPTG to LacI, whereas $n_{ce}$ is the Hill coefficient of LacI to lacO.

##### 3.7. Dimensionless model

A dimensionless model was derived to better understand the nature of the parameters, and to simplify the fitting process, as is explained below. The non-dimensionalisation involved the following transformations of dependent and independent variables of concentration (X), time (t) and space (x and y):

$$X = \frac{b_x}{\mu_x}X^*, t = \frac{t^*}{\mu_a}, x = \sqrt{\frac{k_1 D_u}{\mu_a \mu_u}}x^*, y = \sqrt{\frac{k_1 D_u}{\mu_a \mu_u}}y^* \qquad \text{19}$$

And the following transformations of the system parameters V, μ, K and D:

$$V_x^* = \frac{V_x}{b_x}, \mu_x^* = \frac{\mu_x}{\mu_a}, K_{yx}^* = \frac{\mu_x}{b_x}K, D_r = \frac{k_2 D_v \mu_u}{k_1 D_u \mu_v} \tag{20}$$

This leads to the following dimensionless model:

$$\frac{\partial[A^*]}{\partial t^*} = 1 + V_a^* \left( \frac{1}{1 + \left( \frac{[D^*]}{K_{da}^*} \right)^{n_{da}}} \right) - [A^*] + (\partial_{xx} + \partial_{yy})[A^*] \tag{21a}$$

$$\frac{\partial[B^*]}{\partial t} = \mu_b^* \left( 1 + V_B^* \left( \frac{1}{1 + \left( \frac{\mu_u K u b^*}{k_1 [A^*]} \right)^{n_{ab}}} \right) \cdot \left( \frac{1}{1 + \left( \frac{[E^*]}{K_{eb}^*} \right)^{n_{eb}}} \right) - [B^*] \right) + D_r (\partial_{xx} + \partial_{yy})[B^*] \tag{21b}$$

$$\frac{\partial[C^*]}{\partial t^*} = \mu_c^* \left( 1 + V_c^* \left( \frac{1}{1 + \left( \frac{[D^*]}{K_{da}^*} \right)^{n_{da}}} \right) - [C^*] \right) \tag{21c}$$

$$\frac{\partial[D^*]}{\partial t^*} = \mu_d^* \left( 1 + V_d^* \left( \frac{1}{1 + \left( \frac{\mu_v K_{vd}^*}{k_2 [B^*]} \right)^{n_{vd}}} \right) - [D^*] \right) \tag{21d}$$

$$\frac{\partial[E^*]}{\partial t^*} = \mu_e^* \left( 1 + V_e^* \left( \frac{1}{1 + \left( \frac{[C^*]}{K_{ce}^*} \right)^{n_{ce}}} \right) \left( \frac{1}{1 + \left( \frac{[F^*]}{K_{fe}^*} \right)^{n_{fe}}} \right) \left( \frac{1}{1 + \left( \frac{K_{ee}^*}{[E^*]} \right)^{n_{ee}}} \right) - [E^*] \right) \tag{21e}$$

$$\frac{\partial[F^*]}{\partial t^*} = \mu_f^* \left( 1 + V_f^* \left( \frac{1}{1 + \left( \frac{\mu_v K_{vd}^*}{k_2 [B^*]} \right)^{n_{vd}}} \right) - [F^*] \right) \tag{21f}$$

A summary of the model parameters including notation, original units and dimensionless units can be found in Table S4.

**Supplementary Table 4:** Model parameters and units.

| Parameter | Description | Units | Dimensionless Units |
|---|---|---|---|
| $X^*$ | Molecular species | $nM$ | $\frac{\mu_x}{b_x}X \quad [\frac{nM/h}{nM/h} = 1]$ |
| $b^*_x$ | Background production rate | $nM/h$ | no $b_x$ |
| $V^*_x$ | Induced max production rate | $nM/h$ | $\frac{V_x}{b_x} \quad [\frac{nM/h}{nM/h} = 1]$ |
| $K^*_{yx}$ | Concentration for half max response | $nM$ | $\frac{\mu_x}{b_x}K_{yx} \quad [\frac{nM/h}{nM/h} = 1]$ |
| $n_{yx}$ | Cooperativity constant | 1 | 1 |
| $\mu^*_x$ | Degradation rate | $1/h$ | $\frac{\mu_x}{\mu_a} \quad [\frac{1/h}{1/h} = 1]$ |
| $D_r$ | Diffusion rate | $mm^2/h$ | $\frac{k_2 D_v \mu_u}{k_1 D_u \mu_v} \quad [1]$ |
| $t^*$ | Time | $h$ | $\mu_a \cdot t \quad [h \cdot h^{-1} = 1]$ |
| $x^*$ | Space | $mm$ | $\sqrt{\frac{k_1 D_1}{\mu_a \mu_v}}/x \quad [\sqrt{\frac{h^{-1}mm^2h^{-1}}{h^{-1}h^{-1}}}/mm = \frac{\sqrt{mm^2}}{mm} = 1]$ |

##### 3.8. Study of the model and circuit "matching" or "balancing"

In each system equation, when the gene is completely 'turned off' and the Hill factor is zero, the species concentration [X] reaches the 'basal steady state' $[X]_{ss0}$, whereas when the gene is completely 'turned on' and the Hill term is one, the species concentration [X] moves to the 'induced steady state' $[X]_{ss1}$. These two limits are given by

$$[X]_{ss_0} = \frac{b_X}{\mu_X};\ [X]_{ss_1} = \frac{b_X + V_X}{\mu_X} \qquad 22$$

Briefly, all the concentration variables X* of the dimensionless model are expressed as a fold change from the basal steady state $[X]_{ss0}$, where $[X]^* = [X]/[X]_{ss}$, as in Eq. 19. Hence, the steady state levels of every species $[X]^*$ range from 1 when they are equal to the basal steady state $[X]_{ss0}$ to $(1+V_x^*)$ when they are equal to the induced steady state $[X]_{ss1}$:

$$[X]^*_{ss_0} \leqslant [X]^* \leqslant [X]^*_{ss_1} \;\Rightarrow\; 1 \leqslant [X]^* \leqslant (1 + V_X^*) \qquad 23$$

When an upstream regulator activates/represses a downstream component, the associated $K_{xy}^*$ parameter determines the concentration for the half-maximal response. $K_{xy}^*$ is also expressed relative to the 'basal steady state' of the associated molecule $[X]^*$. For example, if $K_{xy}^*$ equals to 2 then a two-fold induction of [X] from its basal steady state will lead to a half-maximal response in the downstream component. This is given by

$$K_{xy} = \frac{b_X}{\mu_X} K^*_{xy} \;\Rightarrow\; K^*_{xy} = \frac{K_{xy}}{[X]_{ss_0}} \qquad 24$$

Hence, an upstream component is said to be 'matched' with a downstream component when the $K_{xy}^*$ of the downstream component falls within the range of steady state concentrations of the upstream molecule, between $[X]_{ss0} = 1$ and $[X]_{ss1} = (1+V_x^*)$:

$$1 \leqslant K^*_{xy} \leqslant (1 + V_X^*) \qquad 25$$

The closer $K_{xy}^*$ is to these upper and lower bounds, the less balanced the upstream component is with the downstream component. For example, with $K_{xy}^*$ smaller than 1, given that $[X]^*$ can never approach a value smaller than its basal level of 1 at steady state, $[X]^*$ would always be high enough to cause near maximal regulation of the downstream component. On the other hand, with $K_{xy}^*$ larger than $(1+V_x^*)$, $[X]^*$ could never become high enough to cause near maximal regulation of the downstream component.

The matching of the input/output relationships is illustrated with a simple simulation in Fig. S**17**, where the output of upstream component $[X]^*$ is being matched to the input of downstream component $[Y]^*$. In a nutshell, there needs to be a correspondence between the expression strength of the upstream component (determined by $V_x^*$), with the sensitivity of the downstream component (determined by $K_{xy}^*$). "Matched" cases occur when $1 < K_{xy} < V_{max} + 1$, "borderline" cases occur when $0.1 < K_{xy} < 10\,V_{max}$ (and are not matched) and finally "unmatched" cases occur for $K_{xy} < 0.1 \; or \; K_{xy} > 10\,V_{max}$.

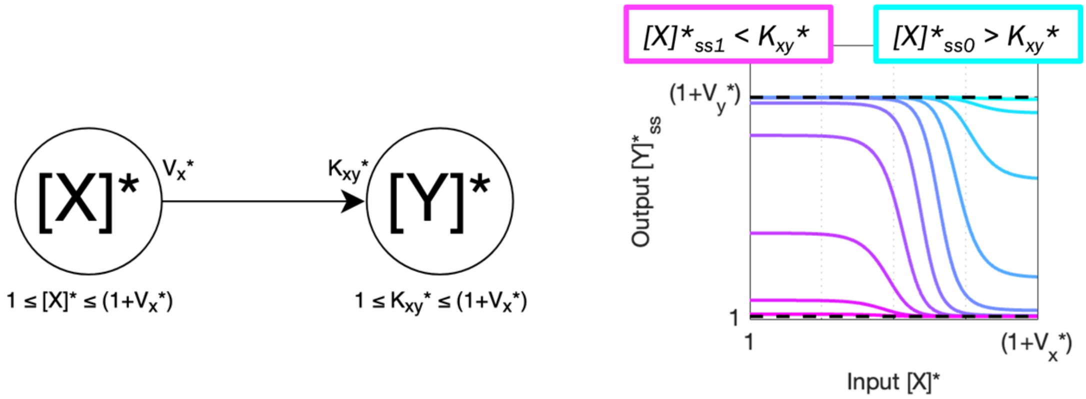

**Supplementary Figure 17: upstream component [X]* acting on downstream component [Y]*.** The downstream component is always 'off' when the upstream component's [X]* fully induced steady-state $[X]^*_{ss1}$ is still lower than the concentration required for a downstream response $K_{xy}^*$ (magenta). Conversely, the downstream component is always 'on' when the upstream component's [X]* basal steady-state $[X]^*_{ss0}$ is high enough for a downstream response $K_{xy}^*$ (cyan).

### Supplementary 4: Model Parameters

In this section, we describe how the model was parametrised using different types of parameter distributions. First, a broad literature-based distribution was used to understand the overall dynamical and patterning properties of the genetic circuit. Then, the circuit was parametrised using liquid culture data and machine learning approaches to constrain the parameters to realistic experimental values. All these parameter regimes were explored using both numerical methods and linear stability analysis.

##### 4.1. Broad literature-based distributions

The model was first studied by initialising the model parameters from distributions derived from the literature. This distribution enables a full understanding of the gene circuit behaviour in all possible parameter regimes. All sampled parameters in this distribution are balanced to ensure optimal robustness as seen in Fig. 2A, and to be consistent with the experiments. This distribution is used to produce Fig. 1D, Fig.2A and Fig.2B. This is a broad distribution that allowed us to explore the overall potential behaviour of the gene circuit.

The types of distributions used for sampling and the values used for each parameter are listed in Table S5. To obtain dimensionless parameters, the bounds of the distributions were recalculated, and the distributions were sampled between these new bounds as specified in the 'Distribution' column of Table S5. For example, for V* (*V/b*) a loguniform distribution was sampled between new bounds [1, 1000].

**Supplementary Table 5: literature-derived parameter distributions.** The distributions used include loguniform and Gaussian. Some parameters were fixed to single values and not sampled. These parameters are dimensional, so these distributions are then used in the dimensionless form of parameters shown in Eq. 17 and Eq. 18.

| Parameter | Distribution | Value |
|---|---|---|
| $V_X$ | Loguniform | 1-100 |
| $b_X$ | Loguniform | 0.1-1 |
| $k_1$, $k_2$ | Loguniform | 0.0183 |
| $D_u$, $D_v$ | Loguniform | 0.1-10 |
| $\mu_u$ , $\mu_v$ | Loguniform | 0.0225 |
| $\mu_{LVA}$ | Gaussian | mean=1.143, std=mean*0.1 |
| $\mu_{ASV}$ | Fixed | 0.3 |

| | | |
|---|---|---|
| $K_X$ | Loguniform | 0.1-1000 |
| $K_{ee}$ | Fixed | 0.01 |
| $n_{vd}$ | Fixed | 2 |
| $n_{ub}$ | Fixed | 1 |
| $n_{da}$ | Fixed | 2 |
| $n_{fe}$ | Fixed | 5 |
| $n_{ee}$ | Fixed | 4 |
| $n_{eb}$ | Fixed | 4 |
| $n_{ce}$ | Fixed | 3 |

The distributions shown in Fig. S**18** were obtained by using the sampling strategies defined in Table S**5**.

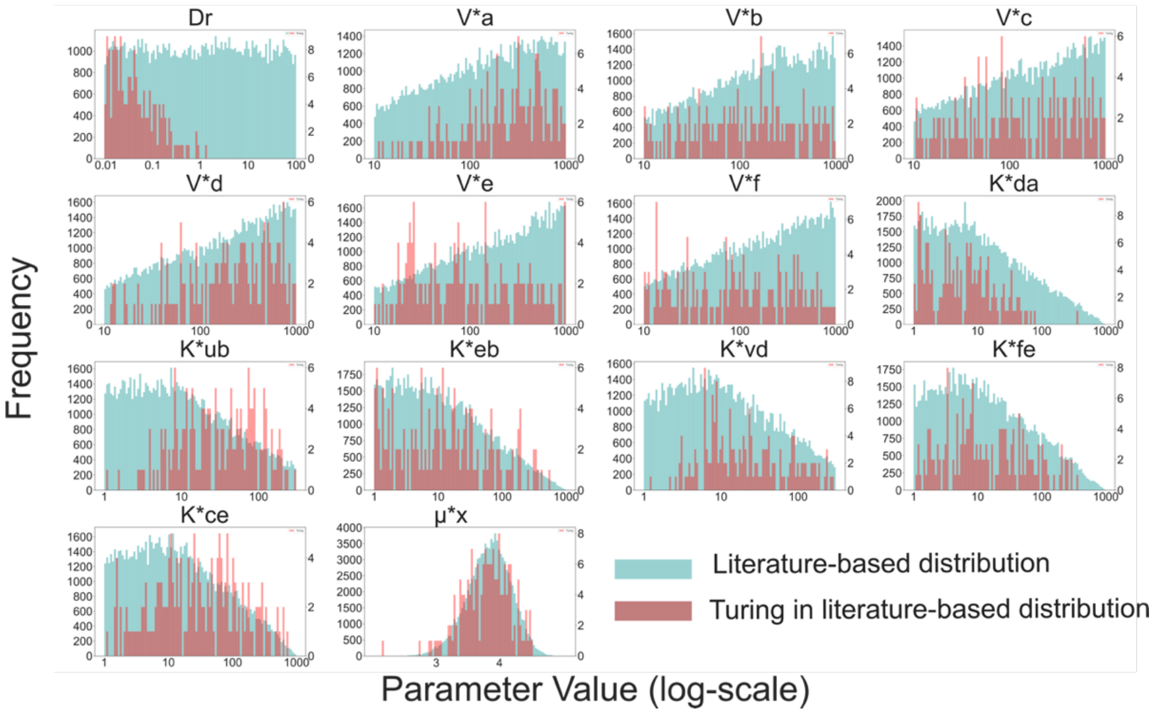

**Supplementary Figure 18: Broad literature-based distributions for model parameters.** Distributions for dimensionless parameters of model based on literature ranges. Blue distributions correspond to the sampled literature-informed distributions, where we made sure to only consider parameters where all the circuit transfer functions are 'matched'. Red distributions correspond to Turing parameter sets found in the blue distributions using linear stability analysis.

Turing pattern systems appear to have a bias for certain parameters regions (Fig. S**18**, red). This is very clear for diffusion, where Turing patterns appear in regions with $D_r < 1$ ($D_v$ / $D_u$) meaning $OC_{14}$ should diffuse slower than pC. This is the case experimentally, where the $D_{OC14}/D_{pC}$ ratio is approximately 0.25 in agar (1.4% w/V, 37 ºC) (Tica et al., unpublished observations).

In terms of the tuning molecules (IPTG, ATC and DAPG) we studied how to tune the gene circuit experimentally to increase Turing pattern probability. IPTG and ATC have a positive monotonic relationship with $K_{da}^*$ and $K_{ce}^*$, respectively (Eq. 15a,b). Therefore, to understand the effects of these two exogenous tuning molecules we can look at their respective K parameters. $K_{da}^*$ Turing parameters have a similar distribution to the sampled distribution, implying that IPTG does not affect the robustness of the gene circuit to Turing pattern formation. On the other hand, the $K_{ce}^*$ Turing parameters are more skewed to higher values than in the sampled distribution, suggesting that ATC increases robustness for Turing pattern formation. This is further shown by a more extensive sampling of the same distribution with 5 different $K_{ce}^*$ values and measuring the Turing robustness in each (Fig. 2B). Finally, DAPG increases diffusor degradation by increasing $\mu_u$ and $\mu_v$, however these parameters are hidden in the dimensionless model, where the diffusor equations were integrated into the protein equations. As seen in Eq. 18b and Eq. 17d, $\mu_u$ and $\mu_v$ are linearly correlated with $K_{ub}^*$ and $K_{vd}^*$, respectively. Both $K_{ub}^*$ and $K_{vd}^*$ have skewed distributions towards higher values compared to the sampled distribution. This means that adding exogenous DAPG could also increase robustness of the circuit for pattern formation. Experimentally, no patterns were observed with a high DAPG concentration (Fig. S**16**), but patterns were consistently observed for high ATC concentration (Fig. 3).

##### 4.2. Constrained parametrised distributions: fitting to liquid culture data of gene subcircuits

The model parameters were later constrained using a machine learning approach that fits the experimental data. In a nutshell, the liquid culture fluorescence data of two subcircuits, subcircuit #1 (Fig. S**2**) and #3 (Fig. S**4**) was fitted to submodels of the dimensionless 6-equation system (Eq. 21a-f). This led to a subset of the model parameters with experimentally relevant values that could further guide the analysis in an experimentally relevant direction. Model parameters for Fig. 2C, Fig. 2D and Fig 2BDE are sampled from the fitted distributions.

##### 4.2.1 Steady-state subcircuit equations for fitting

The non-dimensionalisation of the model facilitates the comparison between the model and experimental dose-response curves. The model dose-response curves are transformed to dimensionless units and range from 1 to $V^*_x + 1$, as explained in Section 3.8 (Fig. S**19**, right).

For the experimental data to match the model, it is divided by the smallest fluorescence value within each experiment and is expressed in relative fold-change units (Fig. S**19**, left). Fold-change units can take values from 1 to $F_{max}/F_{min}$ (maximal and minimal fluorescence levels in the original, untransformed dataset, respectively).

Because the dose-response curves of subcircuit #1 and subcircuit #3 are OC14-dependent, the experimental OC14 units (μM) are also non-dimensionalised using the transform

$$[B]^* = \mathrm{OC}_{14} \cdot \frac{\mu_v \mu_b}{k_2 b_B} \qquad 26$$

Now the model dose-response curves and experimental dose-response curves are both expressed on relative scales and are compatible for fitting (Fig. S**19**, bottom).

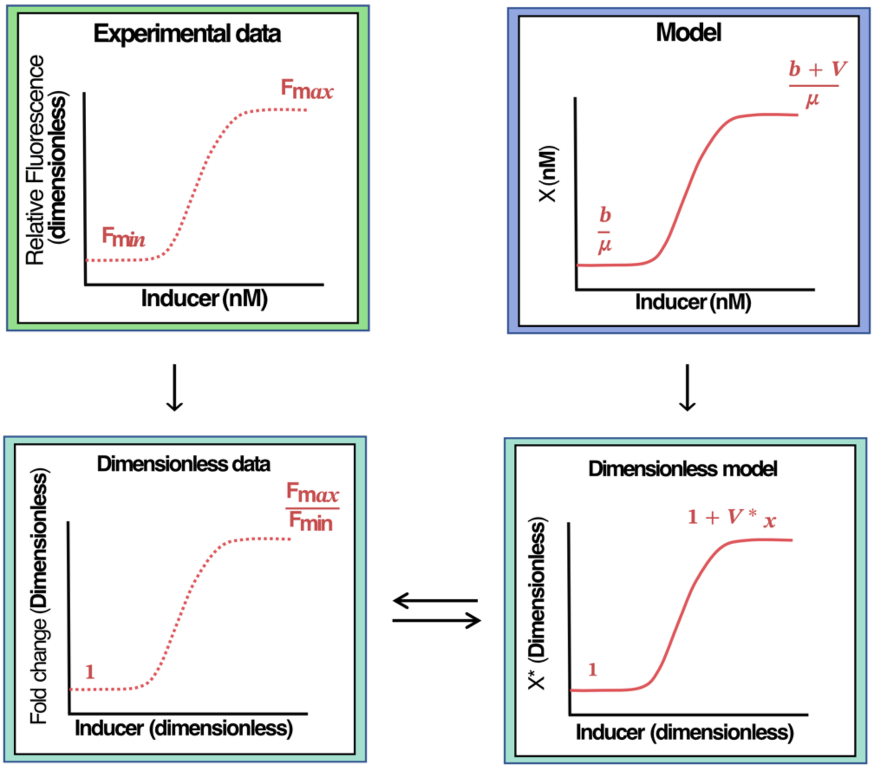

**Supplementary Figure 19: experimental data and model transformation for parametrisation.** Experimental data is scaled by the smallest fluorescence value, so the minimum value is 1. The model is nondimensionalised as explained in the sections above so smallest value is 1 and biggest is $V^*_X - 1$. In both cases, units are dimensionless, and the basal level is 1.

The models for the subcircuits were derived from the main PDE system (Eq. 21a-f). Subcircuit #1 only involves species [F] and [E] (cI and cI*), whereas subcircuit #3 involves species [C], [D] and [E] (TetR, LacI and cI*). All other species were set to zero. For the model parametrisation, we derive steady-state expressions for the dynamically regulated species so that

$$\frac{\partial X}{\partial t} = 0; X = X_{eq} \tag{27}$$

For subcircuit #1 we obtain the following steady-state expressions:

$$[F_1] = 1 + V_f \left( \frac{1}{1 + \left( \frac{\mu_v K_{vd}}{k_v [O_{C14}]} \right)^{n_{vd}}} \right) \tag{28a}$$

$$[E_1] = 1 + V_e \left( 1 + \left( \frac{1 + V_f \left( \frac{1}{1 + \left( \frac{\mu_v K_{vd}}{k_v [O_{C14}]} \right)^{n_{vd}}} \right)}{K_{fe}} \right)^{n_{fe}} \right)^{-1} \tag{28b}$$

And for subcircuit #3 we obtain the following steady-state expressions:

$$[D_3] = 1 + V_d \left( \frac{1}{1 + \left( \frac{\mu_v K_{vd}}{k_v [O_{C14}]} \right)^{n_{vd}}} \right) \tag{29a}$$

$$[C_3] = 1 + V_c \left( \frac{1}{1 + \left( \frac{[D_3]}{K_{da}} \right)^{n_{da}}} \right) \tag{29b}$$

$$[E_3] = 1 + V_e \left( \frac{1}{1 + \left( \frac{[C_3]}{K_{ce}} \right)^{n_{ce}}} \right) \tag{29c}$$

##### 4.2.2 Fitting process and the resulting best fit distributions

In addition to scaling and nondimensionalising, the lowest GFP data points were excluded, because fluorescence readings were insufficiently sensitive to measure concentration at these points (Fig. S**20**). The fits were constrained to the maximal fold-change of the regulation, and to the sensitivity of the regulation (location of half-maximal response). This improved the quality of the fit and allowed us to identify a broader range of suitable solutions.

The two-equation systems (Eq. 28a-b, 29a-c) are fitted independently to the experimental dataset with the python *scipy.optimize.curve_fit* package, which uses the Levenberg-Manquardt to minimise the sum of squared errors (SSE), given by

$$SSE = \sum_{i=1}^{n} (y_i - f(x_i))^2 \qquad 30$$

where $y_i$ is the experimental data and $f(x_i)$ are the two systems of equations parameterised by *Vc, Vd, Vf, Kvd, Kfe,* and *Kce*.

The minimisation algorithm generates a vector of best fit parameters $k$

| $V^*_e$ | $V^*_f$ | $V^*_c$ | $V^*_d$ | $K^*_{vd}$ | $K^*_{fe}$ | $K^*_{da}$ | $K^*_{ce}$ |
|---|---|---|---|---|---|---|---|
| 1.99 | 3.64 | 9.95 | 6.50 | 18.94 | 2.26 | 67.92 | 3.47 |

Two best fit parameters are obtained for $V^*_e$ as this parameter is present in both subcircuit models. However, only $V^*_e$ from subcircuit #1 is considered. These parameters are used to generate the following dose-response curves (Fig. S**20**).

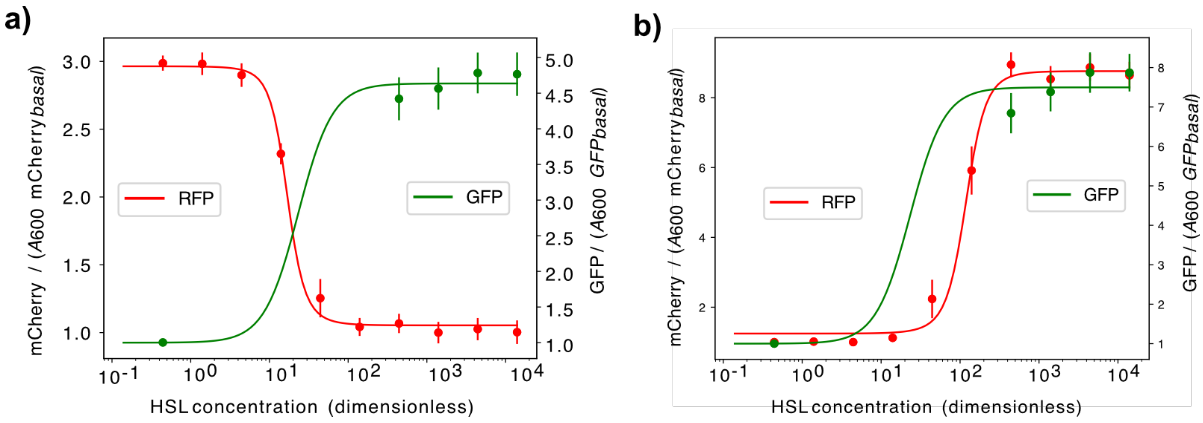

**Supplementary Figure 20: best fit for processed data set.** Processed dataset is fitted using **(a)** Eq. 28a-b for subcircuit #1 and **(b)** Eq. 29a-c for subcircuit #3.

The minimisation algorithm produces a covariance matrix $C_k$, which is the inverse of the Hessian matrix and represents the derivative of the loss function in the different parameter dimensions. In other words, this Hessian matrix represents how the loss increases or decreases when parameters are varied together

$$C_k = H_{L_k}^{-1} \quad 31$$

$$H_{L_{kv}} = \begin{bmatrix} \frac{\partial^2 L}{\partial k_1^2} & \frac{\partial^2 L}{\partial k_1 k_2} & \cdots & \frac{\partial^2 L}{\partial k_1 k_n} \\ \frac{\partial^2 L}{\partial k_2 k_1} & \frac{\partial^2 L}{\partial k_2^2} & \cdots & \frac{\partial^2 L}{\partial k_2 k_n} \\ \vdots & \vdots & \ddots & \vdots \\ \frac{\partial^2 L}{\partial k_n k_1} & \frac{\partial^2 L}{\partial k_n k_2} & \cdots & \frac{\partial^2 L}{\partial k_n^2} \end{bmatrix} \quad 32$$

A multivariate Gaussian distribution is generated using $k$ and $C_k$

$$X \sim \mathcal{N}(k, C_k) \quad 33$$

with a probability density function $p(x; k, C_k)$

$$p(x; k, C_k) = \frac{\exp\left(-\frac{1}{2}(x-k)^T {C_k}^{-1}(x-k)\right)}{\sqrt{(2\pi)^k C_k}} \quad 34$$

The multivariate Gaussian distribution is the generalization of a normal distribution to higher dimensions. For example, for 2-dimensional Gaussian distributions, when the covariance of two parameters X and Y is positive, the parameters are positively correlated (Fig. S**21**, right). This means that if the parameters are increased together, the behaviour of the system should change minimally (and the error to the data should not increase). On the other hand, a negative covariance leads to an inverse correlation of the parameters, meaning when one increases the other should decrease to ensure the error does not increase (Fig. S**21**, left). Finally, a covariance of zero means the X and Y parameters are completely independent, producing a circular distribution (Fig. S**21**, middle).

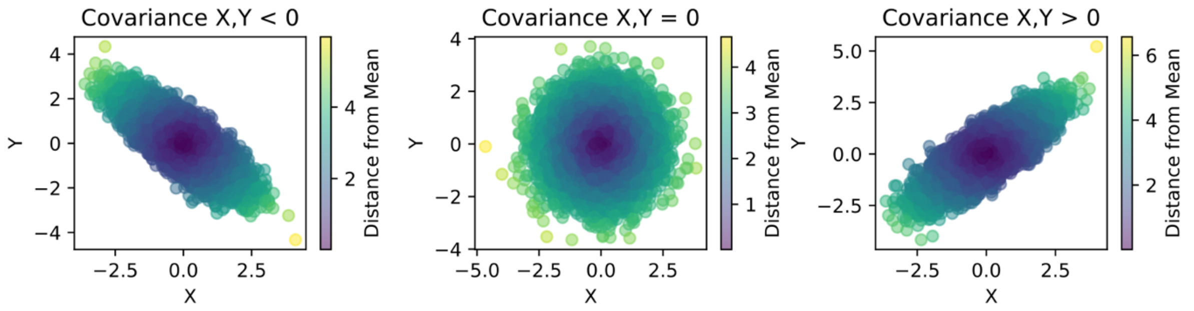

**Supplementary Figure 21: multivariate Gaussian distributions.** X and Y parameter distributions with (A) negative covariance, (B) zero covariance and (C) positive covariance. The colour represents the distance to the mean, or in other words to the best fit parameter *($k$).* Blue is the parameter *($k$)* with minimal loss and yellow is a neighbouring parameter with bigger loss.

The width of the multivariate Gaussian distributions can be increased by multiplying $C_k$ by a scalar factor $q$ where $q > 1$. This process unconstrains the fits and allows more error with respect to the experimental data. To search for Turing patterns close to the fitted parameter combination $k$, the value of $q$ is progressively increased until a Turing parameter set is found through linear stability analysis.

Through this process, the first 3 Turing parameter combinations were found with $q = 10$. These are the closest Turing solutions to the best fit parameter combinations. The dose-response curves generated by the $q = 10$ distribution are shown in Fig. 2c. The $q = 10$ multivariate Gaussian distribution is shown in Fig. S**22**. Within those, 3 curves are generated by Turing parameter sets. The three Turing parameter sets are ensured to be 'balanced', where the input/output relationships of the components are matched (see Section 3.8). Finally, these three parameter sets are simulated using the PDE and cellular automata solver, and are observed to reproduce ring patterns similar to the ones produced by the bacterial colonies (Fig. 2d, Fit #1 and Fit #2).

In Fig. S**22**, we observe some correlations between parameter pairs. Parameters $V*_{c}$ and $K*_{ce}$ exhibit positive correlation. This is what we would expect from the circuit architecture as $V*_{c}$ is the maximum production rate of TetR and $K*_{ce}$ is how much TetR is needed to inhibit node C. In other words, the affinity parameter can be counterbalanced by the expression strength parameter.

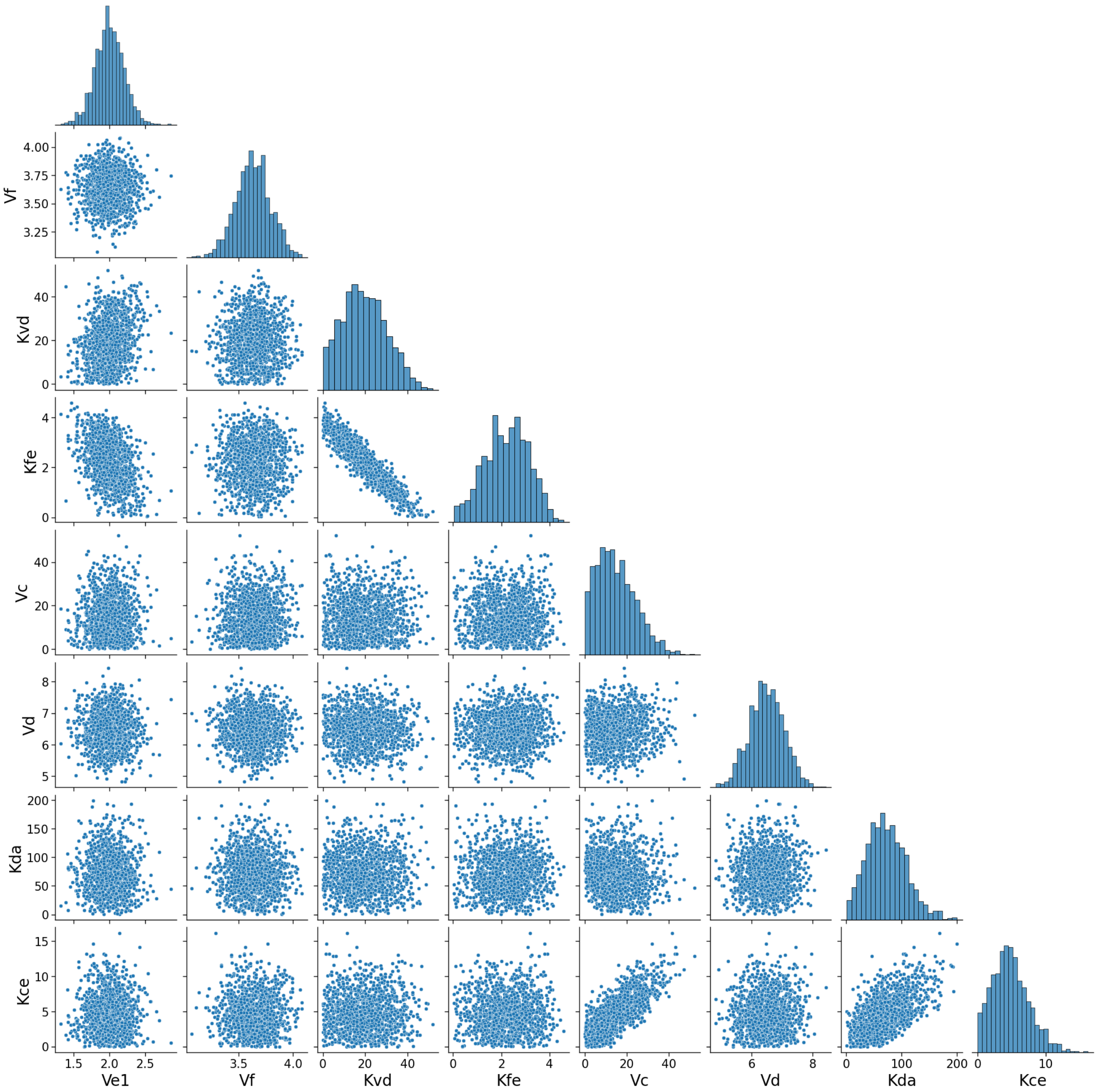

**Supplementary Figure 22: multivariate Gaussian distributions for fitted parameters.** Distributions resulting from fitting subcircuit 1 and 3, using a $q = 10$. Diagonals represent the univariate distributions. The non-diagonals represent the multivariate distributions of parameter pairs. Certain parameters show positive covariance (e.g. Vc and Kce), some negative covariance (e.g. Kvd and Kfe) and some no covariance meaning they are independent (e.g. Vd and Kfe).

Other relationships are observed, such as the negative correlation of $K_{fe}^{*}$ and $K_{vd}^{*}$, which are the Michaelis Menten constants for two consecutive interactions from node B to node C. In biological terms, this means that if the same amount of $OC_{14}$ generates more cI, this can be opposed by more cI being required to repress node C. Therefore, the dynamical behaviour is maintained if these two parameters are varied inversely. All these correlations allowed us to study the circuit dynamics more in depth and to further validate the model.

While our current approach primarily aims to derive model parameters from liquid culture data, it's worth noting that in the future, we could fit to microscopy spatial data with the use of physics-informed neural networks[14].

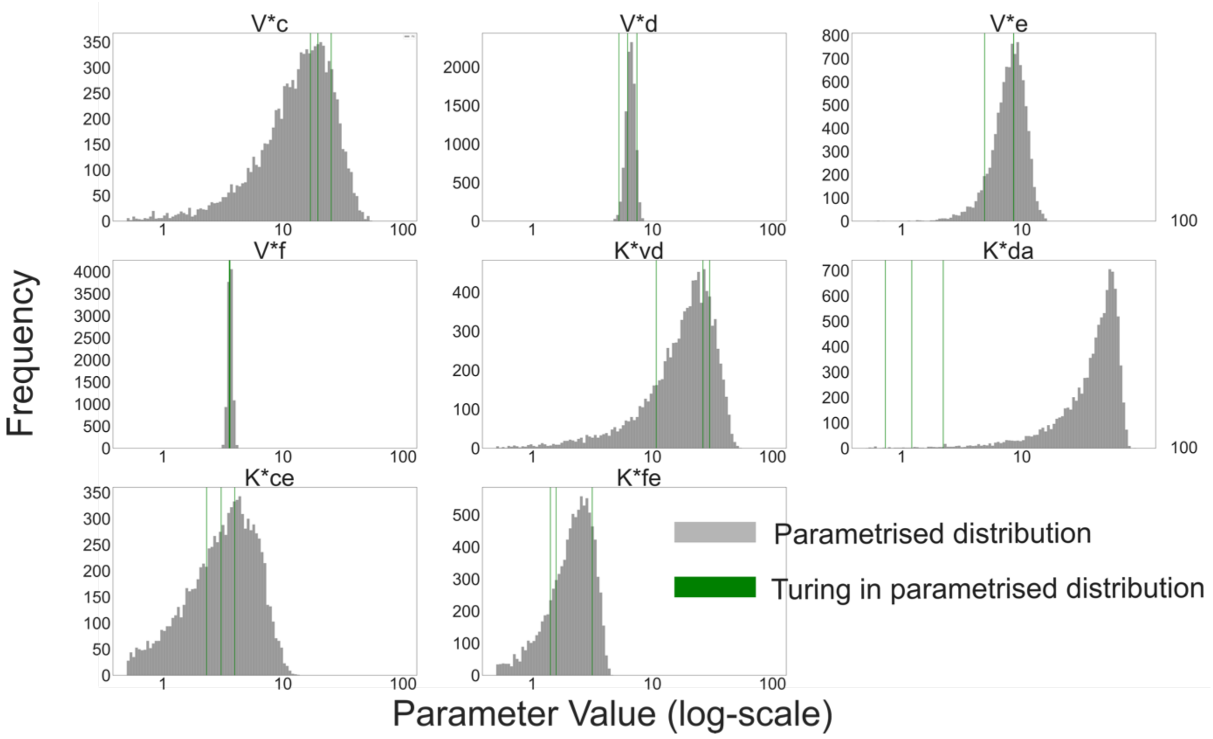

**Supplementary Figure 23: Constrained parametrised distributions for $V^*_X$ and $K^*_{xy}$ parameters.** Distributions for dimensionless parameters of model based on fitting to liquid-culture experimental data. Grey distribution corresponds to parameters from fitting. Green vertical lines correspond to the 3 Turing parameter sets found in grey distribution using linear stability analysis.

##### 4.2.3 Parameter exploration: adding noise to the best fit Turing solutions

To further explore the parameter space found around the fitted Turing parameter sets (Fig. 2d, Fit #1 and Fit #2) a small noise deviation is applied with relative uncertainty of 1%. In other words, for each parameter p, a normal distribution is generated with mean $\mu = p$ and standard deviation $\sigma = p * 0.01$. This small noise perturbation to all parameters, which generates similar steady-state dose response behaviour, allows us to further explore the

Turing parameter space near the best fit to the liquid culture. For each value of uncertainty 2000 parameter combinations were analysed. Simulations using this 'noisy fits' are found in Fig. 2d (Fit + noise #1 and Fit + noise #2) and Fig. 3. The Turing patterning robustness with different amounts of noise is shown in Fig. S**24**, showing how robustness decreases as more noise is added. The different analytical solutions for a relative uncertainty of 1% are shown in Fig. S**24c**, where we observe not only Turing I solutions, but also Turing I Hopf and Hopf solutions.

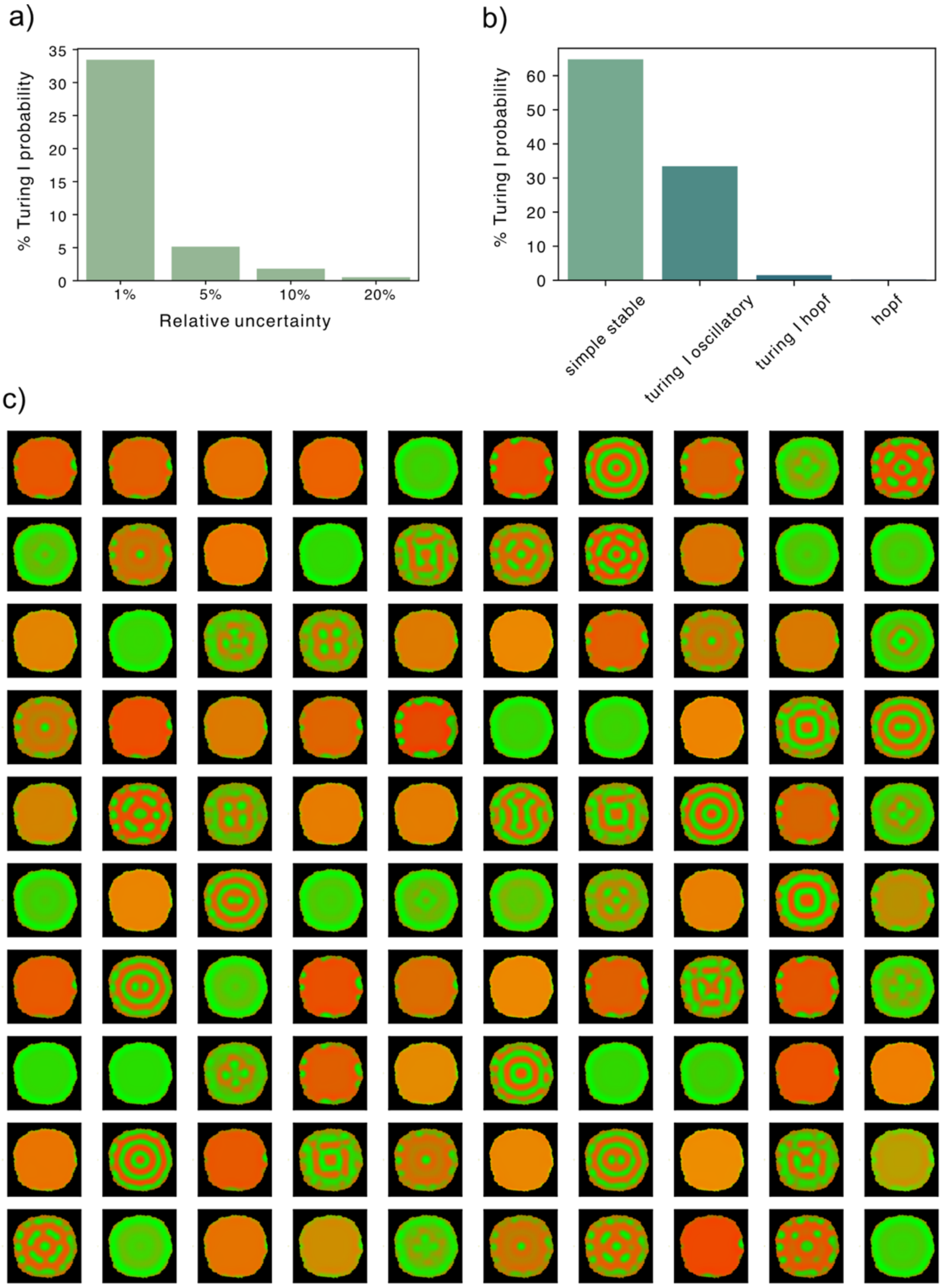

**Supplementary Figure 24: search around fitted Turing parameter set. (a)** Turing robustness with different levels of noise. **(b)** Using 1% noise (mean $\mu = p$ and standard deviation $\sigma = p * 0.01$), frequency of different analytical solutions: Simple stable 64.8%, Turing I oscillatory 33.5%, Turing I Hopf 1.5%, Hopf 0.25%. **(c)** Numerical simulations in growing colonies of 1% noise distribution.

##### 4.3. Summary of model parameters and types of analytical output

Model parameters sampled from these two distributions to produce colony simulations are shown in Table S**6.** Literature-based distributions are used for rows Fig 1D.1 to Fig 2D.2 included. Parametrisation-based distributions are used for rows Fig 2D.3 to Fig 3F Additionally, the analytical solution and growth regimes used for each simulation is shown.

One aspect to highlight is the diffusion ratio Dr of the best fit in Table S**6** (Simulation ID Fig 2B-3F). Assuming that the enzymatic production and degradation rates of the two diffusers pC and OHC14 are equal ($k_1 = k_2$, $\mu_U = \mu_V$), a Dr of 0.15 is very close to an experimentally measured Dr of 0.25 (Tica et al. 2023, unpublished data).

In our models, $K_{ee}$ is fixed to small values as we assume very little cI* is needed to activate itself. This assumption can be justified by looking at the dose response curves which show that the promoter is on by default unless there is inhibition by cI or tetR (See Fig 1B Sub1).

**Supplementary Table 6: model parameters, growth regime and analytical solution of colony simulations.** The analytical solution is derived using linear stability analysis and describes the properties of the steady-states of the system, including Turing I, Hopf, Simple stable, Simple unstable and no steady-states. The kinetic parameters for the dimensionless model are listed here. $\mu_{ASV}$ corresponds to $\mu_a$ while $\mu_{LVA}$ corresponds to $\mu_{b,}$ $\mu_{c,}$ $\mu_d,$ $\mu_{e,}$ $\mu_f$ (see Table S**2**). The growth regimes that determine the PDE and cellular automata parameters are further specified in Table S**7**.

| Simulation ID | Analytical solution | Growth regime | $D_r^*$ | $V_a^*$ | $V_b^*$ | $V_c^*$ | $V_d^*$ | $V_e^*$ | $V_f^*$ | $K_{ub}^*$ | $K_{vd}^*$ | $K_{da}^*$ | $K_{ce}^*$ | $K_{fe}^*$ | $K_{eb}^*$ | $K_{ee}^*$ | $\mu_{LVA}$ * | $\mu_{ASV}^*$ | $n_{ub}$ | $n_{ee}$ | $n_{eb}$ | $n_{vd}$ | $n_{da}$ | $n_{ce}$ | $n_{fe}$ |
|---|---|---|---|---|---|---|---|---|---|---|---|---|---|---|---|---|---|---|---|---|---|---|---|---|---|
| Fig 1D.1 | Turing I | T10 | 0.01 | 150.29 | 54.23 | 28.08 | 266.88 | 45.87 | 10.43 | 29.36 | 42.53 | 2.85 | 100.00 | 1.19 | 17.84 | 0.01 | 3.86 | 1 | 1 | 4 | 4 | 2 | 2 | 3 | 5 |
| Fig 1D.2 | Turing I Hopf Unstable | T10 | 0.09 | 585.33 | 130.73 | 850.91 | 222.44 | 356.89 | 147.60 | 101.86 | 108.87 | 2.00 | 100.00 | 2.43 | 4.60 | 0.01 | 3.96 | 1 | 1 | 4 | 4 | 2 | 2 | 3 | 5 |
| Fig 1D.3 | Turing I Unstable Stable | T25 | 0.02 | 795.88 | 29.76 | 14.26 | 21.69 | 19.04 | 48.66 | 150.98 | 6.42 | 1.46 | 100.00 | 15.15 | 11.34 | 0.01 | 4.46 | 1 | 1 | 4 | 4 | 2 | 2 | 3 | 5 |
| Fig 1D.4 | Turing I | T50 | 0.02 | 45.20 | 603.17 | 234.59 | 158.98 | 70.72 | 13.73 | 13.93 | 239.29 | 1.97 | 100.00 | 2.35 | 60.56 | 0.01 | 3.99 | 1 | 1 | 4 | 4 | 2 | 2 | 3 | 5 |
| Fig 1D.5 | Turing I | T25 | 0.22 | 688.36 | 168.38 | 11.45 | 600.62 | 18.14 | 17.61 | 24.48 | 87.24 | 15.73 | 100.00 | 5.15 | 16.63 | 0.01 | 3.71 | 1 | 1 | 4 | 4 | 2 | 2 | 3 | 5 |
| Fig 1D.6 | Turning I Hopf unstable | T25 | 0.02 | 113.58 | 725.58 | 322.38 | 240.91 | 21.66 | 13.29 | 80.20 | 10.45 | 24.96 | 100.00 | 7.56 | 4.46 | 0.01 | 3.53 | 1 | 1 | 4 | 4 | 2 | 2 | 3 | 5 |
| Fig 1D.7 | Turing I Hopf | T25 | 0.03 | 65.70 | 147.29 | 800.75 | 887.19 | 17.69 | 98.74 | 41.87 | 104.40 | 5.65 | 100.00 | 2.31 | 5.31 | 0.01 | 3.88 | 1 | 1 | 4 | 4 | 2 | 2 | 3 | 5 |
| Fig 2B.1 | Turing I Hopf | T100 | 0.03 | 301.23 | 83.90 | 491.96 | 868.92 | 131.53 | 63.43 | 133.37 | 68.45 | 9.85 | 100.00 | 3.09 | 73.83 | 0.01 | 3.70 | 1 | 1 | 4 | 4 | 2 | 2 | 3 | 5 |
| Fig 2B.2 | Turing I Hopf | T100 | 0.10 | 495.68 | 872.38 | 697.87 | 372.08 | 109.70 | 14.56 | 53.14 | 219.76 | 4.99 | 100.00 | 2.75 | 77.39 | 0.01 | 3.70 | 1 | 1 | 4 | 4 | 2 | 2 | 3 | 5 |
| Fig 2B.3 | Turing I Hopf | T100 | 0.32 | 48.36 | 786.32 | 31.16 | 264.07 | 30.32 | 164.65 | 14.38 | 251.50 | 7.89 | 100.00 | 28.85 | 28.75 | 0.01 | 3.83 | 1 | 1 | 4 | 4 | 2 | 2 | 3 | 5 |
| Fig 2B.4 | Turing I | T100 | 0.01 | 19.53 | 12.34 | 123.40 | 51.73 | 40.95 | 91.98 | 7.07 | 2.98 | 6.18 | 100.00 | 50.56 | 32.69 | 0.01 | 4.56 | 1 | 1 | 4 | 4 | 2 | 2 | 3 | 5 |
| Fig 2B.5 | Turing I | T100 | 0.03 | 193.50 | 22.54 | 626.85 | 106.80 | 402.97 | 39.13 | 13.91 | 4.82 | 6.61 | 100.00 | 23.70 | 336.49 | 0.01 | 3.18 | 1 | 1 | 4 | 4 | 2 | 2 | 3 | 5 |
| Fig 2B.6 | Turing I | T100 | 0.03 | 816.37 | 77.89 | 58.00 | 63.95 | 59.97 | 12.78 | 110.34 | 49.06 | 3.22 | 100.00 | 3.38 | 50.31 | 0.01 | 3.49 | 1 | 1 | 4 | 4 | 2 | 2 | 3 | 5 |
| Fig 2B.7 | Turing I | T100 | 0.02 | 682.27 | 125.41 | 893.16 | 57.16 | 260.91 | 15.47 | 224.29 | 62.41 | 1.70 | 100.00 | 1.49 | 28.51 | 0.01 | 3.94 | 1 | 1 | 4 | 4 | 2 | 2 | 3 | 5 |
| Fig 2B.8 | No steady state | T50 | 9.09 | 74.05 | 169.38 | 669.89 | 41.12 | 39.58 | 76.52 | 15.15 | 120.03 | 1.70 | 100.00 | 10.73 | 5.64 | 0.01 | 3.37 | 1 | 1 | 4 | 4 | 2 | 2 | 3 | 5 |
| Fig 2B.9 | No steady state | T50 | 64.34 | 229.84 | 199.15 | 822.04 | 58.37 | 137.50 | 18.27 | 9.68 | 70.64 | 2.17 | 100.00 | 8.96 | 23.36 | 0.01 | 3.26 | 1 | 1 | 4 | 4 | 2 | 2 | 3 | 5 |
| Fig 2B.10 | Hopf | T50 | 0.16 | 11.99 | 82.80 | 517.54 | 118.80 | 12.02 | 500.10 | 4.90 | 16.60 | 22.21 | 100.00 | 372.67 | 2.19 | 0.01 | 3.59 | 1 | 1 | 4 | 4 | 2 | 2 | 3 | 5 |
| Fig 2B.11 | Hopf | T50 | 3.69 | 138.98 | 718.76 | 731.67 | 574.33 | 893.73 | 274.09 | 45.83 | 231.33 | 3.65 | 100.00 | 127.68 | 5.88 | 0.01 | 3.82 | 1 | 1 | 4 | 4 | 2 | 2 | 3 | 5 |
| Fig 2B.12 | Hopf | T50 | 0.58 | 311.01 | 452.00 | 609.80 | 140.64 | 511.01 | 264.43 | 267.90 | 23.40 | 17.44 | 100.00 | 80.26 | 10.13 | 0.01 | 3.86 | 1 | 1 | 4 | 4 | 2 | 2 | 3 | 5 |
| Fig 2B.13 | Stable Unstable Stable | T25 | 1.25 | 11.76 | 134.85 | 286.24 | 168.35 | 306.73 | 398.68 | 3.15 | 15.85 | 160.35 | 100.00 | 84.35 | 1.32 | 0.01 | 3.13 | 1 | 1 | 4 | 4 | 2 | 2 | 3 | 5 |
| Fig 2B.14 | Stable | T25 | 97.13 | 259.08 | 179.44 | 127.45 | 674.32 | 19.68 | 14.12 | 76.32 | 3.39 | 392.99 | 100.00 | 1.28 | 14.09 | 0.01 | 2.81 | 1 | 1 | 4 | 4 | 2 | 2 | 3 | 5 |
| Fig 2B.15 | Stable | T25 | 25.66 | 825.22 | 30.51 | 130.99 | 30.67 | 166.25 | 630.74 | 238.35 | 4.73 | 13.41 | 100.00 | 1.85 | 147.94 | 0.01 | 3.16 | 1 | 1 | 4 | 4 | 2 | 2 | 3 | 5 |
| Fig 2D.1 | Turing I | T50 | 0.15 | 21.74 | 321.61 | 16.95 | 5.27 | 8.54 | 3.57 | 0.15 | 26.39 | 16.95 | 5.27 | 1.42 | 1.52 | 0.001 | 3.60 | 1 | 1 | 4 | 4 | 2 | 2 | 3 | 8 |
| Fig 2D.2 | Turing I Hopf | T50 | 0.10 | 102.69 | 431.77 | 25.23 | 7.44 | 4.90 | 3.61 | 0.10 | 29.99 | 25.23 | 7.44 | 1.58 | 0.79 | 0.001 | 3.69 | 1 | 1 | 4 | 4 | 2 | 2 | 3 | 8 |
| Fig2D.3 | Turing I | T100 | 0.15 | 21.67 | 321.54 | 16.88 | 5.29 | 8.60 | 3.57 | 43.93 | 26.33 | 0.73 | 2.34 | 1.41 | 1.53 | 0.001 | 3.57 | 1 | 1 | 4 | 4 | 2 | 2 | 3 | 8 |
| Fig 2D.4 | Stable Peak near zero | T100 | 0.15 | 21.94 | 327.41 | 16.71 | 5.30 | 8.55 | 3.58 | 44.06 | 26.51 | 0.72 | 2.31 | 1.41 | 1.53 | 0.001 | 3.62 | 1 | 1 | 4 | 4 | 2 | 2 | 3 | 8 |
| Fig 3C | Turing I | T100 Different time snapshots | 0.15 | 21.67 | 321.54 | 16.88 | 5.29 | 8.60 | 3.57 | 43.93 | 26.33 | 0.73 | 2.34 | 1.41 | 1.53 | 0.001 | 3.57 | 1 | 1 | 4 | 4 | 2 | 2 | 3 | 8 |
| Fig 3D | Turing I | T100 Red and green channels | 0.15 | 21.67 | 321.54 | 16.88 | 5.29 | 8.60 | 3.57 | 43.93 | 26.33 | 0.73 | 2.34 | 1.41 | 1.53 | 0.001 | 3.57 | 1 | 1 | 4 | 4 | 2 | 2 | 3 | 8 |
| Fig 3E.1 | Turing I | T100 | 0.15 | 21.67 | 321.54 | 16.88 | 5.29 | 8.60 | 3.57 | 43.93 | 26.33 | 0.73 | 2.34 | 1.41 | 1.53 | 0.001 | 3.57 | 1 | 1 | 4 | 4 | 2 | 2 | 3 | 8 |
| Fig 3E.2 | Turing I | T100 Open boundary | 0.15 | 21.67 | 321.54 | 16.88 | 5.29 | 8.60 | 3.57 | 43.93 | 26.33 | 0.73 | 2.34 | 1.41 | 1.53 | 0.001 | 3.57 | 1 | 1 | 4 | 4 | 2 | 2 | 3 | 8 |
| Fig 3F | Turing I | T110 Different growth rates | 0.15 | 21.67 | 321.54 | 16.88 | 5.29 | 8.60 | 3.57 | 43.93 | 26.33 | 0.73 | 2.34 | 1.41 | 1.53 | 0.001 | 3.57 | 1 | 1 | 4 | 4 | 2 | 2 | 3 | 8 |

##### 4.4. Linear stability analysis and dispersion relation

All the parameters sets shown in Table S**6** are studied using linear stability analysis to understand their stability profile with and without diffusion. In the context of reaction-diffusion systems, linear stability analysis is carried out to understand the spatial patterns that can evolve over time[15].

For a system of equations describing a two-node network

$$\frac{\partial[A]}{\partial t} = g(A,B) + D_A \cdot \frac{\partial^2[A]}{\partial x^2}$$

$$\frac{\partial[B]}{\partial t} = f(A,B) + D_B \cdot \frac{\partial^2[B]}{\partial x^2}$$

the Jacobian can be obtained by linearising around the steady state and transforming the spatial diffusion into $-Dk^2$ using Fourier transformation where $k$ corresponds to the wavenumber. This leads to the following expression

$$J = \begin{bmatrix} \frac{\partial g_A}{\partial A} - D_A k^2 & \frac{\partial g_A}{\partial B} \\ \frac{\partial f_B}{\partial A} & \frac{\partial f_B}{\partial B} - D_B k^2 \end{bmatrix}$$

The stability of the system is tested by computing the eigenvalues of the Jacobian and testing their sign. A negative eigenvalue indicates stability, while a positive eigenvalue indicates an unstable system. To calculate a dispersion relation, the eigenvalues are computed for various values of $k$ (wavenumber).

The classical Turing instability or diffusion-driven instability is defined by its dispersion relation. The steady state without diffusion ($k = 0$), is stable meaning it has negative eigenvalues. As diffusion is introduced ($k > 0$), the system becomes unstable and eigenvalues become positive. Finally, as $k \to \infty$, the system becomes stable again and eigenvalues drop below zero[15,16].

The analytical solutions in the third column of Table S**6**, are derived from the dispersion relations of our reaction-diffusion model. The different types of dispersion relations that were considered are shown in Fig. S**25**.

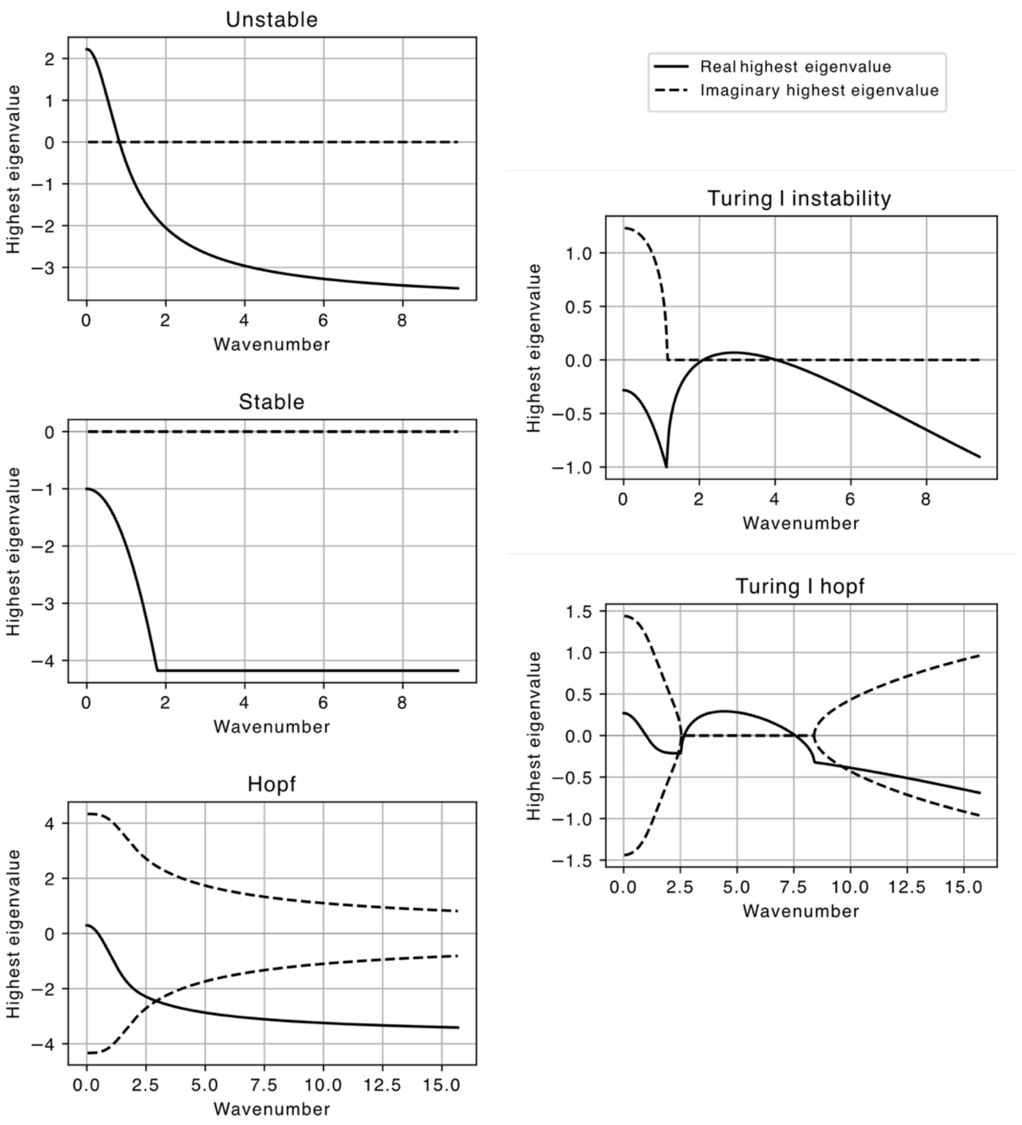

**Supplementary Figure 25: dispersion relations of different types of systems.** These dynamical systems correspond to the systems shown in Fig. 2b. The highest eigenvalue is plotted against the wavenumber *k*, where *k = 2π/wavelength*. The real part of the eigenvalue is plotted with a continuous line, while the complex part is plotted with a dotted line. For Hopf and Turing I Hopf, the two highest complex eigenvalues are plotted to show the complex conjugate feature.

#### 4.5 Patterns beyond Turing: non-predictive linear stability analysis

Throughout this study, linear stability analysis is used as a method for studying the patterning probability of our system. However, it is important to understand that under certain conditions classical linear stability analysis might underestimate the size of the patterning parameter space.

Previous studies have reported growth as a way of increasing Turing pattern robustness, as they observe growth induced instabilities[17]. However, our linear stability analysis does not account for growth and might therefore miss this type of instability and report it as a stable system. In Fig. S**26** we see an example of a growth induced instability: the dispersion relation doesn't show any unstable modes; however, the simulation shows a clear periodic heterogeneity. The dominant mode of this dispersion relation (Fig. S**26a**) has a wavenumber of 1.8, which corresponds to a wavelength of *2π/1.8=3.49.* This approximately corresponds to the wavelength of the produced pattern (Fig. S**26b**), meaning this stable mode has been excited to instability and resulted in a Turing pattern.

Other biological effects, such as noise, could also induce patterning. Noise was shown to promote Turing instabilities in certain non-Turing parameter regimes[18–20]. This is specifically for the case of systems with a dominant mode very close to zero. However, we did not study these noise-dependent effects because of our use of a deterministic differential equation model.

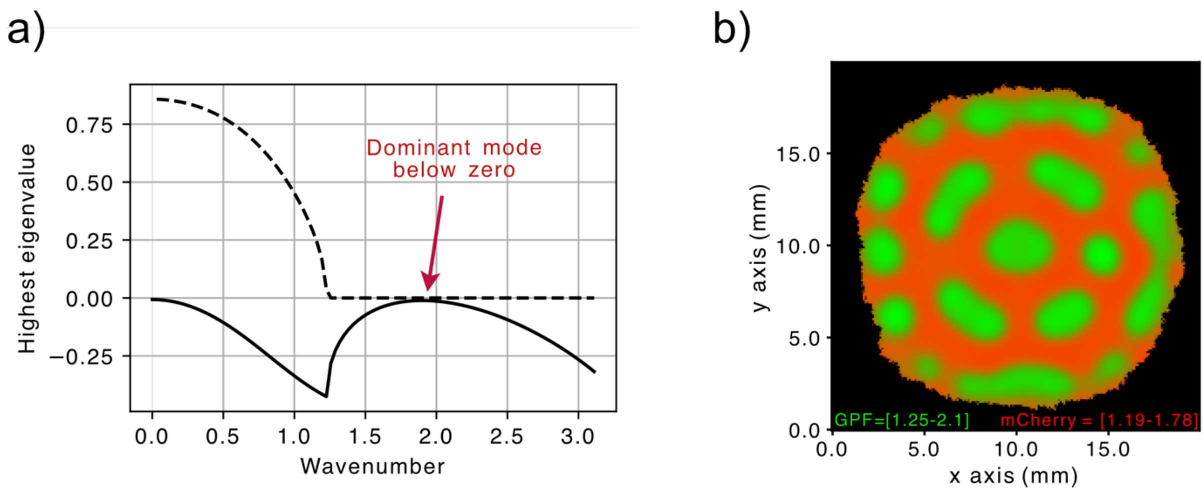

**Supplementary Figure 26: growth induced instability.** (a) Dispersion relation showing a stable system. The most dominant mode (wavenumber=1.8, wavelength=3.49). (b) Using 1% noise (mean μ=p and standard deviation σ=p*0.01), frequency of different analytical solutions. (c) Numerical simulations in growing colonies of 1% noise distribution.

### Supplementary 5: Colony Growth Dynamics

This section describes key aspects of colony growth including experimental growth curves as well as the model to replicate experimental growth, parameters for such model and *in-silico* colony growth outputs.

#### 5.1. Experimental colony growth rate

The diameter of colonies was consistently observed to grow linearly with time – see Fig. S**27** for two examples. The computational simulation of growth was built with a cellular automata algorithm, which is described below, where new cells are added to the edge of the colony depending as a function of the neighbouring pixels and a certain probability of division. The cellular automaton algorithm reproduced the linear growth dynamics observed in the experiments.

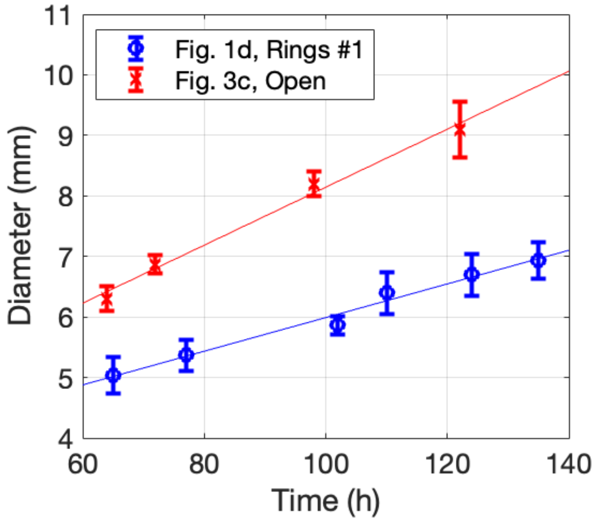

**Supplementary Figure 27: growth rate of two colonies.** Image processing software was used to measure the size of the colony diameter across two different axes, this variability is captured in the error bars (± SEM, n = 2). Two colonies that grow equally in all directions (isotropic growth) were used as examples.

#### 5.2. Cellular automata model

This algorithm is built so that it applies the cellular automata rules to an initial condition matrix. A single colony is modelled by starting with a matrix of 0's (*off*) with a 1 (*on*) in the middle, describing the first cell as it occurs in single cell colonies. In this case J is the number of grid points in the x or the y axis. For two colonies, two 1's are placed at a distance. The division process consists of a probabilistic process where division occurs or does not based on a probability of division ($p_d$). The division process is iteratively applied to the matrix until the final time is reached. The computation is applied every 'm' hours, when $T = \sum_{n=1}^{T/m} m \cdot n$

(e.g. if m=0.2, at T=0.2,0.4,0.6...etc). The growth rate can be tuned by increasing the probability of division ($p_d$) or decreasing m. Furthermore, different $p_d$'s can be applied to different regions of the matrix to achieve faster growing subregions within the colony. The parameters in Table S**7** describe different growth regimes used for simulations. The length of the system is

$$L = J \cdot dx \qquad 35$$

where J corresponds to the corresponds to the number of grid points in the x or y axis, and the matrix has a size of J x J. dx is the length of each grid point. Similarly, time has a length of

$$T = N \cdot dt \qquad 36$$

where N is the number of time points. The bacterial colonies in Fig. S**28** are generated using the parameters in Table S**7**.

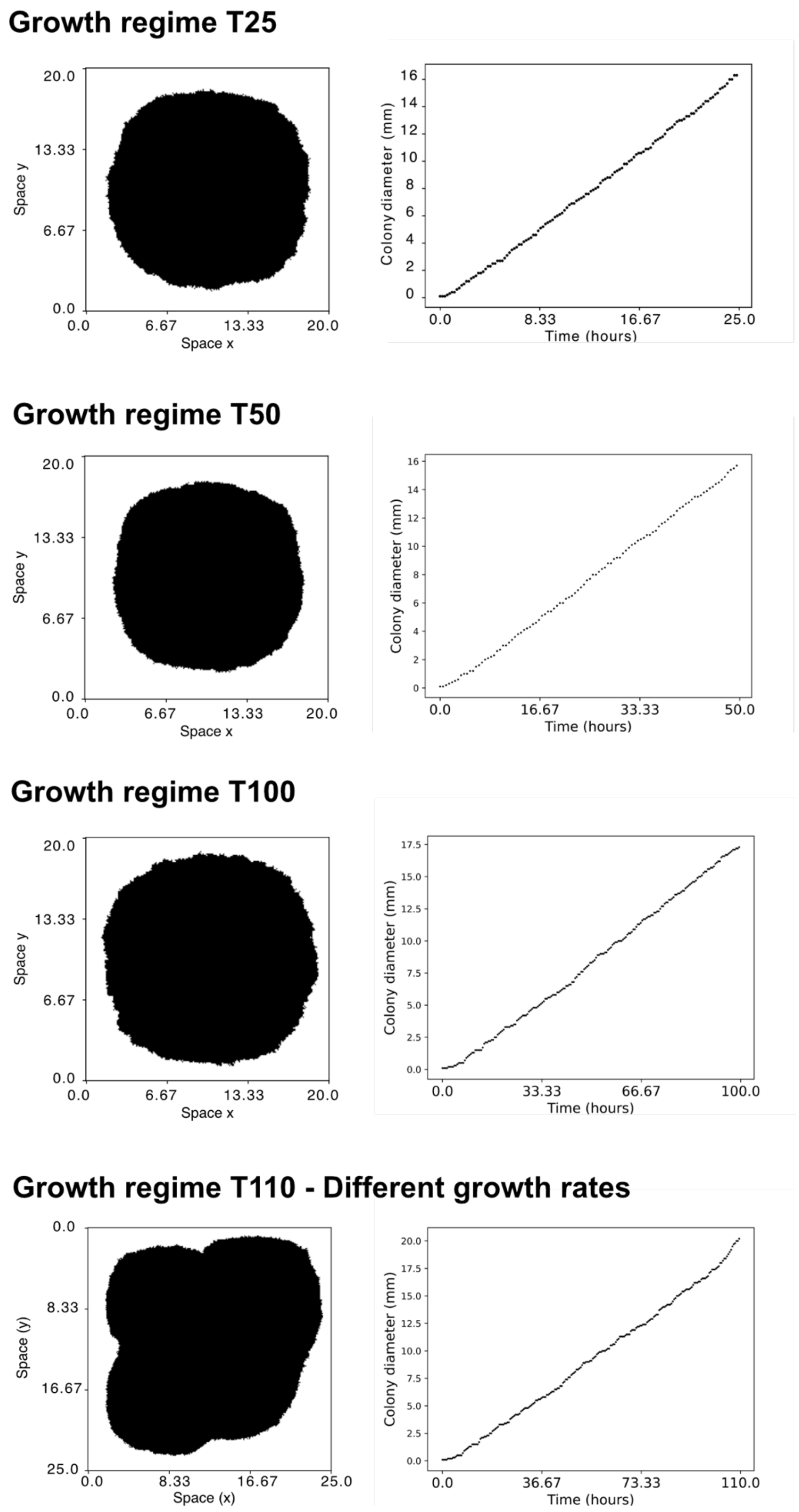

**Supplementary Figure 28: bacterial colony and growth curves from cellular automata simulations.** Using different growth regimes **(left)** final snapshot of bacterial colony and **(right)** growth curve of bacterial colony.

**Supplementary Table 7:** cellular automata and PDE solver parameters for different growth regimes.

| Growth regime | L | T | J | N | dx | dt | boundary leakiness (1 is closed, 2 is open) | shape | Random seed | m | Probability of division ($p_d$) |
|---|---|---|---|---|---|---|---|---|---|---|---|
| T10 | 20 | 10 | 200 | 500 | 0.1 | 0.02 | 1 | cellular automata | 1 | 0.2 | 0.7 |
| T25 | 20 | 25 | 200 | 1250 | 0.1 | 0.02 | 1 | cellular automata | 1 | 0.2 | 0.7 |
| T50 | 20 | 50 | 200 | 2500 | 0.1 | 0.02 | 1 | cellular automata | 1 | 0.5 | 1 |
| T50 Open boundary | 20 | 50 | 200 | 2500 | 0.1 | 0.02 | 2 | cellular automata | 1 | 0.5 | 1 |
| T100 | 20 | 100 | 200 | 5000 | 0.1 | 0.02 | 1 | cellular automata | 1 | 0.5 | 0.38 |
| T110 Different growth rates | 25 | 110 | 250 | 5500 | 0.1 | 0.02 | 1 | cellular automata with different growth rates | 1 | 0.5 | 0.38 and 0.91 |

The state of the growth matrix at every time step is recorded, as well as information on the cell lineage (Fig S23B). This is integrated with the PDE solver (Fig S23A) by computing reaction and diffusion in 1 (*on*) or only diffusion in 0 (*off*) for the matrix configuration at time=T (Fig S23C). When division occurs, the cell lineage information is retrieved to provide the same concentration of molecules from the mother cells to the daughter cells.

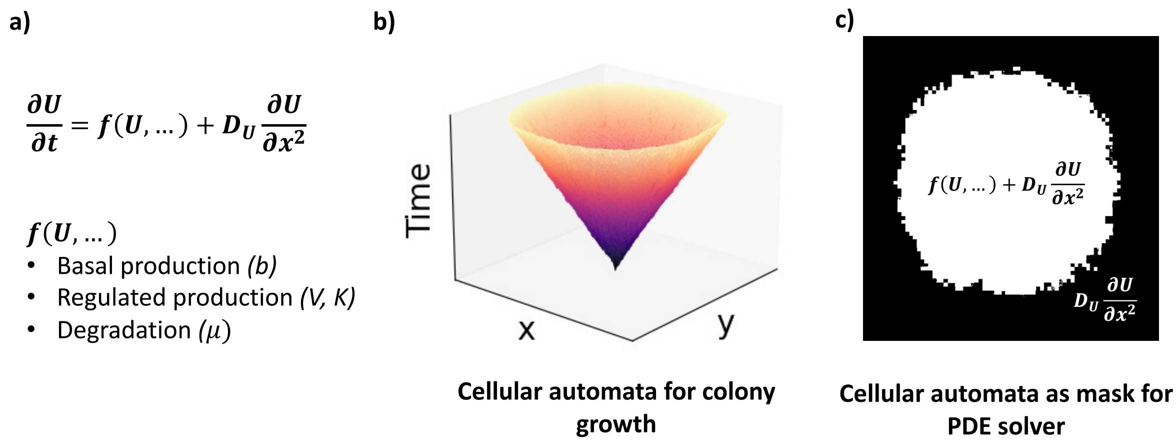

**Supplementary Figure 29: integration of cellular automata and PDE solver.** a) The PDE solver simulates a reaction-diffusion system with temporal and spatial components. b) Cellular automaton simulation in two spatial dimensions (x, y) and time (purple are early timepoints, whereas orange are later timepoints). c) Computation of the PDE solver within the bacterial colony mask. Both reaction and diffusion terms are computed in white pixels, whereas only diffusion terms are computed in black pixels.

#### 5.3. Time and space in the dimensionless model

As seen in Eq. 19, time and space are dimensionless and dependent on the model's parameters

$$t = \frac{t^*}{\mu_a}, x = \sqrt{\frac{k_1 D_u}{\mu_a \mu_u}} x^* \qquad 37$$

In the sampling performed, $\mu_a$ = 0.3 $h^{-1}$, $k_1$ = 0.0183 $h^{-1}$, $\mu_u$ = 0.0225 $h^{-1}$ and $D_U$ is sampled from a range of 0.1-10 $mm^2$ $h^{-1}$ using Eq. 37, time is transformed so that $t^*$=50, t=166h. Space is also transformed but is dependent on $D_U$, which is sampled from a range. Therefore, for $x^*$=16, x = [8.32, 83.2] mm, where with $D_U$ = [0.1, 10] $mm^2$ $h^{-1}$ respectively. These values of t and x lie within realistic physical parameters for our system. These transformations are dependent on parameters that may vary experimentally and therefore some uncertainty must be allowed.

### Supplementary 6: Whole Circuit Sequencing

Whole circuit NanoPore sequencing 6.5 days after seeding the colony of Fig. 1d (Rings #1).

Threshold for SNP: 45%

- Sequencing around strong terminator sequences leads to many artefact SNPs. Increasing the threshold to 45% removes most of these.
- Some high-frequency short tandem repeat SNPs were manually removed because these are often artefacts of the NanoPore sequencing technique.

##### Node A

- Substitution mutation G -> A in promoter J23101 of 4cI-tal, 49% of sequences.
- Substitution mutation G -> T in 4cI gene, 68.5% seq. Silent mutation; CUG -> CUC

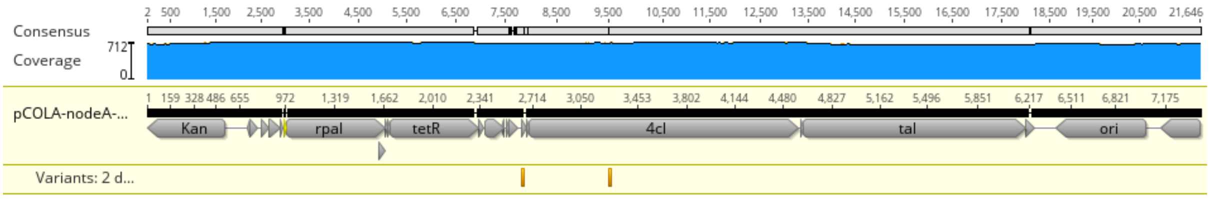

##### Node B

- Point mutation in Spec resistance gene.
- Substitution mutation G -> A in origin of replication may affect copy number. The same mutation is also present in the control, where DNA is transformed into cells, purified straight away and sequenced, therefore it is not a result of adaptation during prolonged culturing.

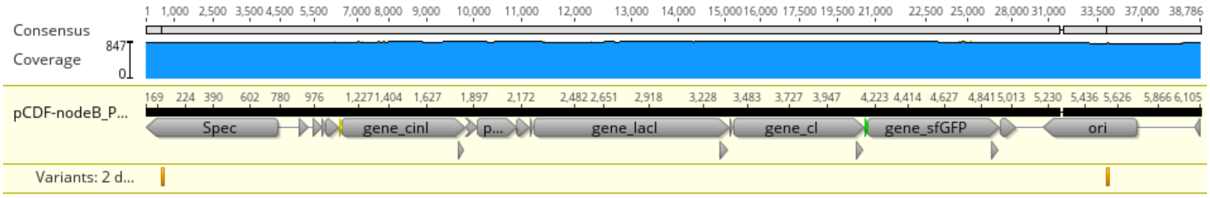

##### Node C

- One mutation in non-coding region.

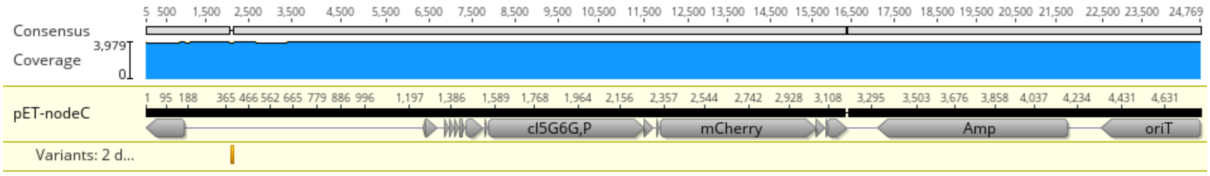

The different regions of the colony were sequenced with NanoPore. The sequences above are for the 'rings' region, towards the edge of the colony. The colony centre and red edge were also sequenced, but sequence alignments showed no additional mutations in any of those regions, other than what is presented above. Overall, no significant functional mutations were found in any of the regions or circuit plasmids.

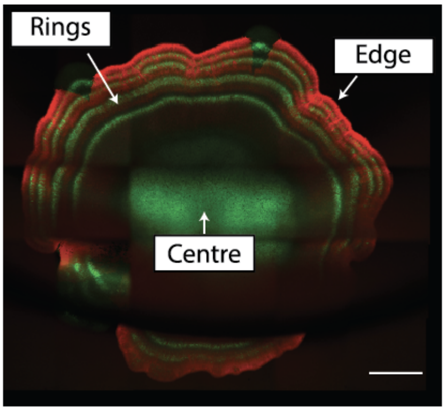

**Supplementary Figure 30: the three sequenced colony regions.**

The number of reads obtained from the NanoPore sequencing can be an indicator for plasmid abundance in the samples. The stripes, the centre, and the red colony regions are relatively consistent amongst each other in terms of plasmid presence and abundance.

The pET copy number is around 4-fold greater compared to pCOLA and pCDF. The copy numbers of pCOLA and pCDF are similar. This is consistent with what is reported in the literature, where copy number is 12, 24 and 38 for pCOLA, pCDF and pET, respectively[21]. The pCC1 copy number is much smaller than for the other plasmids, consistent with the fact that pCC1 is a single-copy BAC (bacterial artificial chromosome)[22].

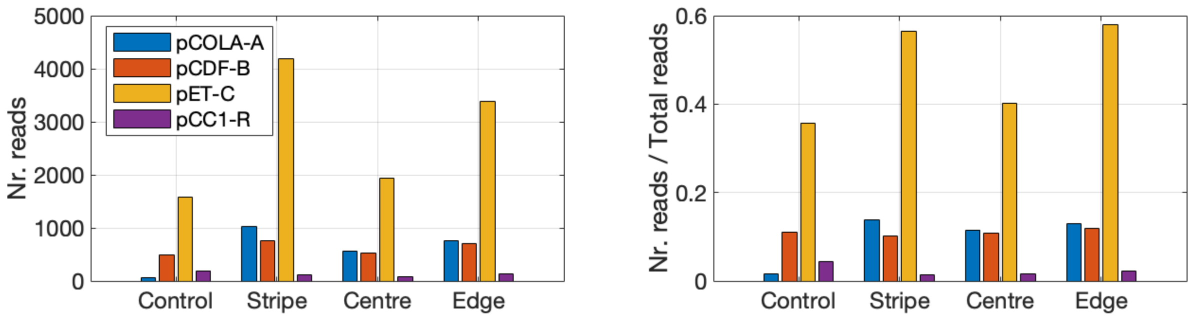

**Supplementary Figure 31: number of reads from NanoPore sequencing.** The sequenced samples are derived from three different regions of the bacterial colony. The control measurement consisted in transforming cells with the circuit plasmids, immediately purifying DNA, and sending for sequencing, without culturing for more than 6 days.

### Supplementary 7: Components and DNA sequences

**Supplementary Table 8: sources of the genetic components that were used.**

| Parts | Category | References |
|---|---|---|
| **Pcin** | Promoter | Meyer et al. (2019)[5]<br>BBa_R0078 |
| **Prpa** | Promoter | Du et al. (2020)[23] |
| **Ptac** | Promoter | Meyer et al. (2019)[5] |
| **PcI*** | Promoter | Brödel et al. (2016)[24] |
| **ASV, AAV, LVA** | Degradation tag | Andersen et al. (1998)[8] |
| **RiboJ, ElvJ** | Ribozyme | Lou et al. (2012)[25,26] |
| **BBa_B0033, BBa_B0064, BBa_B0032, BBa_B0034** | Ribosome Binding Site | iGEM repository |
| **PM-O1-5G6G :: PM-O2-5G6G :: PM-O3-CS** | Promoter/operator | Brödel et al. (2016) [24] |
| **cIO_5G6GP** | Operator | Brödel et al. (2016) [24] |
| **tetO** | Operator | iGEM repository<br>BBa_ K079036 |
| **L3S3P21, L3S2P11, L3S2P21, ECK120033737, rrnB1, IOT** | Terminator | Chen et al. (2013)[27] |
| **BBa_B0015** | Terminator | iGEM repository |
| **pCOLA** | Plasmid backbone | Schaerli et al. (2014)[1] |
| **pCDF** | Plasmid backbone | Schaerli et al. (2014)[1] |
| **pET** | Plasmid backbone | Schaerli et al. (2014)[1] |
| **pCC1** | Plasmid backbone | Proprietary |

**Supplementary Table 9: sources of the genes that were used.** Abbreviations: PCA – protocatechuate, SA – salicylate, Van – vanillin; for other abbreviations see Table S**10**.

| Parts | Description | References |
|---|---|---|
| **rpaI** | Diffuser synthesis enzyme (pC) | Du et al. (2020)[23] |
| **cinI** | Diffuser synthesis enzyme (OC14) | iGEM repository BBa_C0076 |
| **4cl, tal** | Precursor synthesis enzymes (pC) | Du et al. (2020)[23] |
| **tetR, lacI** | Transcription factors | Generic, iGEM repository |
| **cI, $cI_{5G6GP}$** | Transcription factors | Brödel et al. (2016)[24] |
| **$pcaU^{AM}$, $nahR^{AM}$, $phlF^{AM}$, $cinR^{AM}$, $vanR^{AM}$** | Small molecule receptors (PCA, SA, DHBA, OC14, Van, respectively) | Meyer et al. (2019)[5] |
| **rpaR** | Small molecule receptor (pC) | Du et al. (2020)[23] |
| **aiiA** | Lactonase for quorum quenching (pC and OC14 degradation) | iGEM repository BBa_C0160 |

**Supplementary Table 10: chemical inducers.** All were dissolved in DMSO.

| Abbreviation | Full name | Catalogue number |
|---|---|---|
| **OC14** | N-(3-Hydroxytetradecanoyl)-DL-homoserine lactone | Sigma-Aldrich 51481 |
| **pC** | N-(p-Coumaroyl)-L-homoserine lactone | Sigma-Aldrich 07077 |
| **DHBA** | 3,4-dihydroxybenzoic acid | Thermo Scientific 452800010 |
| **ATC** | Anhydrotetracycline | Sigma-Aldrich 37919 |
| **IPTG** | Isopropyl β-D-1-thiogalactopyranoside | Sigma-Aldrich I6758 |

**Supplementary Table 11: full circuit plasmids.** The concentrations next to the antibiotics are in µg/ml. Abbreviations: Kan – kanamycin, Spec – spectinomycin, Amp – ampicillin/carbenicillin, CA – chloramphenicol.

| Plasmid | Resistance | Contains ... | Copy number |
|---|---|---|---|
| **pCOLA** | Kan 50 | Node A | Medium (20 – 40) [21] |
| **pCDF** | Spec 50 | Node B | Medium (20 – 40) [21] |
| **pET** | Amp 100 | Node C | Medium (20 – 40) [21] |
| **pCC1** | CA 10 | Regulator cassette | Single copy [22] |

**Supplementary Table 12: subcircuit architecture.**

| Subcircuit | Plasmids | Description |
|---|---|---|
| **#1** | pCDF-B-ΔcinI | Node B with cinI cassette deleted |
| | pET-C | Node C |
| | HPVoT | p15A ori, constitutive TetR – BBa_J23106::tetR |
| | pCC1R | Regulator plasmid |
| **#2** | pCDF-B-ΔcI | Node B with cI gene deleted |
| | pET-C | Node C |
| | HPVoT | p15A ori, constitutive TetR – BBa_J23106::tetR |
| | pCC1R | Regulator plasmid |
| **#2b** | pCDF-B | Node B |
| | pCC1R | Regulator plasmid |
| **#3** | pCDF-B- ΔcI | Node B with cI gene deleted |
| | pCOLA-A | Node A |
| | pET-C | Node C |
| | pCC1R | Regulator plasmid |

##### 7.1. Legend

Promoter
Operator
RBS
RiboJ
Terminator
Gene

##### 7.2. Node A, plasmid pCOLA-A-pC-tac

Parts sequence

TT L3S3P21 :: Ptac :: riboJ :: RBS BBa_B0033 :: rpaI_ASV :: RBS BBa_B0064 :: tetR_LVA :: TT B0015 :: TT ECK120033737 :: Pc BBa_J23101 :: RBS BBa_B0064 :: 4cl :: RBS BBa_B0032 :: tal :: TT L3S2P11 :: pCOLA ori :: KmR

DNA sequence

ACCGGTAGATCTCCAATTATTGAAGGCCTCCCTAACGGGGGGCCTTTTTTTGTTTCTGGTCTCCCGCT
TAACGATCGTTGGCTGGGTACCtgttgacaattaatcatcggctcgtataatgtgtggaattgtgagc
gctcacaattGGATCCAGCTGTCACCGGATGTGCTTTCCGGTCTGATGAGTCCGTGAGGACGAAACAG
CCTCTACAAATAATTTTGTTTAAgaattcATTTCACCCAAGgagctctcacacaggacTTACTAGATG
CAGGTTCATGTCATCCGTCGAGAGAACCGCGCTCTATGCCGGTCTGCTCGAAAAGTATTTCCGCAT
CCGTCACCAGATCTACGTCGTCGAGCGCGGCTGGAAGGAACTCGATCGGCCGGATGGCCGCGAGATCG
ATCAGTTCGACACCGAAGACGCCGTGTATCTGCTCGGCGTCGACAATGACGACATCGTCGCCGGCATG
CGGATGGTGCCGACCACGTCACCGACGCTCCTCAGCGACGTCTTCCCGCAGCTTGCGCTGGCAGGCCC
GGTGCGGCGGCCGGATGCCTACGAGCTGTCGCGGATCTTCGTGGTACCGCGCAAGCGCGGCGAGCATG
GCGGCCCGCGCGCCGAAGCCGTGATCCAGGCCGCCGCGATGGAGTACGGCCTGTCGATCGGTCTGTCG
GCCTTCACCATCGTGCTGGAAACCTGGTGGCTGCCGCGACTGGTGGACCAGGGCTGGAAGGCAAAGCC
GCTCGGCCTGCCTCAGGACATCAACGGATTCTCGACCACCGCAGTGATCGTCGACGTCGACGACGACG
CCTGGGTCGGCATCTGCAATCGCCGCTCGGTGCCCGGACCCACGCTGGAATGGCGCGGGCTCGAAGCC
ATCCGCCGTCATTCGCTTCCGGAGTTTCAGGTGATTTCAaggcctgctgctaacgatgaaaactacgc
agctagcgttTGAtaaACGCGTTCTAGAGAAAGAGGGGAAATACTAGATGTCTAGATTAGATAAAAGT
AAAGTGATTAACAGCGCATTAGAGCTGCTTAATGAGGTCGGAATCGAAGGTTTAACAACCCGTAAACT
CGCCCAGAAGCTAGGTGTAGAGCAGCCTACATTGTATTGGCATGTAAAAAATAAGCGGGCTTTGCTCG
ACGCCTTAGCCATTGAGATGTTAGATAGGCACCATACTCACTTTTGCCCTTTAGAAGGGGAAAGCTGG
CAAGATTTTTTACGTAATAACGCTAAAAGTTTTAGATGTGCTTTACTAAGTCATCGCGATGGAGCAAA
AGTACATTTAGGTACACGGCCTACAGAAAAACAGTATGAAACTCTCGAAAATCAATTAGCCTTTTTAT
GCCAACAAGGTTTTTCACTAGAGAATGCATTATATGCACTCAGCGCTGTGGGGCATTTTACTTTAGGT
TGCGTATTGGAAGATCAAGAGCATCAAGTCGCTAAAGAAGAAAGGGAAACACCTACTACTGATAGTAT
GCCGCCATTATTACGACAAGCTATCGAATTATTTGATCACCAAGGTGCAGAGCCAGCCTTCTTATTCG
GCCTTGAATTGATCATATGCGGATTAGAAAAACAACTTAAATGTGAAAGTGGGTCCAGGCCTGCTGCT
AACGATGAAAACTATGCTCTGGTCGCCTGATAAccatggccaggcatcaaataaaacgaaaggctcag
tcgaaagactgggcctttcgttttatctgttgtttgtcggtgaacgctctctactagagtcacactgg
ctcaccttcgggtgggcctttctgcgtttataCCTAGGAGcaacccagtcTTAACGATCGTTGGCTGA
CTAGTggaaacacagAAAAAGCCCGCACCTGACAGTGCGGGCTTTTTTTTTcgaccaaaggagtact

CACCCAAGGTTGAGTTACCtttacagctagctcagtcctaggtattatgctagcTCTAGAGAAAGAGG
GGAAATACGAGATGGACGCCATGACCGATCCCATTACATTCGCCGCTACTGCCGCCGGGCGCGACGCG
GCTCATCCGCTGCGCGACATTTCTTTCGGCGAGATAGGTATCGCTACCGAGCGAAAGCCGGACGGCAC
GATCTACGTGCGCTCGACCACCACGCTCACCGACTATCCGGTGCGGATTACCGACCGGCTGCATCACT
TCGCCGAGACGGCGCCCGACCGGGTGTTCATGGCCGAGCGGAACGGTGAGGGCGGCTGGCGCAGGATC
AGCTATGCCGAGATGCTGCGCGCCGCGCAGACCATCGCCTCGGCGCTGATCGCGCGCGGACTGTCGGC
CGAACGGCCGGTGATGATTCTGTCCGGTAATTCGATCGACCATGCGATGGTGATGTTCGGCGCGCTGT
ATGCGGGCGTCGCGATGTGTCCGGTGTCGCCGCCGTATTCGCTGGTGTCCAAGGATTACGGCAAGCTG
CGCCATATCGTCGGGCTGCTGACTCCGGGGCTGATCTTCGCCGATGACATCACTGCCTTCGCGCCCGC
GATCCTCGCCACCGTGCCGGAGGATGTCGAACTGGCTGCCACGCGCGGCGAGGTGAAGGGGCGCAAGG
TGACGTCGCTTGCCGAACTGCTGGCGACGCCGGAACATCCCGAACTCGCCGCCAAGCACGAAGCGATC
GGCCACGACACCATCGCTAAGTTTTTGCTGACGTCGGGATCGACCGGCAATCCGAAGGCGGTGATCAA
TACGCAGCGGATGATCTGCGCCAATCAGGTGATGATCCGCGAGGCGATGGCGTTCCTGAAAGACGAGC
CGCCGGTGATCGTCGACTGGCTGCCGTGGAATCACACCTTCGGCGGCAACCACAATATCGGTCTGACG
CTGTTCAACGGCGGCTCGATGTATATCGACGACGGCAAGCCGACGCCGGCCGGGATCGCTTCCACCAT
CCGCAATCTGCGCGAGATCGCGCCGACGGTGTATTTCAACGTTCCGAAGGGCTACGAGTCGCTGCTGC
CGGTGCTGCGCGAAGACCAGCAGTTGCGCAAATTGTTCTTCAGCCGGCTGCATGCGATGTTCTTCTCC
GGCGCCAGCCTCGCGGCGCATGTCTGGAACGGGCTTGACGAGGTCGCGGTCGCGGAGACCGGCGCGCG
GGTGCCGATGCTCACCGGCCTTGGCGCCACCGAGACCGCGCCGTTCTTCATGTCGGTGACGCCGCAGA
CCAGTCGCTCCGGCCATGTCGGCCTGCCGGTGCCCGGCAACGAGGCCAAGCTGGTGCCGAACAACGGC
AAGCTCGAAGTCCGCGCCAAGGGGCCGAACATCACCCCCGGCTATTGGCGCGCGCCCGAGCTGACCGA
TAAGGCGTTCGACGAGGAGGGCTTCTACAAGCTCAACGATGCGCTGAAGCCGGTCGATGCCAACGACC
TTTCGCGCGGCTTCGATTTCGACGGCCGGATCTCGGAAGACTTCAAACTGGCGTCGGGCACCTGGGTC
AGCGTCGGTCCGCTACGCGCCAAGTTCATTGCCGCCTGCGCCTCCCTGGTGCGCGACGTGGTGATCGC
CGGGCTCGACCGCGATTACGTCACCGCGCTGGCGATCCTCGATCCCGACGGCTGCAAGCTGATCAATG
CGACGCTGCCGCTGGAAGACCTCGCCGGCATGGCGGCCGACCATCTGATCCGCGAGGCGTTCCGCGAG
CGCTTCGCCACGCTGCTGACGCAGGCGACCGGCTCGTCCAACCGCGTCACCCGCGCCGTGCTGCTCGG
CGAACCGCTGTCGATCGACAAGGGGGAGATCACCGACAAGGGCTCGGTCAACCAGCGGGCCGTGCTGG
AATATCGCGCCTCGTTGATCGCGGATCTTTACGCCGACCCACCGCCCGCGCATGTGATTGCGCTGGAG
TGAATTAAAGAGGAGAAAtcacacaggaaagGGTACCATGCTGGCCATGAGCCCGCCGAAACCGGCAG
TGGAACTGGATCGTCATATTGATCTGGACCAGGCACACGCTGTTGCAAGTGGCGGTGCACGTATCGTC
CTGGCACCGCCGGCCCGTGATCGCTGCCGTGCATCCGAAGCTCGTCTGGGTGCAGTCATTCGTGAAGC
TCGCCATGTGTATGGCCTGACCACGGGTTTTGGTCCGCTGGCAAACCGTCTGATCAGCGGTGAAAATG
TGCGCACCCTGCAAGCTAACCTGGTTCATCACCTGGCCTCTGGCGTGGGTCCGGTTCTGGATTGGACC
ACGGCACGTGCAATGGTCCTGGCACGCCTGGTGAGCATTGCACAAGGTGCCAGTGGTGCATCCGAAGG
CACCATTGCGCGTCTGATCGATCTGCTGAACAGCGAACTGGCTCCGGCGGTGCCGAGCCGCGGTACCG
TTGGTGCATCTGGCGACCTGACGCCGCTGGCACATATGGTGCTGTGTCTGCAAGGCCGCGGTGATTTC
CTGGATCGTGACGGTACCCGCCTGGACGGTGCAGAAGGCCTGCGTCGCGGCCGTCTGCAACCGCTGGA
TCTGAGTCACCGCGACGCCCTGGCACTGGTGAACGGTACCAGCGCGATGACGGGCATCGCCCTGGTTA
ATGCTCATGCGTGCCGTCACCTGGGTAACTGGGCAGTTGCACTGACCGCTCTGCTGGCAGAATGTCTG
CGTGGTCGTACGGAAGCATGGGCGGCCGCACTGAGTGATCTGCGCCCGCATCCGGGTCAGAAAGACGC
TGCGGCCCGCCTGCGTGCACGTGTTGATGGCTCCGCTCGTGTGGTTCGCCACGTCATTGCGGAACGTC
GCCTGGATGCCGGTGACATCGGCACCGAACCGGAAGCCGGTCAGGATGCATACAGTCTGCGTTGCGCA
CCGCAAGTGCTGGGTGCTGGTTTTGATACCCTGGCGTGGCATGACCGCGTTCTGACCATTGAACTGAA
CGCGGTTACGGATAATCCGGTCTTTCCGCCGGACGGTTCAGTCCCGGCACTGCATGGCGGTAATTTCA
TGGGCCAGCACGTTGCACTGACCTCGGATGCTCTGGCAACCGCCGTTACGGTCCTGGCAGGTCTGGCA

GAACGTCAAATCGCGCGCCTGACGGACGAACGTCTGAATCGCGGCCTGCCGCCGTTTCTGCATCGTGG
TCCGGCGGGCCTGAACAGCGGTTTCATGGGTGCACAGGTGACCGCAACGGCTCTGCTGGCTGAAATGC
GTGCAACCGGTCCGGCATCAATTCACAGCATCTCTACGAACGCAGCTAATCAAGATGTCGTGTCGCTG
GGCACCATTGCGGCCCGTCTGTGTCGCGAAAAAATCGACCGTTGGGCAGAAATTCTGGCTATCCTGGC
ACTGTGCCTGGCACAGGCAGCTGAACTGCGCTGTGGCTCAGGTCTGGATGGTGTCTCGCCGGCAGGCA
AAAAACTGGTGCAGGCGCTGCGTGAACAATTCCCGCCGCTGGAAACCGATCGTCCGCTGGGTCAGGAA
ATTGCAGCACTGGCGACGCATCTGCTGCAACAATCTCCGGTGTAACTCGGTACCAAATTCCAGAAAAG
AGACGCTTTCGAGCGTCTTTTTTCGTTTTGGTCCCTGCAGGACCTCAGCAATCGAGcaacccagtcag
ctccttccggtgggcgcggggcatgactaacatgagaattacaacttatatcgtatggggctgacttc
aggtgctacatttgaagagataaattgcactgaaatctagagtgatggtgtcgggaatccgtaaagga
tcttcttgagatccttttacgatcgtcgtaatctcctgctctgtaaacgaaaaaaccgcctggggagg
cggtttgatcgaaggttaagtcagttggggaactgcttaacctggtaactggctttagtggagcgcag
ataccaaatactgtcctttcagtgtagcctctgttaggccaccacttcaagactctcgatatctaaat
ccactaattctcagttaccaatggctgctgccagtggcgttttgtcgtgtctttccgggttggactca
agatgatagttaccggataaggcgcagcagtcggctgaacggggggttcttgcacacagcccagctt
ggagcgaactgtctacacggaacggacgtggtgatttgggtaaagcctccaccacaacacggacgcc
gcaggacgggaacaggagagcgcaagagggagccatcaggggaaacgcctggtatctttatagtcct
gtcgggtttcgccaccactgatttgagcgtcagatttcgtgatgttcgtcaggggggcggagcctatg
gaaaaacggcttcgctccggccttattgtctctctgctaagtatcctcctggcatcttctaggacgtt
tctgcgctagcatgcctatttgtttatttttctaaatacattcaaatatgtatccgctcatgagacaa
taaccctgataaatgcttcaataatattgaaaaaggaagagtatgagccatattcaacgggaaacgtc
ttgctctaggccgcgattaaattccaacatggatgctgatttatatgggtataaatgggctcgcgata
atgtcgggcaatcaggtgcgacaatctatcgattgtatgggaagcccgatgcgccagagttgtttctg
aaacatggcaaaggtagcgttgccaatgatgttacagatgagatggtcagactaaactggctgacgga
atttatgcctcttccgaccatcaagcattttatccgtactcctgatgatgcatggttactcaccactg
cgatccccgggaaaacagcattccaggtattagaagaatatcctgattcaggtgaaaatattgttgat
gcgctggcagtgttcctgcgccggttgcattcgattcctgtttgtaattgtccttttaacagcgaccg
cgtatttcgtctcgctcaggcgcaatcacgaatgaataacggtttggttgatgcgagtgattttgatg
acgagcgtaatggctggcctgttgaacaagtctggaaagaaatgcataaacttttgccattctcaccg
gattcagtcgtcactcatggtgatttctcacttgataaccttatttttgacgaggggaaattaatagg
ttgtattgatgttggacgagtcggaatcgcagaccgataccaggatcttgccatcctatggaactgcc
tcggtgagttttctccttcattacagaaacggctttttcaaaaatatggtattgataatcctgatatg
aataaattgcagtttcatttgatgctcgatgagtttttctaagaattaattcatgagcggatacatat
ttgaatgtatttagaaaaataaacaaataggggttccgcgcacatttccccgaaaagtgccacttgcg
gagacccggtcgtcagcttgtcgtcggttcagggcagggtcgttaaatagccgcttatgtctattgct
ggttt

##### 7.3. Node B, plasmid pCDF-B-pC

Parts sequence

TT L3S3P21 :: Prpa* :: cIO_5G6GP :: elvJ :: RBS BBa_B0033 :: cinI_LVA :: TT L3S2P21 :: Pcin :: riboJ :: RBS BBa_B0064 :: lacI_LVA :: RBS BBa_B0064 :: cI_LVA :: RBS BBa_B0034 :: sfGFP_ASV :: TT rrnB1 :: pCDF ori :: SpecR

DNA sequence

```
TAGATCTCCAATTATTGAAGGCCTCCCTAACGGGGGGCCTTTTTTTGTTTCTGGTCTCCCGCTTAACG
ATCGTTGGCTGGTCGACACCTGTCCGATCGGACAGTTTTACGCAAGAAAATGGTTTGTTACTTTCGAA
TAAATATCGGCGCCGCCGATAAGCCCCATAGGGTGGTGTGTACCACCCCTGATGAGTCCAAAAGGACG
AAATGGGGCCTCTACAAATAATTTTGTTTAAGGATCCtcacacaggacGAATTCATGttcgttatcat
tcaggcacatgagtatcagaaatacgctgccgtactcgaccagatgtttcgtctgcgcaagaaggtct
tcgccgatacgctctgctgggacgttcctgtcatcggcccttacgaacgtgacagctacgattcgctt
gctcccgcctatctcgtctggtgcaacgacagccgcacccgtctttatggcggcatgcgcctgatgcc
gacgaccggcccgacccttctctacgacgtcttccgcgagacgttccctgatgccgccgatcttatcg
cccccggcatctgggaaggcacgcgcatgtgcatcgacgaggaggcgatcgccaaggatttccccgag
atcgacgccggccgcgccttctccatgatgctgctcgcgctttgcgaatgcgcgctcgatcacggcat
ccacacgatgatctccaactacgagccctacctcaagcgcgtctacaagcgcgccggcgccgaggtgg
aagaactcggccgcgcagacggctacggcaaatatcccgtctgctgcggcgccttcgaagtctcggac
cgcgtgctgcgcaagatgcgcgccgccctcggcctcaccctacccctttatgtcaggcacgtgccggc
ccgctcggtcgtgacccaattcctggagatggcagcaACTAGTgctgcaaacgacgaaaactacgctt
tagtagctTGAACTAGTTGAtaaccaattattgCTCGGTACCAAATTCCAGAAAAGAGGCCTCCCGAA
AGGGGGGCCTTTTTTCGTTTTGGTCCCCCTTTGTGCGTCCAAACGGACGCACGGCGCTCTAAAGCGGG
TCGCGATCTTTCAGATTCGCTCCTCGCGCTTTCAGTCTTTGTTTTGGCGCATGTCGTTATCGCAAAAC
CGCTGCACACTTTTGCGCGACATGCTCTGATCCCCCTCATCTGGGGGGGCCTATCTGAGGGAATTTCC
GATCCGGCTCGCCTGAACCATTCTGCTTTCCACGAACTTGAAAACGCTACGCGTAGCTGTCACCGGAT
GTGCTTTCCGGTCTGATGAGTCCGTGAGGACGAAACAGCCTCTACAAATAATTTTGTTTAATCTAGAG
AAAGAGGGGAAATACTAGATGAAACCAGTAACGTTATACGATGTCGCAGAGTATGCCGGTGTCTCTTA
TCAGACCGTTTCCCGCGTGGTGAACCAGGCCAGCCACGTTTCTGCGAAAACGCGGGAAAAAGTGGAAG
CGGCGATGGCGGAGCTGAATTACATTCCCAACCGCGTGGCACAACAACTGGCGGGCAAACAGTCGTTG
CTGATTGGCGTTGCCACCTCCAGTCTGGCCCTGCACGCGCCGTCGCAAATTGTCGCGGCGATTAAATC
TCGCGCCGATCAACTGGGTGCCAGCGTGGTGGTGTCGATGGTAGAACGAAGCGGCGTCGAAGCCTGTA
AAGCGGCGGTGCACAATCTTCTCGCGCAACGCGTCAGTGGGCTGATCATTAACTATCCGCTGGATGAC
CAGGATGCCATTGCTGTGGAAGCTGCCTGCACTAATGTTCCGGCGTTATTTCTTGATGTCTCTGACCA
GACACCCATCAACAGTATTATTTTCTCCCATGAAGACGGTACGCGACTGGGCGTGGAGCATCTGGTCG
CATTGGGTCACCAGCAAATCGCGCTGTTAGCGGGCCCATTAAGTTCTGTCTCGGCGCGTCTGCGTCTG
GCTGGCTGGCATAAATATCTCACTCGCAATCAAATTCAGCCGATAGCGGAACGGGAAGGCGACTGGAG
TGCCATGTCCGGTTTTCAACAAACCATGCAAATGCTGAATGAGGGCATCGTTCCCACTGCGATGCTGG
TTGCCAACGATCAGATGGCGCTGGGCGCAATGCGCGCCATTACCGAGTCCGGGCTGCGCGTTGGTGCG
GACATCTCGGTAGTGGGATACGACGATACCGAAGACAGCTCATGTTATATCCCGCCGTTAACCACCAT
CAAACAGGATTTTCGCCTGCTGGGGCAAACCAGCGTGGACCGCTTGCTGCAACTCTCTCAGGGCCAGG
CGGTGAAGGGCAATCAGCTGTTGCCCGTCTCACTGGTGAAAAGAAAAACCACCCTGGCGCCCAATACG
CAAACCGCCTCTCCCCGCGCGTTGGCCGATTCATTAATGCAGCTGGCACGACAGGTTTCCCGACTGGA
AAGCGGGCAGAGGCCTGCTGCTAACGATGAAAACTATGCTCTGGTCGCCTGATAAccatggTCTAGAG
```

AAAGAGGGGAAATACTAGATGAGCACAAAAAAGAAACCATTAACACAAGAGCAGCTTGAGGACGCACG
TCGCCTTAAAGCAATTTATGAAAAAAAGAAAAATGAACTTGGCTTATCCCAGGAATCTGTCGCAGACA
AGATGGGGATGGGGCAGTCAGGCGTTGGTGCTTTATTTAATGGCATCAATGCATTAAATGCTTATAAC
GCCGCATTGCTTGCAAAAATTCTCAAAGTTAGCGTTGAAGAATTTAGCCCTTCAATCGCCAGAGAAAT
CTACGAGATGTATGAAGCGGTTAGTATGCAGCCGTCACTTAGAAGTGAGTATGAGTACCCTGTTTTTT
CTCATGTTCAGGCAGGGATGTTCTCACCTGAGCTTAGAACCTTTACCAAAGGTGATGCGGAGAGATGG
GTAAGCACAACCAAAAAAGCCAGTGATTCTGCATTCTGGCTTGAGGTTGAAGGTAATTCCATGACCGC
ACCAACAGGCTCCAAGCCAAGCTTTCCTGACGGAATGTTAATTCTCGTTGACCCTGAGCAGGCTGTTG
AGCCAGGTGATTTCTGCATAGCCAGACTTGGGGGTGATGAGTTTACCTTCAAGAAACTGATCAGGGAT
AGCGGTCAGGTGTTTTTACAACCACTAAACCCACAGTACCCAATGATCCCATGCAATGAGAGTTGTTC
CGTTGTGGGGAAAGTTATCGCTAGTCAGTGGCCTGAAGAGACGTTTGGCAGGCCTGCTGCTAACGATG
AAAACTATGCTCTGGTCGCCTGATAATCTAGAGAAAGAGGAGAAATACTAGATGCGTAAAGGCGAAGA
ACTGTTTACCGGTGTGGTTCCGATTCTGGTGGAACTGGACGGCGATGTTAATGGTCATAAATTCAGTG
TTCGCGGCGAAGGTGAAGGCGATGCGACGAACGGCAAACTGACCCTGAAATTTATCTGCACCACGGGT
AAACTGCCGGTCCCGTGGCCGACGCTGGTGACCACGCTGACCTATGGCGTTCAATGTTTTGCGCGTTA
CCCGGATCACATGAAACAGCACGACTTTTTCAAATCGGCCATGCCGGAAGGCTATGTGCAGGAACGTA
CGATTAGCTTTAAAGACGATGGTACGTATAAAACCCGCGCGGAAGTGAAATTCGAAGGCGATACCCTG
GTTAACCGTATCGAACTGAAAGGTATCGATTTCAAAGAAGACGGCAATATTCTGGGTCATAAACTGGA
ATATAACTTCAATTCCCACAACGTGTACATCACCGCGGATAAACAGAAAAACGGCATTAAAGCCAATT
TCAAAATCCGCCATAATGTGGAAGATGGTAGCGTTCAGCTGGCCGACCACTATCAGCAAAACACGCCG
ATTGGTGATGGCCCGGTCCTGCTGCCGGACAATCACTACCTGAGTACCCAGTCCGTGCTGTCAAAAGA
TCCGAACGAAAAACGTGACCACATGGTCCTGCTGGAATTTGTGACGGCTGCGGGTATCACCCACGGCA
TGGACGAACTGTATAAAAGGCCTGCTGCTAACGATGAAAACTACGCAgctAGCgttTGATAATAGAGG
CATCAAATAAAACGAAAGGCTCAGTCGAAAGACTGGGCCTTTCGTTTTATCTGTTGTTTGTCGGTGAA
CGCTCTCCTGAGTAGGACAAATCCCTCGAGTCACTGCATCCTAGGAGcaacccagtcagctccttccg
gtgggcgcggggcatgactaacatgagaattacaacttatatcgtatggggctgacttcaggtgctac
atttgaagagataaattgcactgaaatctagagcggttcagtagaaaagatcaaaggatcttcttgag
atcctttttttctgcgcgtaatcttttgccctgtaaacgaaaaaaccacctggggaggtggtttgatc
gaaggttaagtcagttggggaactgcttaaccgtggtaactggctttcgcagagcacagcaaccaaat
ctgtccttccagtgtagccggactttggcgcacacttcaagagcaaccgcgtgtttagctaaacaaat
cctctgcgaactcccagttaccaatggctgctgccagtggcgttttaccgtgcttttccgggttggac
tcaagtgaacagttaccggataaggcgcagcagtcgggctgaacggggagttcttgcttacagcccag
cttggagcgaacgacctacaccgagccgagataccagtgtgtgagctatgagaaagcgccacacttcc
cgtaagggagaaaggcggaacaggtatccggtaaacggcagggtcggaacaggagagcgcaagaggga
gcgacccgccggaaacggtggggatctttaagtcctgtcgggtttcgcccgtactgtcagattcatgg
ttgagcctcacggctcccacagatgcaccggaaaagcgtctgtttatgtgaactctggcaggagggcg
gagcctatggaaaaacgccaccggcgcggccctgctgttttgcctcacatgttagtcccctgcttatc
cacggaatctgtgggtaactttgtatgtgtccgcagcgcccgccgcagtctcacgcccggagcgtagc
gaccgagtgagctagctatttgtttatttttctaaatacattcaaatatgtatccgctcatgagacaa
taaccctgataaatgcttcaataatattgaaaaaggaagagtatgagggaagcggtgatcgccgaagt
atcgactcaactatcagaggtagttggcgtcatcgagcgccatctcgaaccgacgttgctggccgtac
atttgtacggctccgcagtggatggcggcctgaagccacacagtgatattgatttgctggttacggtg
accgtaaggcttgatgaaacaacgcggcgagctttgatcaacgaccttttggaaacttcggcttcccc
tggagagagcgagattctccgcgctgtagaagtcaccattgttgtgcacgacgacatcattccgtggc
gttatccagctaagcgcgaactgcaatttggagaatggcagcgcaatgacattcttgcaggtatcttc
gagccagccacgatcgacattgatctggctatcttgctgacaaaagcaagagaacatagcgttgcctt

ggtaggtccagcggcggaggaactctttgatccggttcctgaacaggatctatttgaggcgctaaatg
aaaccttaacgctatggaactcgccgcccgactgggctggcgatgagcgaaatgtagtgcttacgttg
tcccgcatttggtacagcgcagtaaccggcaaaatcgcgccgaaggatgtcgctgccgactgggcaat
ggagcgcctgccggcccagtatcagcccgtcatacttgaagctagacaggcttatcttggacaagaag
aagatcgcttggcctcgcgcgcagatcagttggaagaatttgtccactacgtgaaaggcgagatcacc
aaggtagtcggcaaataatgtctaacaattcgttcaagccgaggggccgcaagatccggccacgatga
cccggtcgtcggttcagggcagggtcgttaaatagccgcttatgtctattgctggtttaccgg

#### 7.4. Node C, plasmid pET-C

Parts sequence

TT L3S3P21 :: PM-O1-5G6G :: PM-O2-5G6G :: PM-O3-CS :: tetO :: riboJ :: RBS BBa_B0064 :: cI-5G6GP_LVA :: RBS BBa_B0034 :: mCherry_ASV :: TT rrnB1 :: AmpR :: pET ori

DNA sequence

gctcgcgtatcggtgattcattctgctaaccagtaaggcaaccccgccagcctagccgggtcctcaac
gacaggagcacgatcatgcgcacccgtggccaggacccaacgctgcccgagatctCCAATTATTGAAG
GCCTCCCTAACGGGGGGCCTTTTTTTGTTTCTGGTCTCCCGCTTAACGATCGTTGGCTGGGTACCGCA
ACCATTATCGGCGCCGCCGATAAAATAGTCAACGGCGGCGCCGATAGATATTTATCACCGCCGGTGAT
AGATTTAACGTATCCCTATCAGTGATAGAGAGAATTCAGCTGTCACCGGATGTGCTTTCCGGTCTGAT
GAGTCCGTGAGGACGAAACAGCCTCTACAAATAATTTTGTTTAATCTAGAGAAAGAGGGGAAATACTA
GATGAGCACAAAAAAGAAACCATTAACACAAGAGCAGCTTGAGGACGCACGTCGCCTTAAAGCAATTT
ATGAAAAAAAGAAAAATGAACTTGGCTTATCCCAGGAATTGGTCGCATACGAGATGGGGATGGGGCAG
TCCGCGGTTTCCGAGTTATTTAATGGCATCTGGGCATTAAATGCTTATAACGCCGCATTGCTTGCAAA
AATTCTCAAAGTTAGCGTTGAAGAATTTAGCCCTTCAATCGCCAGAGAAATCTACGAGATGTATGAAG
CGGTTAGTATGCAGCCGTCACTTAGAAGTGAGTATGAGTACCCTGTTTTTTCTCATGTTCAGGCAGGG
ATGTTCTCACCTGAGCTTAGAACCTTTACCAAAGGTGATGCGGAGAGATGGGTAAGCACAACCAAAAA
AGCCAGTGATTCTGCATTCTGGCTTGAGGTTGAAGGTAATTCCATGACCGCACCAACAGGCTCCAAGC
CAAGCTTTCCTGACGGAATGTTAATTCTCGTTGACCCTGAGCAGGCTGTTGAGCCAGGTGATTTCTGC
ATAGCCAGACTTGGGGGTGATGAGTTTACCTTCAAGAAACTGATCAGGGATAGCGGTCAGGTGTTTTT
ACAACCACTAAACCCACAGTACCCAATGATCCCATGCAATGAGAGTTGTTCCGTTGTGGGAAAGTTA
TCGCTAGTCAGTGGCCTGAAGAAACGTTTGGCAGGCCTGCTGCTAACGATGAAAACTACGCACTGGTG
GCTTGATAATCTAGAGAAAGAGGAGAAATACTAGATGGTGAGCAAGGGCGAGGAGGATAACATGGCTA
TCATCAAGGAGTTCATGCGCTTCAAGGTGCACATGGAGGGCTCCGTGAACGGCCACGAGTTCGAGATC
GAGGGCGAGGGCGAGGGCCGCCCCTACGAGGGCACCCAGACCGCCAAGCTGAAGGTGACCAAGGGTGG
CCCCCTGCCCTTCGCCTGGGACATCCTGTCCCCTCAGTTCATGTACGGCTCCAAGGCCTACGTGAAGC
ACCCCGCCGACATCCCCGACTACTTGAAGCTGTCCTTCCCCGAGGGCTTCAAGTGGGAGCGCGTGATG
AACTTCGAGGACGGCGGCGTGGTGACCGTGACCCAGGACTCCTCCCTGCAGGACGGCGAGTTCATCTA
CAAGGTGAAGCTGCGCGGCACCAACTTCCCCTCCGACGGCCCCGTAATGCAGAAGAAGACTATGGGCT
GGGAGGCCTCCTCCGAGCGGATGTACCCCGAGGACGGCGCCCTGAAGGGCGAGATCAAGCAGAGGCTG
AAGCTGAAGGACGGCGGCCACTACGACGCTGAGGTCAAGACCACCTACAAGGCCAAGAAGCCCGTGCA
GCTGCCCGGCGCCTACAACGTCAACATCAAGTTGGACATCACCTCCCACAACGAGGACTACACCATCG
TGGAACAGTACGAACGCGCCGAGGGCCGCCACTCCACCGGCGGCATGGACGAGCTGTACAAGAGGCCT
GCTGCTAACGATGAAAACTACGCAgctAGCgttTGATAATAGAGGCATCAAATAAAACGAAAGGCTCA
GTCGAAAGACTGGGCCTTTCGTTTTATCTGTTGTTTGTCGGTGAACGCTCTCCTGAGTAGGACAAATC
CCTCGAGgacgtcaggtggcacttttcggggaaatgtgcgcggaacccctatttgtttatttttctaa
atacattcaaatatgtatccgctcatgagacaataaccctgataaatgcttcaataatattgaaaaag
gaagagtatgagtattcaacatttccgtgtcgcccttattcccttttttgcggcattttgccttcctg
tttttgctcacccagaaacgctggtgaaagtaaaagatgctgaagatcagttgggtgcacgagtgggt
tacatcgaactggatctcaacagcggtaagatccttgagagttttcgccccgaagaacgttttccaat
gatgagcacttttaaagttctgctatgtggcgcggtattatcccgtgttgacgccgggcaagagcaac
tcggtcgccgcatacactattctcagaatgacttggttgagtactcaccagtcacagaaaagcatctt
acggatggcatgacagtaagagaattatgcagtgctgccataaccatgagtgataacactgcggccaa

cttacttctgacaacgatcggaggaccgaaggagctaaccgcttttttgcacaacatgggggatcatg
taactcgccttgatcgttgggaaccggagctgaatgaagccataccaaacgacgagcgtgacaccacg
atgcctgcagcaatggcaacaacgttgcgcaaactattaactggcgaactacttactctagcttcccg
gcaacaattaatagactggatggaggcggataaagttgcaggaccacttctgcgctcggcccttccgg
ctggctggtttattgctgataaatctggagccggtgagcgtgggtctcgcggtatcattgcagcactg
gggccagatggtaagccctcccgtatcgtagttatctacacgacggggagtcaggcaactatggatga
acgaaatagacagatcgctgagataggtgcctcactgattaagcattggtaactgtcagaccaagttt
actcatatatactttagattgatttaaaacttcatttttaatttaaaaggatctaggtgaagatcctt
tttgataatctcatgaccaaaatcccttaacgtgagttttcgttccactgagcgtcagaccccgtaga
aaagatcaaaggatcttcttgagatcctttttttctgcgcgtaatctgctgcttgcaaacaaaaaaac
caccgctaccagcggtggtttgtttgccggatcaagagctaccaactctttttccgaaggtaactggc
ttcagcagagcgcagataccaaatactgtccttctagtgtagccgtagttaggccaccacttcaagaa
ctctgtagcaccgcctacatacctcgctctgctaatcctgttaccagtggctgctgccagtggcgata
agtcgtgtcttaccgggttggactcaagacgatagttaccggataaggcgcagcggtcgggctgaacg
gggggttcgtgcacacagcccagcttggagcgaacgacctacaccgaactgagatacctacagcgtga
gctatgagaaagcgccacgcttcccgaagggagaaaggcggacaggtatccggtaagcggcagggtcg
gaacaggagagcgcacgagggagcttccagggggaaacgcctggtatctttatagtcctgtcgggttt
cgccacctctgacttgagcgtcgatttttgtgatgctcgtcaggggggcggagcctatggaaaaacgc
cagcaacgcggcctttttacggttcctggccttttgctggccttttgctcacatgttctttcctgcgt
tatcccctgattctgtggataaccgtattaccgcctttgagtgagctgataccgctcgccgcagccga
acgaccgagcgcagcgagtcagtgagcgaggaagcggaagagcgcctgatgcggtattttctccttac
gcatctgtgcggtatttcacaccgcatatatggtgcactctcagtacaatctgctctgatgccgcata
gttaagccagtatacactccgctatcgctacgtgactgggtcatggctgcgccccgacacccgccaac
acccgctgacgcgccctgacgggcttgtctgctcccggcatccgcttacagacaagctgtgaccgtct
ccgggagctgcatgtgtcagaggttttcaccgtcatcaccgaaacgcgcgaggcagctgcggtaaagc
tcatcagcgtggtcgtgaagcgattcacagatgtctgcctgttcatccgcgtccagctcgttgagttt
ctccagaagcgttaatgtctggcttctgataaagcgggccatgttaagggcggttttttcctgtttgg
tcactgatgcctccgtgtaagggggatttctgttcatgggggtaatgataccgatgaaacgagagagg
atgctcacgatacgggttactgatgatgaacatgcccggttactggaacgttgtgagggtaaacaact
ggcggtatggatgcggcgggaccagagaaaaatcactcagggtcaatgccagcgcttcgttaatacag
atgtaggtgttccacagggtagccagcagcatcctgcgatgcagatccggaacataatggtgcagggc
gctgacttccgcgtttccagactttacgaaacacggaaaccgaagaccattcatgttgttgctcaggt
cgcagacgttttgcagcagcagtcgcttcacgttc

##### 7.5. Receptor array and AiiA, plasmid pCC1R

Parts sequence

TT L3S3P21 :: BBa_J23101 :: RBS pca3 :: pcaU$^{AM}$ :: RBS nah3 :: nahR$^{AM}$ :: RBS BBa_B0034 :: rpaR :: TT BBa_B1006 :: BBa_J23119 :: RBS :: phlF$^{AM}$ :: RBS :: cinR$^{AM}$ :: RBS :: vanR$^{AM}$ :: TT L3S2P21 :: PphlF :: riboJ :: RBS BBa_B0064 :: aiiA :: TT IOT :: camR :: repE ori

DNA sequence

CCAATTATTGAAGGCCTCCCTAACGGGGGGCCTTTTTTTGTTTCTGGTCTCCCGCTTAACGATCGTTG
GCTGGAGATTTTGAGGGTCGAATtttacagctagctcagtcctaggtattatgctagctcatgaCGCT
TACAATAGACGAACAATAAAGGAGGAATTAACCGATGTGGTCGAACATGGATGACAAGAAAGTGAAAG
AGGAGAATATTCTGCACAATTCCACCAACAAGAAGATCATCCGCCACGAAGATTTTGTAGCCGGCATT
AGCAAAGGGATGGCGATTCTGGATTCGTTTGGTACAGATCGTCATCGCCTCAATATCACCATGGCCGC
AGAGAAAACCGGTATGACACGTGCAGCAGCTCGTCGCCACCTGCTTACTCTGGAGTATCTGGGCTATC
TGGAAAGTGACGGCCACTACTTCTACTTAACTCCCAAAATCCTGAAATTCAGTGGTTCATATTTGGGT
GGTGCTCAATTGCCGAAAATTTCCCAACCACTGTTGAACTTGCTTACGACCCAGACCAGCCTGATTTA
CAGCGTGATGGTGTTGGATGGCTATGAAGCCATTACCATTGCGCGTTCTGCCGCTCATCAGCAAACCG
ACCGCGTTAACCCGTATGGTTTACATCTCGGGAATCGCTTACCAGCGCATACAACGTCAGCGGGCAAA
ATCCTGTTAGCGTATTTGGATGACCATGCCCAGCAAGAGTGGCTCAATCAGTACCCTCTGCAACGGCT
CACGAAATACACGTATACCAACCACATCGACTTTCTGCGCCTTTTGAGTGAAATCAAGGAACAGGGTT
GGTGCTATAGTTCGGAAGAACACGAACTGGGAGTACACGCCCTTGCGGTTCCGATTTACGGACAACAG
TCTCGCGTCGTAGCGGCACTGAACATTGTCAGCCCGACAATGCGGACCACGAAAGAATACCTGATTCA
GCATATTCTGCCGTTACTGCAAGAAACTGCGCGTGAATTGCGCAATATCCTGTAATGAACCCCCTATA
AGAAAAAGACTTAACTATCCATGGAACTGCGTGACCTTGATTTAAACCTGCTGGTGGTGTTCAACCAG
TTGCTGGTCGACAGACGCGTCTCTGTCACTGCGGAGAACCTGGGCCTGACCCAGCCTGCCGTGAGCAA
TGCGCTGAAACGCCTGCGCACCTCGCTACAGGACCCACTCTTCGTGCGCACACATCAGGGAATGGAAC
CCACACCCTATGCCGCGCATCTGGCCGAGCACGTCACTTCGGCCATGCACGCACTGCGCAACGCCCTA
CAGCACCATGAAAGCTTCGATCCGCTGACCAGCGAGCGTACCTTCACCCTGGCCATGACCGACATTGG
CGAGATCTACTTCATGCCGCGGCTGATGGATGCGCTGGCTCACCAGGCCCCCAATTGCGTGATCAGTA
CGGTGCGCGACAGTTCGATGAGCCTGATGCAGGCCTTGCAGAACGGAACCGTGGACTTGGCCGTGGGC
CTGCTTCCCAATCTGCAAACTGGCTTCTTTCAGCGCCGGCTGCTCCGTAATCACTACGTGTGCCTATG
TCGCAAGGACCATCCAGTCACCCGCGAACCCCTGACTCTGGAGCGCTTCTGTTCCTACGGCCACGTGC
GTGTCATCGCCGCTGGCACCGGCCACGGCGAGGTGGACACGTACATGACACGGGTCGGCATCCGGCGC
GACATCCGTCTGGAAGTGCCGCACTTCGCCGCCGTTGGCCACATCCTCCAGCGCACCGATCTGCTCGC
CACTGTGCCGATATGTTTAGCCGACTGCTGCGTAGAGCCCTTCGGCCTAAGCGCCTTGCCGCACCCAG
TCGTCTTGCCTGAAATAGCCATCAACATGTTCTGGCATGCGAAGTACCACAAGGACCTAGCCAATATT
TGGTTGCGGCAACTGATGTTTGACCTGTTTACGGATTGATAAGAATTAAAGAGGGGAAAGGTACCATG
ATCGTCGGCGAAGATCAGCTTTGGGGACGGCGTGCGCTGGAGTTCGTCGATTCCGTCGAACGGCTCGA
GGCGCCGGCGCTGATCAGCCGGTTCGAATCGCTGATCGCGAGCTGCGGATTTACCGCCTACATCATGG
CCGGCCTGCCGTCGCGCAATGCCGGACTACCGGAGCTGACGCTGGCCAATGGCTGGCCGCGAGACTGG
TTCGATCTGTATGTCAGCGAAAACTTCAGCGCGGTCGATCCGGTGCCGCGCCACGGCGCTACCACGGT
TCATCCTTTCGTATGGTCCGATGCACCCTACGACCGCGACCGTGATCCGGCCGCCCACCGGGTCATGA
CCCGGGCGGCGGAGTTCGGACTGGTCGAGGGTTACTGCATTCCGCTGCACTACGACGACGGTAGCGCC
GCGATCAGCATGGCCGGCAAAGATCCGGACCTCAGCCCGGCCGCGCGCGGCGCGATGCAGCTGGTCAG
CATCTACGCGCATAGTCGCCTGCGCGCACTCAGCCGGCCAAAGCCGATCCGGCGCAACCGGCTCACGC

CGCGCGAGTGCGAGATCCTGCAATGGGCAGCGCAGGGCAAGACCGCCTGGGAAATCTCGGTAATCCTC
TGCATCACCGAACGCACGGTGAAATTCCATCTGATCGAAGCCGCCCGCAAGCTCGACGCCGCCAACCG
CACCGCGGCGGTTGCCAAGGCATTGACGCTCGGATTGATCCGTTTGTGAAATTCaaaaaaaaaccccg
cccctgacagggcggggttttttTGAGATTTTGAGACACAAGGTCGAAtCGCACCAAGACAGGTTTG
TCCATTGACAGCTAGCTCAGTCCTAGGTATAATGCTAGCCTATGGACTATGTTTGAAAGGGAGAAATA
CTAGATGGCACGTACCCCGAGCCGTAGCAGCATTGGTAGCCTGCGTAGTCCGCATACCCATAAAGCAA
TTCTGACCAGCACCATTGAAATCCTGAAAGAATGTGGTTATAGCGGTCTGAGCATTGAAAGCGTGGCA
CGTCGCGCCGGTGCAGGCAAACCGACCATTTATCGTTGGTGGACCAACAAAGCAGCACTGATTGCCGA
AGTGTATGAAAATGAAATCGAACAGGTACGTAAATTTCCGGATTTGGGTAGCTTTAAAGCCGATCTGG
ATTTTCTGCTGCATAATCTGTGGAAAGTTTGGCGTGAAACCATTTGTGGTGAAGCATTTCGTTGTGTT
ATTGCAGAAGCACAGTTGGACCCTGTAACCCTGACCCAACTGAAAGATCAGTTTATGGAACGTCGTCG
TGAGATACCGAAAAAACTGGTTGAAGATGCCATTAGCAATGGTGAACTGCCGAAAGATATCAATCGTG
AACTGCTGCTGGATATGATTTTTGGTTTTTGTTGGTATCGCCTGCTGACCGAACAGTTGACCGTTGAA
CAGGATATTGAAGAATTTACCTTCCTGCTGATTAATGGTGTTTGTCCGGGTACACAGTGTTGATGAAG
GTCCGAGACGCCCGTCAACGGAGAACGGCGAATGATTGAGAATACCTATAGCGAAAAGTTCGAGTCCG
CGTTCGAACAGATCAAAGCGGCGGCCAACGTGGATGCCGCCATCCGTATTCTCCAGGCGGAATATAAC
CTCGATTTCGTCACCTACCATCTCGCCCAGACAATCGCGAGCAAGATCGATTCGCCCTTCGTGCGCAC
CACCTATCCGGATGCCTGGGTTTCCCGTTACCTCCTCAACTGCTATGTGAAGGTCGATCCGATCATCA
AGCAGGGCTTCGAACGCCAGCTGCCCTTCGACTGGAGCGAGGTCGAACCGACGCCGGAGGCCTATGCC
ATGCTGGTCGACGCCCAGAAACACGGCATCGATGACAATGGCTACTCCATCCCCGTCGCCGACAAGGC
GCAGCGCCGCGCCCTGCTGTCGCTGAATGCCCATATACCGGCCGACGAATGGACCGAGCTCGTGCGCC
GCTGCCGCAATGAGTGGATCGAGATCGCCCATCTGATCCACCGCAAGGCCGTATATGAGCTGCATGGC
GAAAACGATCCGGTGCCGGCATTGTCGCCGCGCGAGATCGAGTGTCTGCACTGGACCGCCCTCGGCAA
GGATTACAAGGATATTTCGGTCATCCTGGGCATATCAGAGCATACCACACGCGATTACCTGAAAACCG
CCCGCTTCAGGCTCGGCTGCACCACGATCTCGGCCGCCGCGTCGCGGGCTGTTCAATTGCGCATCATC
AATCCCTATAGGATCCGCATGACGCGACGTAATTGGTAATGAGCTTAAACTAACGAACGTAAATAAGG
AGGATAGACATGGACATGCCTCGTATTAAACCGGGTCAGCGTGTTATGATGGCACTGCGTAAAATGAT
TGCAAGCGGTGAAATCAAAAGTGGTGAACGTATTGCAGAAATTCCGACCGCAGCAGCACTGGGTGTTA
GCCGTATGCCGGTTCGTATCGCACTGCGTTCACTGGAACAAGAAGGTCTGGTTGTTCGTCTGGGTGCA
CGTGGTTATGCAGCCCGTGGTGTTAGCAGCGATCAGATTCGTGATGCAATTGAAGTTCGTGGTGTTCT
GGAAGGTTTTGCAGCACGTCGTCTGGCAGAACGTGGTATGACCGCAGAAACCCATGCACGTTTTGTTG
TACTGATTGCAGAAGGTGAAGCACTGTTTGCAGCCGGTCGCCTGAATGGTGAAGATCTGGATCGTTAT
GCCGCATATAATCAGGCATTTCATGATACCCTGGTTAGCGCAGCAGGTAATGGTGCAGTTGAAAGCGC
ACTGGCACGTAATGGTTTTGAACCGTTTGCAGCAGCCGGTGCACTGGCCCTGGATCTGATGGACCTGT
CTGCCGAATATGAACATCTGCTGGCAGCACATCGTCAGCATCAGGCAGTTCTGGATGCAGTTAGCTGT
GGTGATGCCGAAGGTGCAGAACGTATTATGCGTGATCATGCACTGGCAGCAATTCGTAATGCAAAAGT
TTTTGAAGCAGCAGCAAGCGCAGGCGCACCGCTGGGTGCAGCATGGTCAATTCGTGCAGATTGATAAC
TCGGTACCAAATTCCAGAAAAGAGGCCTCCCGAAAGGGGGGCCTTTTTTCGTTTTGGTCCCGACGTAC
GGTGGAATCTGATTCGTTACCAATTGACATGATACGAAACGTACCGTATCGTTAAGGTAGCTGTCACC
GGATGTGCTTTCCGGTCTGATGAGTCCGTGAGGACGAAACAGCCTCTACAAATAATTTTGTTTAATCT
AGAGAAAGAGGGGAAATACTAGatgacagtaaagaagctttatttcgtcccagcaggtcgttgtatgt
tggatcattcgtctgttaatagtacattaacaccaggagaattattagacttaccggtttggtgttat
cttttggagactgaagaaggacctattttagtagatacaggtatgccagaaagtgcagttaataatga
aggtctttttaacggtacatttgtcgaagggcaggttttaccgaaaatgactgaagaagatagaatcg
tgaatattttaaaacgggttggttatgagccggaagaccttctttatattattagttctcacttgcat
tttgatcatgcaggaggaaatggcgcttttataaatacaccaatcattgtacagcgtgctgaatatga

ggcggcgcagcatagcgaagaatatttgaaagaatgtatattgccgaatttaaactacaaaatcattg
aaggtgattatgaagtcgtaccaggagttcaattattgcatacaccaggccatactccagggcatcaa
tcgctattaattgagacagaaaaatccggtcctgtattattaacgattgatgcatcgtatacgaaaga
gaattttgaaaatgaagtgccatttgcgggatttgattcagaattagctttatcttcaattaaacgtt
taaaagaagtggtgatgaaagagaagccgattgttttctttggacatgatatagagcaggaaaggga
tgtaaagtgttccctgaatatatatgataaATAGTAATTGGTAACGAATCAGACAATTGACGGCTCGA
GGGAGTAGCATAGGGTTTGCAGAATCCCTGCTTCGTCCATTTGACAGGCACATTATGCATCGATGATA
AGCTGTCAAACATGAGCAGATCCTCTACGCCGGACGCATCGTGGCCGGCATCACCGGCGCCACAGGTG
CGGTTGCTGGCGCCTATATCGCCGACATCACCGATGGGGAAGATCGGGCTCGCCACTTCGGGCTCATG
AGCAAATATTTTATCTGTGACCAAATTCTCAGTGGTTGTCGCAGGCGGTGGAAGCACCTTTACGCCAc
tgcagggcgtaatcatggtcatagctgtttcctgtgtgaaattgttatccgctcacaattccacacaa
catacgagccggaagcataaagtgtaaagcctggggtgcctaatgagtgagctaactcacattaattg
cgttgcgctcactgcccgctttccagtcgggaaacctgtcgtgccagctgcattaatgaatcggccaa
cgcgaaccccttgcggccgcccgggccgtcgaccaattctcatgtttgacagcttatcatcgaatttc
tgccattcatccgcttattatcacttattcaggcgtagcaaccaggcgtttaagggcaccaataactg
ccttaaaaaaattacgccccgccctgccactcatcgcagtactgttgtaattcattaagcattctgcc
gacatggaagccatcacaaacggcatgatgaacctgaatcgccagcggcatcagcaccttgtcgcctt
gcgtataatatttgcccatggtgaaaacgggggcgaagaagttgtccatattggccacgtttaaatca
aaactggtgaaactcacccagggattggctgagacgaaaaacatattctcaataaaccctttagggaa
ataggccaggttttcaccgtaacacgccacatcttgcgaatatatgtgtagaaactgccggaaatcgt
cgtggtattcactccagagcgatgaaaacgtttcagtttgctcatggaaaacggtgtaacaagggtga
acactatcccatatcaccagctcaccgtctttcattgccatacgaaattccggatgagcattcatcag
gcgggcaagaatgtgaataaaggccggataaaacttgtgcttatttttctttacggtctttaaaaagg
ccgtaatatccagctgaacggtctggttataggtacattgagcaactgactgaaatgcctcaaaatgt
tctttacgatgccattgggatatatcaacggtggtatatccagtgatttttttctccattttagcttc
cttagctcctgaaaatctcgataactcaaaaaatacgcccggtagtgatcttatttcattatggtgaa
agttggaacctcttacgtgccgatcaacgtctcattttcgccaaaagttggcccagggcttcccggta
tcaacagggacaccaggatttatttattctgcgaagtgatcttccgtcacaggtatttattcgcgata
agctcatggagcggcgtaaccgtcgcacaggaaggacagagaaagcgcggatctgggaagtgacggac
agaacggtcaggacctggattggggaggcggttgccgccgctgctgctgacggtgtgacgttctctgt
tccggtcacaccacatacgttccgccattcctatgcgatgcacatgctgtatgccggtataccgctga
aagttctgcaaagcctgatgggacataagtccatcagttcaacggaagtctacacgaaggtttttgcg
ctggatgtggctgcccggcaccgggtgcagtttgcgatgccggagtctgatgcggttgcgatgctgaa
acaattatcctgagaataaatgccttggcctttatatggaaatgtggaactgagtggatatgctgttt
ttgtctgttaaacagagaagctggctgttatccactgagaagcgaacgaaacagtcgggaaaatctcc
cattatcgtagagatccgcattattaatctcaggagcctgtgtagcgtttataggaagtagtgttctg
tcatgatgcctgcaagcggtaacgaaaacgatttgaatatgccttcaggaacaatagaaatcttcgtg
cggtgttacgttgaagtggagcggattatgtcagcaatggacagaacaacctaatgaacacagaacca
tgatgtggtctgtccttttacagccagtagtgctcgccgcagtcgagcgacagggcgaagccctcggc
tggttgccctcgccgctgggctggcggccgtctatggccctgcaaacgcgccagaaacgccgtcgaag
ccgtgtgcgagacaccgcggccggccgccggcgttgtggatacctcgcggaaaacttggccctcactg
acagatgagggcggacgttgacacttgaggggccgactcacccggcgcggcgttgacagatgagggg
caggctcgatttcggccggcgacgtggagctggccagcctcgcaaatcggcgaaaacgcctgattta
cgcgagtttcccacagatgatgtggacaagcctggggataagtgccctgcggtattgacacttgaggg
gcgcgactactgacagatgaggggcgcgatccttgacacttgaggggcagagtgctgacagatgaggg
gcgcacctattgacatttgaggggctgtccacaggcagaaaatccagcatttgcaaggtttccgccc

gtttttcggccaccgctaacctgtcttttaacctgcttttaaaccaatatttataaaccttgtttta
accagggctgcgccctgtgcgcgtgaccgcgcacgccgaagggggtgcccccccttctcgaaccctc
ccggtcgagtgagcgaggaagcaccagggaacagcacttatatattctgcttacacacgatgcctgaa
aaaacttcccttggggttatccacttatccacggggatatttttataattattttttttatagttttt
agatcttcttttttagagcgccttgtaggcctttatccatgctggttctagagaaggtgttgtgacaa
attgccctttcagtgtgacaaatcaccctcaaatgacagtcctgtctgtgacaaattgcccttaaccc
tgtgacaaattgccctcagaagaagctgtttttttcacaaagttatccctgcttattgactcttttttta
tttagtgtgacaatctaaaaacttgtcacacttcacatggatctgtcatggcggaaacagcggttatc
aatcacaagaaacgtaaaaatagcccgcgaatcgtccagtcaaacgacctcactgaggcggcatatag
tctctcccgggatcaaaaacgtatgctgtatctgttcgttgaccagatcagaaaatctgatggcaccc
tacaggaacatgacggtatctgcgagatccatgttgctaaatatgctgaaatattcggattgacctct
gcggaagccagtaaggatatacggcaggcattgaagagtttcgcggggaaggaagtggtttttttatcg
ccctgaagaggatgccggcgatgaaaaaggctatgaatcttttccttggtttatcaaacgtgcgcaca
gtccatccagagggctttacagtgtacatatcaacccatatctcattcccttctttatcgggttacag
aaccggtttacgcagtttcggcttagtgaaacaaaagaaatcaccaatccgtatgccatgcgtttata
cgaatccctgtgtcagtatcgtaagccggatggctcaggcatcgtctctctgaaaatcgactggatca
tagagcgttaccagctgcctcaaagttaccagcgtatgcctgacttccgccgccgcttcctgcaggtc
tgtgttaatgagatcaacagcagaactccaatgcgcctctcatacattgagaaaaagaaaggccgcca
gacgactcatatcgtattttccttccgcgatatcacttccatgacgacaggatagtctgagggttatc
tgtcacagatttgagggtggttcgtcacatttgttctgacctactgagggtaatttgtcacagttttg
ctgtttccttcagcctgcatggattttctcatactttttgaactgtaatttttaaggaagccaaattt
gagggcagtttgtcacagttgatttccttctctttcccttcgtcatgtgacctgatatcgggggttag
ttcgtcatcattgatgagggttgattatcacagtttattactctgaattggctatccgcgtgtgtacc
tctacctggagtttttcccacggtggatatttcttcttgcgctgagcgtaagagctatctgacagaac
agttcttctttgcttcctcgccagttcgctcgctatgctcggttacacggctgcggcggatgtgctgc
aaggcgattaagttgggtaacgccagggttttcccagtcacgacgttgtaaaacgacggccagtgcgg
ccgcTAACCAATCAGGCTTCCTACTTACAGAATTGAGAAAAGAGGATGTGGAA
